# Modular Scaffold Crystals for Programmable Installation and Structural Observation of DNA-Binding Proteins

**DOI:** 10.64898/2026.03.04.709581

**Authors:** Ethan T. Shields, Caroline K. Slaughter, Fadwa Mekkaoui, Emma N. Magna, Cole Shepherd, Philip S. Lukeman, Donald E. Spratt, Christopher D. Snow

## Abstract

Inducing biomacromolecules to self-assemble into diffraction-quality crystals remains a major challenge, typically overcome by brute-force experimental screening. Inspired by Seeman’s vision of DNA-junction-based scaffolds organizing guest biomacromolecules, we developed a protein–DNA co-crystal combining modular DNA programmability with robust protein-lattice diffraction. Our engineered co-crystals are composed of stacked double-stranded DNA scaffolded by protein columns, surrounding solvent channels designed to enable guest protein diffusion. DNA ‘strut’ variation allows positionally-controlled installation of diverse DNA binding guest proteins. Experimentally, we simply grow scaffold crystals under standardized conditions, ligate the scaffold, and soak guest proteins. Decoupling crystal growth from guest installation will enable high-throughput structure determination of diverse DNA-binding proteins <u>and protein-macromolecule conjugates. Sub-nanometer position and orientation control of guest macromolecules will also enable functional applications beyond structural biology.</u>

## Introduction

Engineered biomolecular crystals built from proteins and/or DNA are highly precise, tunable, and modular materials that have been used to host guest molecules for structure determination (1–5), catalyze reactions (6–9), store or deliver cargo (10–14), and function as molecular sieves or other nanoscale devices (15–17). Maximally useful scaffold crystals will embody several key properties. *Modular* crystals permit routine site-specific changes. *Porous* crystals allow guest macromolecules to diffuse through the crystal interior. *Programmable* crystals offer site-specific guest capture/installation. *Precise* crystals provide high-resolution X-ray diffraction (XRD). DNA crystals are especially attractive for their predictable assembly (18–25), low cost, and sequence flexibility, but they rarely achieve the desirable but challenging combination of large pore size and high-resolution diffraction (26,27). In contrast, protein crystals offer diverse chemical environments, and in some cases form highly porous lattices that nonetheless diffract to atomic resolution (28–31). However, protein crystal growth can be very sensitive to point mutations (32,33) and engineering protein-protein interfaces is intrinsically more challenging than designing paired DNA sticky overhangs. Thus, by our definitions above, protein crystals are less modular and programmable.

To combine the modularity of DNA assembly with the precision of protein lattices, we developed a new class of porous protein–DNA co-crystals (henceforth co-crystal 1, CC1). Our designed lattice is stabilized in two directions by coaxial DNA:DNA interfaces (fine-tunable blunt and sticky ended cohesion). Each dsDNA is also held by a scaffold protein, and the third lattice direction is stabilized by a protein stack. We previously reported an interpenetrating version of these crystals (ipCC1) capable of incorporating varied sequences and capturing small molecules, but with limited pore size (34–36). We now report variants with solvent channels large enough for guest protein diffusion. We demonstrate installation of six different DNA-binding domains (DBDs) with varied binding modes that bind with sufficient specificity and affinity to be resolved by XRD. The resulting family of modular scaffold crystals constitutes a versatile new tool for structural studies of diverse DBDs, instantiating the vision first articulated by Seeman in 1982 of a crystalline framework that reliably hosts guest proteins at specific sites for XRD observation (37).

The division of crystal preparation into two distinct phases—scaffold growth and guest installation—is impactful in multiple ways. First, host crystals can be engineered independent of the target guest (e.g. improving diffraction resolution or preserving diffraction in a wide range of solvent conditions). Second, modular host crystals can be reliably grown under standard conditions, eliminating the traditional crystallography bottleneck. Third, guests are added simply by soaking, and we can use mass action to observe weaker interactions that are recalcitrant to traditional methods. Fourth, the resulting 3D molecular pegboards could ultimately offer routes for the detailed control of the position and orientation of functional guest macromolecules (nanobiotechnology) as well as the structure determination of targets beyond DBD by providing multiple anchor sites to constrain the orientation of suitably tagged guest macromolecules.

## Results

### Modular Crystal Construction and Scaffold Expansion

The parent co-crystal scaffold (CC1) consists of the replication initiator protein RepE54 and a 21-base pair DNA duplex containing the 19-base pair cognate binding sequence (38). The lattice is stabilized by one symmetry-distinct coaxial DNA–DNA interface and two protein–protein interfaces (Fig. S1). This geometry enables two-dimensional expansion of the lattice by inserting dsDNA struts at the interface (“telescoping” the DNA), while preserving the protein-protein contacts (Fig. 1). Critically, we successfully grew expanded crystals with no observed limit to inserted DNA sequence diversity (Table S1).

**Figure 1.**
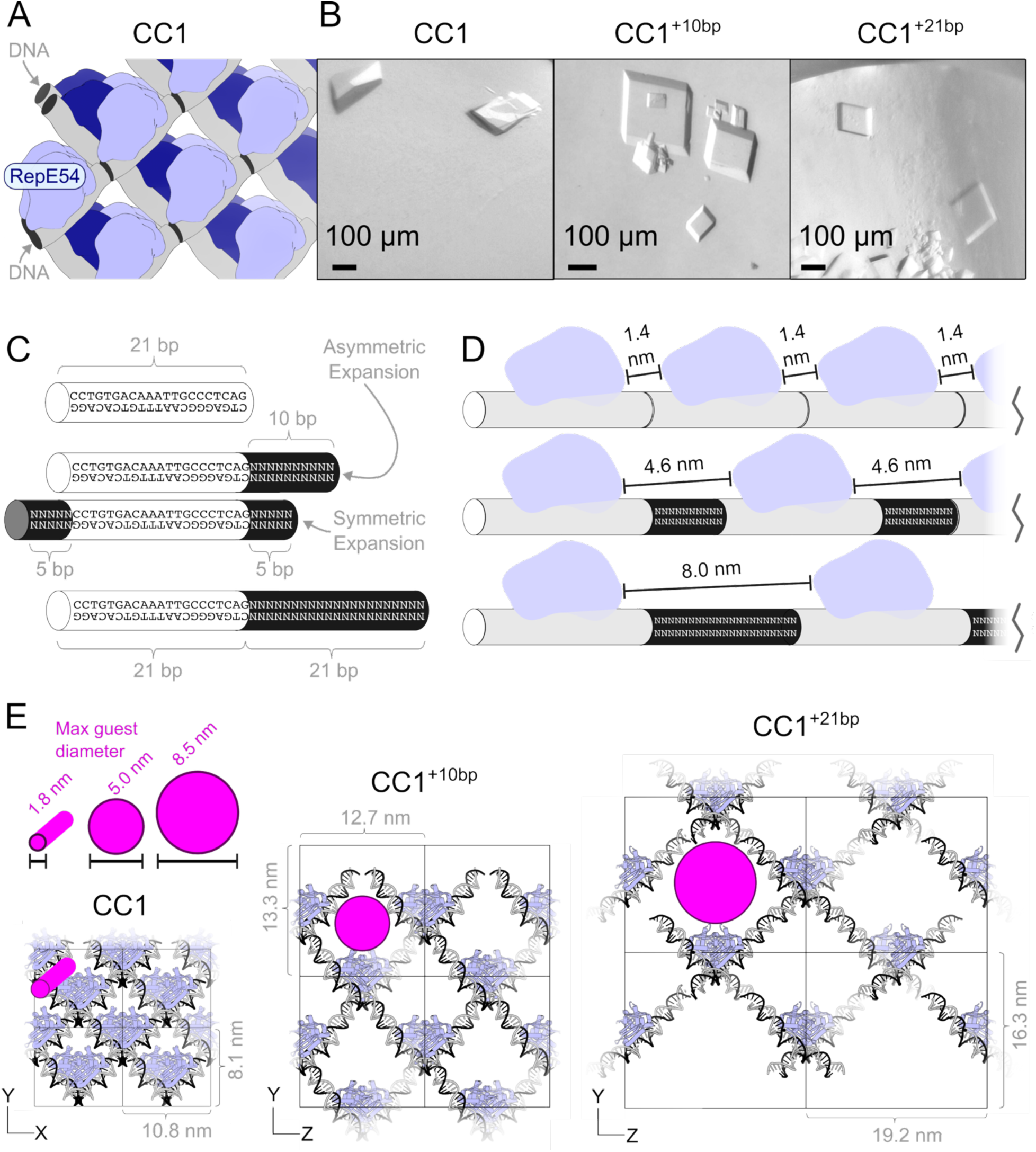
Co-crystal isoreticular expansion. **(A)** CC1 crystals grow via coaxial DNA:DNA interfaces (gray) in two directions and a protein:protein stack (blue shades) in the last direction. **(B)** Light microscope images of representative CC1 family crystals with characteristic parallelepiped habit. 100 micron scale bars. **(C)** CC1 crystals are expanded by adding new dsDNA (black cylinders) to the parent sequence (white cylinders) that encodes RepE54 binding. **(D)** Expansion along the DNA increases the minimum distance between RepE54 protein (light blue) symmetry mates. **(E)** The CC1 lattice family is shown with magenta discs depicting the maximum diffusible object diameter (see Fig. S4 for calculation details).

In prior work (35), adding 10 base pairs (i.e. CC1^+10^, 31 bp total) yielded crystals in space group I222 (PDB 7U6K) consisting of two interpenetrating lattices rotated 180°, a phenomenon also observed in porous MOFs (39) (Fig. S2). To favor non-interpenetrating growth, we reduced the Mg(II) concentration (Fig. S3). Underscoring assembly robustness, 15 crystal variants grew (Fig. S2C, Table S2) by varying only magnesium acetate (10-80 mM) and lithium sulfate (0.3-1.8 M). The porous CC1^+10^ lattices had ∼82% solvent content and solvent channels wide enough (5 nm diameter) to accommodate guest proteins (Fig. S4).

The porous CC1^+10^ lattice retained the high modularity of the prior, interpenetrating scaffold crystals. Crystals formed rapidly (commonly 1-7 days) using both symmetric (5 bp flanking each side) and asymmetric (10 bp on one side) expansions (Fig. 1C). Despite substantial differences in DNA sequence, sticky ends, terminal phosphorylation, and expansion strategy (Table S1), 15 variants formed diffraction-quality porous crystals with similar unit cell dimensions. Two variants have not yet grown (Table S3).

To further probe lattice tunability, we extended the inserted DNA to 21 base pairs (CC1^+21^, 42 bp total). These crystals grew with the same characteristic habit (Fig. 1B), but crystal growth was less frequently successful (Tables S4 and S5). Expansion led to significantly larger channels (8.5 nm diameter) and reduced empty-crystal resolution to 5.1Å. In both cases (Fig. 1E), expansion was isoreticular, i.e. with no substantive change to the molecular interfaces or lattice (space group no. 5, in either the C121 or I121 setting, Fig. S5). Despite reduced crystal yield for CC1^+21^, successful growth of CC1, CC1^+10^, and CC1^+21^ demonstrated robust modularity across increasingly porous designs.

### Asynchronous Loading of Guest Proteins

To evaluate CC1^+10^ as a general scaffold for structural studies, we engineered the DNA struts to include binding sites for model guest proteins drawn from three structurally diverse DBD families—homeodomains, coiled-coils, and zinc-fingers—selected for their fundamental biological importance and diverse binding modality. From the homeodomain family, we selected four *Drosophila melanogaster* variants: Even-skipped (EVE-HD), Ultrabithorax (UBX-HD), Antennapedia (ANTP-HD), and Engrailed homeodomain fused to enhanced green fluorescent protein (EnH-eGFP). From the coiled-coil family, we designed bZip, an engineered variant of the DBD of the *Saccharomyces cerevisiae* GCN4 transcription factor (Fig. S6). From the zinc-finger family, we designed a 33-a.a. peptide (C-clamp) corresponding to the C-clamp domain of human HDBP1 (Fig. S7). For each candidate guest binding register, CC1^+10^ crystal symmetry was applied to AlphaFold3 models (40)(Fig. S8) to predict steric feasibility in the lattice.

During the pursuit of guest binding within the CC1^+10^ lattice, fluorophore-labeled oligos (Fig. S9) and fluorescence polarization (FP) assays were used to scout the effect of several ionic conditions on homeodomain binding affinity for CC1^+10^ target sequences (Fig. S10). Buffers that maximized homeodomain binding had much lower salt concentrations CC1^+10^ lattice growth buffers (Fig. S2). Fortunately, ligated CC1^+10^ crystals (34,36) make it easy to soak in new guest molecules *and* use new solution conditions. Lower salt guest loaded crystals (50 mM KCl, 4 mM CaCl_2_) diffracted similarly to as-grown CC1^+10^ crystals with high Li_2_SO_4_ (Table S6). We then demonstrated that a narrow set of solution conditions was sufficient to resolve multiple diverse DBDs (Table S7).

To monitor guest loading in real time, UBX-HD was trace-labeled with N-hydroxysuccinimide fluorescein (41) and incubated with ligated CC1^+10^ crystals. Confocal fluorescence microscopy revealed uniform guest loading (30,31,42,43) throughout the lattice after one day (Fig. 2). Time-resolved z-stacks (Fig. 2B) revealed that the top layers farthest from the microscope slide adsorbed fluorescent guest (UBX-HD) most rapidly, consistent with the major solvent pores (5 nm diameter) being perpendicular to the slide. However, guests as small as UBX-HD might also diffuse laterally within the crystals (Fig. S11). For biophysicists, the quantitative study of intra-crystal guest transport via confocal microscopy should provide new opportunities to study the effect of DNA sequence and functional groups on the diffusion and binding kinetics of fluorescent guests in an extraordinarily precise 3D environment (31,43).

**Fig. 2.**
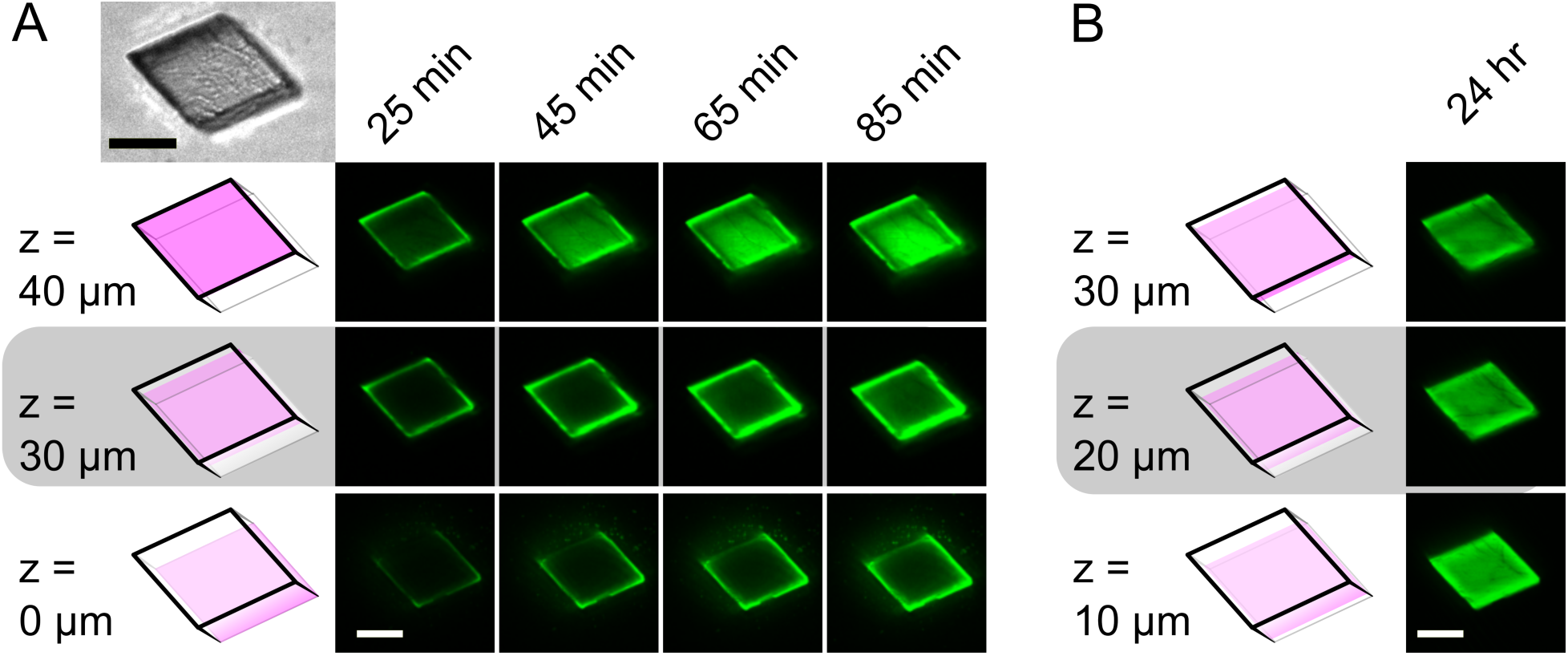
Confocal microscopy of a CC1^+10^ crystal adsorbing UBX-HD. **(A)** Fluorescein-trace-labeled UBX-HD (23 μM) was diluted 1:200 with unlabeled UBX-HD and used to soak a CC1^+10^ crystal containing an asymmetric 11-nt insert 5’<u>TAATTA</u>GGCCG 3’. We then obtained a time-resolved, 1 μm z-stack. Planes (pink in the schematic) near the microscope slide (z = 0 μm), near the crystal top (z = 40 μm), or interior (z = 30 μm) were selected to show the anisotropic guest loading pattern. A differential interference contrast image (inset top left) is aligned to the z-plane schematics for comparison. **(B)** After 24 hr, the laser intensity was reduced from 20% to 12% to avoid over-saturation. Loading was comparable for 3 internal planes evenly spaced within the crystal interior. For confocal microscopy, dilute trace labeling was helpful to avoid extinction effect complications (Fig. S12-S14). All scale bars are 50 μm.

We successfully soaked six guest proteins into ligated CC1^+10^ crystals and observed clear electron density for each bound DBD by X-ray diffraction. EVE-HD, UBX-HD, ANTP-HD, bZip, and C-clamp *were all visualized in expected binding modes* (Fig. 3). EnH–eGFP showed ordered density for the homeodomain but not for the flexibly tethered eGFP domain. The XRD resolutions of the resulting structures (Table S8) ranged from 3.0Å (EVE-HD and bZip) to 7.2Å (C-clamp). Structures for the model guest proteins were consistent with known conventional crystal structures (Tables S9 and S10), despite featuring DBD:DNA pairings new to the Protein Data Bank. To verify that the observed electron density was unbiased, we fully refined structures without modeling the guests (44)(Fig. 3AD) and calculated omit maps for each guest-bound structure (Fig. S15) (45–47). In all cases, strong electron density features (both 2F_0_-F_C_ and F_0_-F_C_) were present at the expected DNA-binding sites prior to model building, confirming that guest placement in the refined models reflects genuine experimental signal. Surprisingly, while EVE-HD did bind the expected target sequence register, a “bonus copy” of EVE-HD bound even more clearly to an adventitious site opposite RepE54 (see Discussion). This site consists of base pairs 10-15 (henceforth Register 1. R1:10–15).

**Fig. 3.**
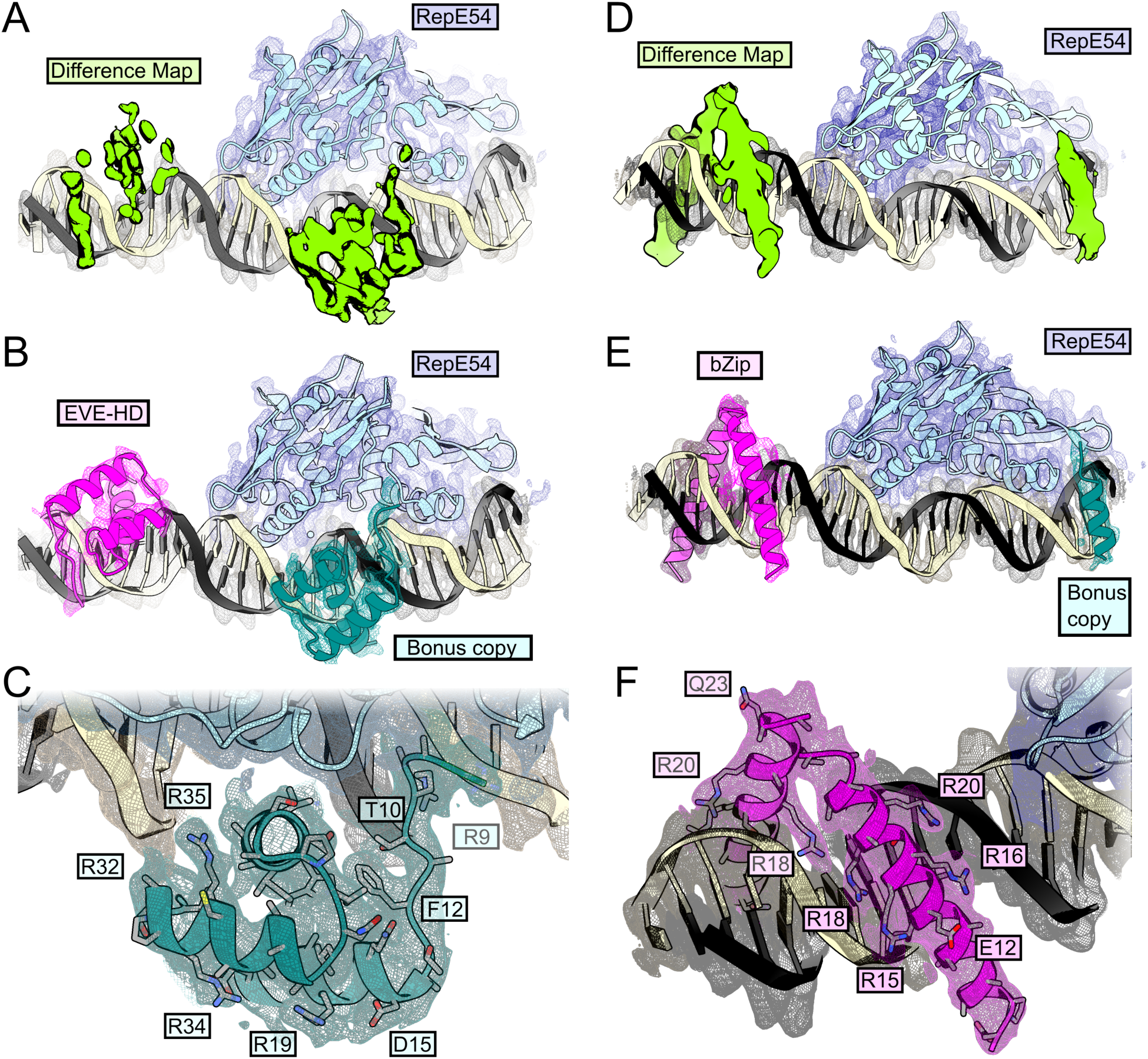
Guest structure determination of EVE-HD and bZip. **(A)** Unbiased F_0_-F_C_ difference map for EVE-HD from PDB 9YZM contoured at 2σ (green). **(B)** Corresponding crystal structure of PDB 9YZM with 2F_0_-F_C_ map contoured at 1σ, Eve-HD bound to canonical sequence 5’ TAATTA 3’ (magenta) and bonus copy of EVE-HD bound to R1 site (teal). **(C)** Close view of EVE-HD bound to R1 register with selected residues shown as sticks and 2F_0_-F_C_ map contoured at 0.5σ (teal). **(D)** Unbiased F_0_-F_C_ difference map for bZip from PDB 9YZU contoured at 2σ (green). **(E)** Corresponding crystal structure of PDB 9YZU with 2F_0_-F_C_ map contoured at 1σ, bZip bound to canonical sequence 5’ ATGAGTCAT 3’ (magenta) and bonus partial structure of bZip (teal). (**F**) Close view of bZip bound to its canonical target sequence with selected residues shown as sticks and 2F_0_-F_C_ map contoured at 0.5σ (magenta).

Binding to multiple DNA sites was not unique to EVE-HD. All homeodomains tested displayed similar behavior, and in the case of UBX-HD, a second adventitious site (R2:18-23, 5’-TCATAA-3’) was observed in addition to the first noncanonical site (R1: 10–15, 5’-AATTGC-3′). Isothermal calorimetry (ITC) experiments (Fig. S16) showed that EVE-HD and UBX-HD bind the R1 and R2 sequences with relatively weak affinity in solution (0.1-9 μM).

Together, these results demonstrate that CC1^+10^ can capture and resolve a structurally and phylogenetically diverse set of DBDs. Asynchronous installation of guest proteins via soaking provides a new route for high-throughput structural characterization of DNA–protein interactions, circumventing the trial-and-error aspects of traditional crystallography. No fundamental barrier prevents scale-up to obtain DBD structures in hundreds of solution conditions, nor with numerous DNA sequence variants. We anticipate high-throughput structure determination will aid future machine learning models for protein-DNA recognition. Combining scaffold crystal technology with room-temperature XRD may further increase throughput (Fig. S17)

### Operating a Crystalline Molecular Goniometer

To further probe the versatility of CC1^+10^ for asynchronous guest loading, we systematically shifted the binding register/sequence of EVE-HD and bZip along the DNA strut. Due to the helical DNA twist, register shifts are concomitant with rotation, echoing the goniometer analogy established by Douglas (48). Five CC1^+10^ variants were designed, three with shifting EVE-HD capture sites and two with shifting bZip sites (Fig. S18). EVE-HD or bZip was successfully installed into the corresponding ligated crystals. Clear electron density allowed guest structure determination in each case, despite the use of a palindromic sequence (TAATTA) for two EVE-HD capture sites (Fig. 4).

**Fig. 4.**
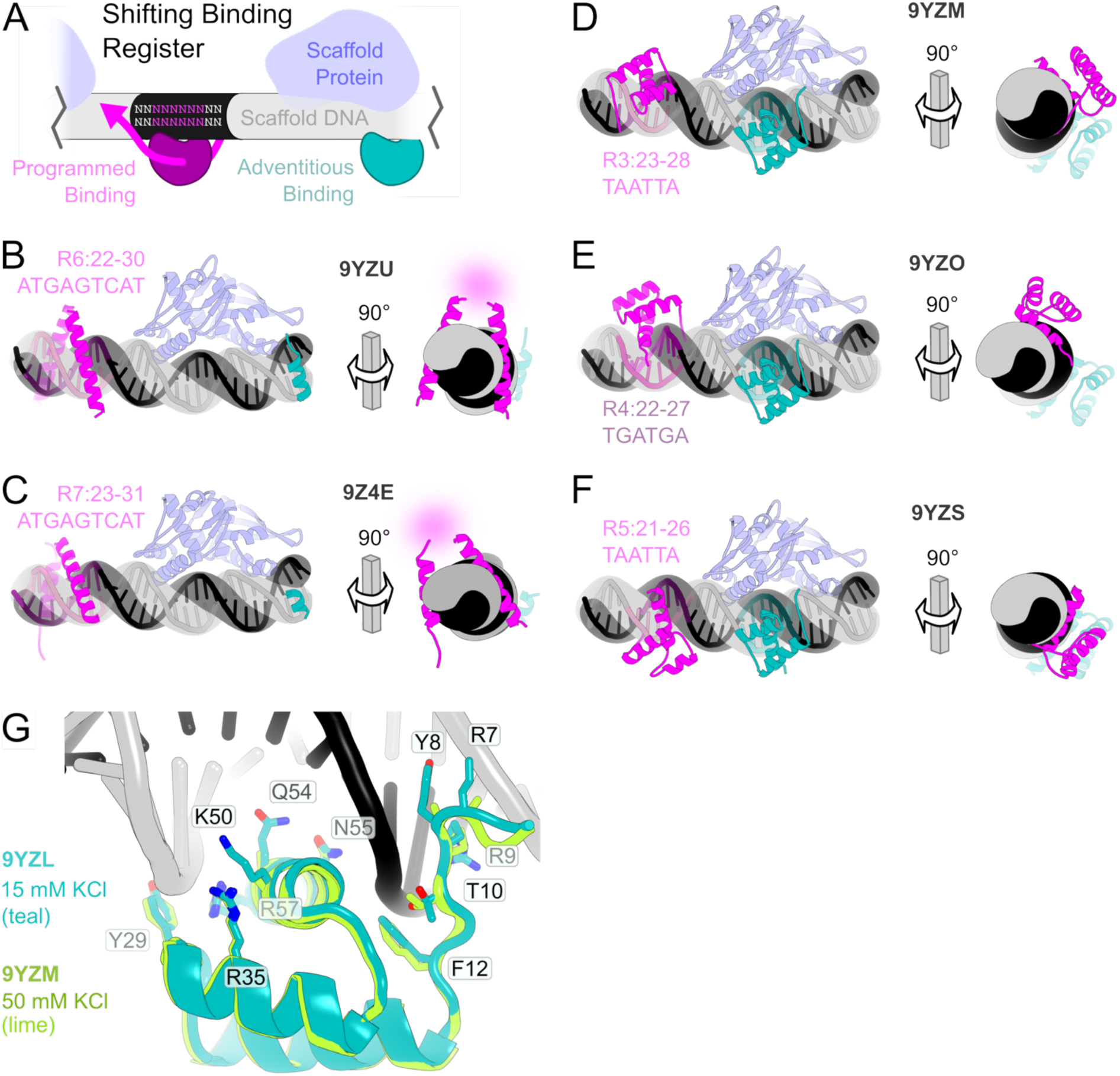
Guest structures with shifted and varied binding sites. **(A)** Schematic representation of CC1^+10^ with shifting programmed binding sites (magenta) and adventitious binding (teal). Each crystal structure below is likewise shown with guest proteins bound to target locations (magenta) and bonus sites (teal). Each model is also shown with a 90° rotated view. **(B)** CC1^+10^ with asymmetric insert 5’<u>ATGAGTCAT</u>A 3’ loaded with bZip (PDB 9YZU). **(C)** CC1^+10^ with asymmetric insert 5’G<u>ATGAGTCAT</u> 3’ loaded with bZip (PDB 9Z4E), with unresolved putative C-terminal disulfide (magenta blur). **(D)** CC1^+10^ with asymmetric insert 5’-GC<u>TAATTA</u>GGC-3’ loaded with EVE-HD (PDB 9YZM). **(E)** CC1^+10^ with asymmetric insert 5’-T<u>TGATGA</u>GCAG-3’ loaded with EVE-HD (PDB 9YZO). **(F)** CC1^+10^ with asymmetric insert 5’-<u>TAATTA</u>GGCCG-3’ loaded with EVE-HD (PDB 9YZS). **(G)** Comparison of Eve-HD bound to R1 site under two different monovalent cation concentrations (PDBs 9YZL and 9YZM) with a subset of the resolved residues shown with the stick representation.

## Discussion

### CC1 vs Other Crystal Scaffold Systems

The CC1 family (Table S8) achieves a combination of properties that have proven exceptionally difficult to achieve in other highly porous engineered macromolecular crystal systems – modular assembly, significant freedom in the selection of guest installation sites, and relatively high diffraction resolution (∼3Å). To date, highly porous crystals (pore diameter > 5 nm) composed solely of DNA have rarely provided high-resolution (<4Å) diffraction (Fig. S19)(15,49). We attribute the improved CC1^+10^ diffraction to the elimination of Holliday junctions and the “insulating” effect of protein components that rigidify and separate adjacent DNA duplexes. Further resolution improvements might be achieved by reducing defects, enhancing scaffold rigidity (50), optimizing crystallographic interfaces (51), or by combining multiple datasets with advanced analysis strategies (52–54) to combat flexibility of the exposed DNA struts.

Our scaffold-assisted structure determination approach favorably complements other structural biology scaffold strategies developed to date. Proteins can be encapsulated within MOFs, but controlling guest orientation and large single crystal growth remain challenging (55–57). Crystals composed entirely of DNA origami assemblies have enabled remarkable *in-crystallo* control of guests such as gold nanoparticles (58,59). As with DNA-directed colloidal crystals (22–25), where bridging DNA controls the assembly of nanoparticles or proteins, DNA origami crystals have been more suitable for electron microscopy and small angle X-ray scattering than high-resolution XRD. Scaffold-engineering for cryo-EM (48,60–64) is a partially-related research direction that is complementary to our approach, since scaffold crystals intrinsically favor guest macromolecules small enough for intra-crystal diffusion whereas cryo-EM systems favor larger targets. Likewise, porous crystals composed entirely of protein can support guest soaking and site-specific installation (30,65) but typically lack the facile modular programmed assembly and guest DBD installation capacity of nucleic acid components. In contrast, the CC1 family combines the design flexibility of DNA scaffolds with the diffraction quality and chemical robustness of protein crystals, providing a platform that is simultaneously modular, programmable, porous, and structurally precise. As a result, CC1^+10^ offers the first plug-and-play installation of guest macromolecules within scaffold crystals.

The ability to rapidly obtain structures under varying conditions, thanks to ligation-stabilized host crystals, is another notable strength of the scaffold-assisted crystallography method. Subtle differences in side-chain electron density were observed for EVE-HD when we varied the ionic conditions (Fig. 4G). Future efforts can map the range of buffer conditions simultaneously compatible with guest binding and high-resolution XRD and obtain high-throughput guest structures under hundreds of solution conditions.

### Tunable Assembly

The CC1 platform offers easy tuning of lattice assembly through oligonucleotide design. Because crystal growth in two dimensions is mediated by coaxial DNA–DNA contacts, assembly can be modulated by varying the DNA building blocks. As described previously (35), the total insert length (e.g., 10 bp) was chosen to allow each building block to form Watson–Crick overhangs with its neighbors with minimal geometric strain. Here, we further validate this principle by demonstrating successful growth of CC1^+10^ lattices with a range of overhang lengths (0, 1, and 2 nucleotides) and base compositions (AT-rich and GC-rich) (Table S1). In one instance (PDB 9Z08), the DNA insert was a separate molecule. The successful growth of crystals with 0-nt blunt-end interfaces underscores the dominant role of base-stacking in lattice formation (Fig. S20) (66).

### Size Constraints

A defining feature of highly porous biomolecular crystals is the capacity for intra-crystal guest macromolecule diffusion. Analysis with MAP_CHANNELS (67) revealed that the maximum guest sphere diameter capable of diffusing through the CC1^+10^ lattice is expected to be ∼5 nm (Fig. S4 and Table S8). Many globular proteins fall below this threshold; for example, eGFP approximates a 3 nm × 5 nm cylinder (68). Indeed, by revealing bound EnH, XRD data (PDB 9YZQ) confirmed that eGFP fused to the Engrailed homeodomain (EnH-eGFP) via a flexible 7 a.a. linker (Table S11) could access the crystal interior. Guests larger than 5 nm may be more suitable for the CC1^+21^ host crystals.

### Scaffold Binding Driving Forces

We conclude with a discussion of the driving forces needed to induce guest molecules to adopt a coherent structure within the host lattice. Despite covalent tethering to EnH, eGFP was not visible in the electron density. This result is consistent with previous observations that tethering guests via flexible linkers is not necessarily sufficient to induce high occupancy of a unique guest conformation within porous crystals; Maita (2) genetically fused ubiquitin to the building blocks of porous R1EN crystals and observed no density for a flexible 11-a.a. linker, but was able to model 50% occupancy ubiquitin binding sites for shorter linker lengths (3–7 aa).

To serve as a platform technology for the study of protein-DNA interactions, host crystals should be able to resolve both expected and unexpected complexes; this capacity was quite evident. Within CC1^+10^, homeodomains had a striking tendency to bind noncanonical sequences, especially the R1 site. This behavior was observed across all tested homeodomains and was not limited to a single site.

Since the R1 register DNA sequence contains 5’-AATT-3’, which partially matches the canonical 5’-TAATTA-3’ recognition motif, we decided to evaluate in-solution binding. ITC measurements (Fig. S18) found that the binding affinity for the R1 and R2 sequences (embedded in longer oligos) was significantly weaker than the low nanomolar affinity generally expected for homeodomains with cognate DNA (69). However, UBX-HD binding to the unexpected register R2:18-23 (5’TCATAA’3) was confirmed by ITC as a moderately strong (147±53 nM) recognition sequence in solution. The relative weakness of the R1 and R2 sites in solution supports a role for factors beyond DNA sequence readout in driving guest binding *in crystallo*.

Protein-DNA recognition is widely understood to involve both base readout and shape readout (70,71). In this context, the RepE54 component of the CC1 scaffold provides a rich environment by bending the scaffold DNA and compressing the minor groove within its binding site (Fig. S21). We hypothesize that this pre-organized compressed minor groove forms a thermodynamically favorable binding site for the positively charged N-terminal arm of the homeodomains (72). Lattice contacts may also play a role. For example, positioning EVE-HD at the R1 site enables favorable interactions with symmetry mates (Fig. S22). Notably, by varying the target installation site (Fig. S23) it is possible to either avoid or embrace lattice contacts.

The cost of inducing bending in the *strut* region due to guest binding might also contribute to binding preferences. We observed changes to the exposed DNA strut upon guest binding (Fig. S24), consistent with DNA flexibility as a contributing factor for register selection. We do not expect guests that require pre-existing extreme DNA curvature to bind the CC1^+10^ lattice, but induced-fit or bind-then-bend DBDs (73) might be observed at an early stage of biomolecular recognition.

Importantly, the ability of CC1 scaffolds to withstand high local guest concentrations (3Å diffraction after soaking in 30 μM guest protein) implies that even relatively weak interactions might be amenable to structure determination. Via mass action, otherwise invisible protein–DNA complexes can be stabilized and visualized, thereby providing access to a broader landscape of binding modes than is typically observable in solution. Therefore, scaffold-assisted crystallography may ultimately be useful to provide information for guest DBDs that lack strong sequence specificity.

Our collective findings suggest that binding register selection within the CC1 scaffold is driven by a combination of sequence recognition, shape recognition, and the lattice environment. Reliance on adventitious interactions to drive guests to adopt a uniform conformation has been a hallmark of prior successes (74), such as the crystalline sponge method for small molecules (55) and Maita’s tethered Ubiquitin (2). We also show the capacity to benefit from fortuitous binding.

#### Critically, fortuitous binding does not interfere with programmed binding

Reliable control and XRD observation of installed guest macromolecule position and orientation is a milestone achievement in programmed self-assembly, with future applications (Fig. S25) that extend beyond the structural biology of DNA-binding proteins.

## Acknowledgements

We thank Hataichanok (Mam) Scherman, Director of the Histone Source at Colorado State University for the expression and purification of RepE54 transcription factor; Jay Nix at ALS beamlines 4.2.2, 8.2.1, and 8.2.2 for support of XRD data collection; Kay Perry at APS beamline 24-IDE for support of XRD data collection. We thank the Massachusetts Life Sciences Center for their support in collected ITC data. We thank Abigail Orun for her early leadership on the project starting in 2017 and dedicate the work to the memory and scientific legacy of Nadrian C Seeman.

## Author Contributions

Conceptualization: CDS, ETS

Methodology: ETS, CKS, FM, ENM, CS

Investigation: ETS, FM

Visualization: ETS, CDS

Funding acquisition: PSL, CDS

Project administration: CDS

Supervision: CDS, DES

Writing – original draft: ETS, CDS

Writing – review & editing: ETS, CKS, FM, DES, PSL, CDS

## Data Availability

Key data are available in the main text, the supplementary materials, or the Protein Data Bank. Raw diffraction data is available (e.g. https://proteindiffraction.org/search/?q=SnowShieldsCC1). Fluorescence polarization data and fitting scripts are available at https://github.com/cdsnow/FPfitting. Isothermal calorimetry data and fitting scripts are available at https://github.com/cdsnow/ITCfitting. Code to model guest DBD within the scaffold crystals (register scanning) is available for use at https://www.engr.colostate.edu/∼cdasnow/guestscan.html or download at https://github.com/cdsnow/DBDscan. Confocal microscope data is available in a Zenodo repository: https://zenodo.org/records/18328922.

