## supplementary materials for "Modular Scaffold Crystals for Programmable Installation and Structural Observation of DNA-Binding Proteins"

**The PDF file includes:**

Materials and Methods
Figs. S1 to S27
Tables S1 to S8
References 75 to 119

### Materials and Methods

#### ***Protein Expression and Purification***

*Replication Initiator Protein RepE54*

Cloning and expression of the protein Replication initiator RepE54 was previously described by Komori *et al.* (38). Briefly, we cloned RepE54 into pSB3 vector (Addgene plasmid #82027) with an N-terminal His-tag. The CC1 RepE54 was expressed and purified by the Histone Source at Colorado State University as described in our previous ligation study (34). In short, the N-terminally hexahistagged protein from PDB code 7RVA was overexpressed in *E.coli* CodonPlus RIPL competent cells (Agilent #230280). Sonicated cell lysate was purified with Ni Excel Sepharose (Cytiva #17371202) and HiLoad Superdex 200 PG column (Cytiva). The resulting CC1 protein was concentrated to 15 mg/mL in storage buffer (100 mM sodium citrate pH 6.2, 100 mM KCl, 10 mM $MgCa_2$, and 10% glycerol). Aliquots were flash frozen with liquid nitrogen, and stored at -80°C. This variant of the RepE54 protein was used for all crystal structures other than 9YZA (which used the triple point mutant described immediately below).

*Replication Initiator Protein RepE54 Variant (L53G, Q54G, E55G)*

During early attempts to disrupt the interpenetrating habit, we noticed that the surface loop from residues 49-56 closely approached the same loop from a symmetry mate. In hopes of disrupting any favorable intermolecular contacts at that site we removed three sidechains. Variant RepE54 (L53G, Q54G, E55G) was cloned into the pETDuet plasmid (Novagen, Sigma-Aldrich # 71146-3) using Gibson cloning (Hi-Fi assembly New England Biolabs #E2621L). The protein was overexpressed with a T7 promoter in *E. coli* BL21(DE3) competent cells (New England Biolabs #C2527H). Upon addition of 0.5 mM IPTG, the cells were incubated at 25 °C for 20 h. The cell pellets were sonicated in lysis buffer (1X PBS, 300 mM NaCl, 25 mM imidazole, pH 7.4) and applied to HisTrap (HisPur$^{TM}$ Ni-NTA resin, ThermoFisher #88222) equilibrated with HisTrap buffer (1X PBS, 300 mM NaCl, 25 mM imidazole, pH 7.4). The protein was eluted with 100 mM imidazole in HisTrap buffer. Fractions containing the desired protein were pooled and further purified at the CSU Histone Source facility with HiLoad Superdex 200 PG column (Cytiva

#28989335). RepE54 variants (L53G, Q54G, E55G) were concentrated to 15 mg/mL in CC1 storage buffer (100 mM sodium citrate pH 6.2, 100 mM KCl, 10 mM $MgCa_2$, and 10% glycerol), flash frozen in liquid nitrogen, and stored at −80°C. Since this triple mutant was not noticeably better than the original construct at avoiding the interpenetrating habit, we devoted little additional effort to this variant, though we did crystallize it (PDB: 9YZA) in the new conditions that favor the porous lattice (the loop with the mutations was not resolved).

*Engrailed Homeodomain-eGFP (EnH-eGFP) Fusion*

The guest protein, Engrailed Homeodomain-eGFP Fusion (EnH-eGFP), was cloned into the pETDuet plasmid (Novagen, Sigma-Aldrich # 71146-3) using Gibson assembly (Hi-Fi assembly New England Biolabs #E2621L). EnH-eGFP protein was overexpressed with a T7 promoter in *E. coli* BL21(DE3) competent cells (New England Biolabs #C2527H). Upon addition of 0.5 mM IPTG, the cells were incubated at 25°C for 20 h. The cell pellets harvested by centrifugation, then sonicated in lysis buffer (100 mM Tris-HCl, 200 mM NaCl, 10% glycerol, 10 mM imidazole, pH 8.0) and loaded on to HisTrap (HisPur™ Ni-NTA Resin, ThermoFisher #88222) equilibrated with HisTrap buffer (100 mM Tris HCl, 200 mM NaCl, 10% glycerol, 10 mM imidazole, pH 8.0). The protein was eluted with 100 mM imidazole in HisTrap buffer. The EnH-eGFP protein was purified further with Nuvia™ cPrime™ Hydrophobic Cation Exchange Media (BioRad #1563401), equilibrated with cation exchange buffer (100 mM Tris HCl, 100 mM NaCl, 10% glycerol, pH 6.8), and eluted with 400 mM NaCl in cation exchange buffer. Following cation exchange, the samples were purified using HiLoad Superdex 200 PG column (Cytiva #28989335) at the Histone Source at Colorado State University. The fractions containing EnH-eGFP protein were pooled, concentrated using an Amicon Ultra-15 10 kDa MWCO centrifugal filter unit (EMD Millipore), and dialyzed with EnH-eGFP storage buffer (100 mM Tris HCl, 200 mM NaCl, 10% glycerol, pH 8.0). EnH-eGFP protein was collected, concentrated to 42 mg/mL, flash frozen, and stored at −80°C.

Protein purification steps were analyzed with SDS-PAGE (NuPAGE 4−12% Bis-Tris Gel, ThermoFisher #NP0321BOX) with 2-(N-morpholino)ethanesulfonic acid (MES) pH 7.3 SDS running buffer (NuPAGE MES SDS Running Buffer #NP0002) and stained with Imperial Protein Stain (ThermoFisher #24615). Bradford Assay using Coomassie Plus Protein Assay Reagent (ThermoFisher #23238) was used to determine final protein concentrations.

***bZip and C-clamp Synthesis***

bZip protein (Fig. S6) was commercially synthesized (GenScript) and resuspended in 100 mM sodium chloride,100 mM sodium phosphate (pH 7.4). Following resuspension, samples were incubated on ice for 12 hours to promote disulfide bond formation prior to downstream applications.

C-clamp protein (Fig. S7) was commercially synthesized (LifeTein) in two variants: an unlabeled construct and a C-terminal TAMRA-conjugation construct. Lyophilized material was resuspended in 50 mM Tris-HCl (pH 7.4) supplemented with 1 mM $ZnCl_2$. To remove excess zinc, samples were concentrated and buffer-exchanged using Amicon Ultra centrifugal filter units (3 kDa MWCO; Millipore) prior to downstream applications

***DNA Duplex Annealing***

Individual oligomers were synthesized and HPLC purified by Integrated DNA Technologies. The DNA oligomer sequences used for co-crystallization are listed in Tables S1 and S4. With limited exceptions, each oligomer contained the 19-bp iteron sequence TGTGACAAATTGCCCTCAG, but typically also contained the core 21-bp DNA block used in PDB:1REP from Komori *et al*. (38): CCTGTGACAAATTGCCCTCAG (omitting a dangling 3'_T). In Table S1, the standard 21-bp footprint is shaded in gray, and (purposeful) deviations are shaded in cyan. Flanking variable DNA sequences depended on the expansion scheme and desired sequence. Notable potential guest-binding sequences are marked with underlines. Potential HhaI restriction sites are marked with an orange font. The oligomers were resuspended in a standardized buffer (henceforth CC1 oligo buffer: 50 mM Tris HCl, 100 mM KCl, pH 6.7) and combined in equal molar ratio (1:1) with the complementary strand. The strands were annealed by heating to 94 °C for 2 min and slowly cooling to room temperature over approximately 60 minutes. The final concentration of all CC1 dsDNA blocks following duplexing was 4 mM.

***Scaffold Protein: DNA Complex Crystallization***

The scaffold protein and DNA were mixed in a 1.2:1 ratio (440:364 µM, 55 µL total) and were incubated on ice for 30 minutes prior to crystallization via sitting drop vapor diffusion. Crystallization trials used CrysChem sitting-drop plates (Hampton Research HR3-159), with 400 µL reservoir solution per well and 4 µL crystallization drops prepared by mixing 2 µL each of the protein–DNA complex and reservoir solution directly on the platform; drops were mixed by gentle pipetting. Plates were sealed immediately after setup and incubated at 25 °C under static conditions. Crystallization conditions for both the porous lattice ($CC1^{+10}$) and for the large-porous lattice ($CC1^{+21}$) were 10-80 mM Magnesium Acetate, 0.3-1.8 M Lithium Sulfate, and 50 mM MES pH 6.5 (or occasionally 50 mM sodium cacodylate pH 6.5). A detailed list of individual crystals, PDB codes, and corresponding growth conditions are reported in Table S2. Crystals grew to full size in a range of 24 hours to 30 days (most often in 1-2 days). Crystal images were taken with a Moticam X5 Plus camera attached to a Motic SMZ-168 stereozoom microscope.

***X-Ray Diffraction Data Collection and Refinement***

Crystals were flash frozen in liquid nitrogen after briefly transferring the crystal to a cryo-protectant solution (a 1:1 mixture of the reservoir solution and 50% glycerol stock for a final concentration of 25% glycerol). Single crystal X-Ray diffraction data was collected at either ALS beamline 4.2.2 (CMOS detector), ALS beamline 8.2.2 (Pilatus3 detector), ALS beamline 8.2.1 (Dectris EIGER2 detector), or APS beamline 24-IDE (Dectris EIGER2 detector) from 0 to 180° with an omega delta of 0.2° and an exposure of 0.5 s or 0.25 s. Data was processed with XDS (75) and refined using PHENIX and COOT (76,77). The first porous $CC1^{+10}$ dataset (PDB 9Z08) was refined using newly generated $R_{Free}$ flags. All subsequent isomorphous structures were refined using the same $R_{Free}$ flags to preserve a fixed crossvalidation benchmark across datasets. Decisions about truncating the structure factor data were made on a case-by-case basis, jointly considering the following thresholds $CC_{1/2} > 0.3$, mean $\langle I/\sigma(I) \rangle > 1.0$, and completeness > 85% as target guidelines for the highest-resolution shell. Most datasets satisfy all three thresholds in the highest-resolution shell. For a small number of datasets (9ZYB, 9YZE, 9Z08, 9Z1A, and 9Z44), resolution limits were selected such that no more than one or two of the metrics fell below the target thresholds, relying on modern maximum likelihood refinement methods to appropriately down-weight the marginal data (78,79).

Omit maps were generated for every model containing a guest protein structure (9YZL, 9YZM, 9YZN, 9YZO, 9YZP, 9YZQ, 9YZR, 9YZS, 9YZT, 9YZU, 9Z44, 9Z4E) using PHENIX composite omit map with simulated annealing (45–47). Figure S15 shows one representative omit map example per guest type.

#### *EDC Ligation*

Co-crystals were washed in conditions similar to their growth conditions (0.3-2.0 M Lithium sulfate, 20-50 mM Magnesium acetate, and 50 mM MES, pH 6.5). The co-crystals were washed in 200 µL on 9-well glass plates (Hampton) to remove additional protein and DNA monomers. Note: For EDC ligation it is important to avoid buffers with 1°/2° amines or carboxylic acids. 1-Ethyl-3(3-dimethylaminopropyl)carbodiimide (EDC) (Advanced Chemtech CAS#:25952-53-8) was resuspended in the crosslinking solution (1.0 M Lithium sulfate, 50 mM Magnesium acetate, and 50 mM MES, pH 6.0) to final concentration values of 30-60 mg/mL and used immediately. The co-crystals were crosslinked in a 200 µL EDC solution volume overnight (16-24 hrs). The crosslinking reaction (Fig. S26) was ended by looping the co-crystals back into the wash solution.

#### *Fluorescence Polarization*

The first DNA oligomer sequences used for fluorescence polarization measurements (Fig. S9) matched a full $CC1^{+10}$ DNA block (31 bp) , contained the canonical binding site (5'-TAATTA-3') for a guest homeodomain, lacked terminal phosphate modification, and had one strand modified with a 3′-terminal 6-carboxyfluorescein (FAM) for fluorescence detection. A second truncated DNA duplex (11 bp) was prepared to probe binding in the absence of the RepE54 binding site. Individual oligomers were again synthesized by Integrated DNA Technologies, and purified via standard desalting. The oligomers were resuspended in CC1 oligo buffer (50 mM Tris HCl, 100 mM KCl, pH 6.7) and combined in equal molar ratio (1:1) with the complementary strand. The strands were annealed by heating to 94 °C for 2 min and then allowed to cool to room temperature for one hour. The stock solution concentration of all annealed FP duplexes was 1 mM. Prior to 1:1 mixing with the protein (see below), FP duplexes were diluted to a concentration of 20 nM in the solvent conditions being tested.

Experiments were carried out in black, flat-bottom 384-well plates (Greiner Bio-One, Ref#:781900) in a final volume of 30 μL per well (15 μL FP duplex and 15 μL of protein). Each well contained 10 nM final concentration of FP duplex in binding buffer. Guest proteins were serially diluted in their storage buffer starting from 50 μM, generating a 15-point dilution series (final concentration range from 50 μM to ~1.5 nM), with one well containing no protein as a fluorescence polarization baseline.

After adding guest protein, the plates were incubated at room temperature for 30-45 minutes to allow binding equilibrium to be reached. FP was measured on a CLARIOStar Plus (BMG Laboratories) plate reader using a 485 nm Ex: 535 nm Em filter. Initial readings were performed in duplicate (Fig. S10AB). Later readings were measured with eightfold replication (Fig. S10C). Baseline polarization values were measured using protein-free blanks. Fluorescence polarization data were analyzed using a custom Python script (https://github.com/cdsnow/FPfitting) that averages replicate measurements, calculates standard errors, and fits each titration curve to a 1:1 binding model (Fig. S10). The script fits the protein–DNA complex concentration at each titration point, incorporating the fixed final 10 nM concentration of fluorescent duplex via the quadratic Morrison equation to account for ligand depletion. Nonlinear regression, weighted by the inverse variance (with a minimum uncertainty threshold of 5% of the dynamic range to prevent overweighting), was used to simultaneously optimize $K_d$, the baseline polarization value, and the bound-state polarization value. One outlier data point at 200 mM NaCl and 6.25 μM was excluded from the analysis. The resulting fits (Fig. S10) were plotted on a log-scaled protein concentration axis and used to estimate $K_d$ values for the salt scouting conditions. The provided Python fitting scripts also allow simple Langmuir binding equation fitting. We omit the simple Langmuir fits here since the results were redundant with the more sophisticated model (Fig. S10).

***Isothermal Titration Calorimetry (ITC)***

The binding affinity of EVE-HD and UBX-HD to dsDNA sequences R1 and R2 (Fig. S16) were measured using an Affinity ITC Auto Low Volume System (190 μL) equipped with a gold cell (TA Instruments; New Castle, DE). ITC runs were performed and controlled using the built-in ITCRun software. The sample cell (set to 25°C) containing 20 μM HD in 20 mM Tris, 50 mM KCl, 4 mM $CaCl_2$ (pH 7.4) was titrated with 2 μL x 20 injections of 200 μM dsDNA. All runs were performed in duplicate to ensure reproducibility. Isotherms and titration curves were plotted using Microsoft Excel.

Python scripts (https://github.com/cdsnow/FPfitting) were used to fit the data and obtain thermodynamic parameters for the HD-dsDNA interaction (i.e, $K_d$, binding stoichiometry, ΔH, and ΔS). Specifically, integrated heat data (kJ $mol^{-1}$ of injectant vs. molar ratio) were fit to the Wiseman isotherm, a closed-form model for 1:1 binding (80). The four adjustable parameters were the apparent stoichiometry (N), a dimensionless affinity parameter $c = K_a$ [M] (the isotherm is algebraically equivalent to the traditional parameterization with $c = n K_a$ [M]), the molar enthalpy of binding (ΔH), and a constant offset accounting for the heat of dilution. Nonlinear least-squares regression was performed with the trust-region reflective algorithm as implemented in scipy.optimize.curve_fit (SciPy v1.16.3), using unweighted residuals. The macromolecule concentration was fixed at 20 μM for all experiments and the temperature was fixed at 298.15 K. Thermodynamic quantities were derived from the fitted parameters using standard relationships: $K_d = [M]/c$, $\Delta G = RT \ln K_d$, and $-T\Delta S = \Delta G - \Delta H$.

Uncertainties in the fitted parameters were obtained from the diagonal of the covariance matrix returned by the least-squares optimizer. For derived quantities that depend on multiple fitted parameters (notably -TΔS, which depends on both c and ΔH), uncertainties were propagated analytically using the full covariance matrix, including off-diagonal terms, to account for parameter correlations. As an independent check, residual-resampling bootstrap analysis (81) was performed with 1,000 trials per dataset: residuals from the best-fit model were resampled with replacement, added back to the model prediction to generate synthetic datasets, and each synthetic dataset was re-fit to yield a bootstrap distribution of all parameters. Bootstrap standard deviations (with Bessel's correction) served as the reported uncertainties.

***Guest Loading***

Crosslinked co-crystals containing DNA sequences for guest proteins were washed at room temperature for 1 hour to remove components that could interfere with guest protein binding and to equilibrate the crystals to the impending guest loading soak. We used wash solution that matched the subsequent guest loading buffers, and these guest loading buffers (Table S7) were varied to 1) demonstrate the capacity of the crystals to diffract in different conditions and 2) determine how the salt concentration changed the resulting electron density map. Accordingly, the wash solutions consisted of 15-50 mM KCl, 2-4 mM $CaCl_2$, 10 mM Tris HCl, pH 7.4, and 10% glycerol. A fresh 180 µL of guest loading / wash solution was mixed with 20-30 µL of guest protein, and the washed co-crystals were moved into the guest loading solution using cryo loops (the concentrations in Table S7 are the final concentrations after mixing). The co-crystals sitting in guest loading solutions were sealed for 24-72 hours to allow guest proteins to diffuse and reach binding equilibrium within the crystal lattice.

***NHS-fluorescein conjugation***

UBX-HD was dialyzed into labeling buffer (50 mM $Na_2PO_4$, 300 mM NaCl, pH 8.1) using Slide-A-Lyzer™ dialysis pucks (1 kDa MWCO; ThermoFisher) to remove free amines and equilibrate the sample for N-hydroxysuccinimide (NHS) conjugation. The protein concentration was 200 µM prior to labeling. NHS-fluorescein (Thermo Scientific; 5 mg/mL, ~10 mM) was added at a 1:5 dye:protein molar ratio, and the reaction was incubated overnight at 4 °C with gentle agitation. With five surface lysine residues and the N-terminus, the dye conjugation location and extent is presumably heterogeneous. To minimize photobleaching, dye-handling steps were performed in opaque 1.5 mL microcentrifuge tubes and under low-light conditions. Unreacted dye was removed by dialysis against fresh labeling buffer (50 mM $Na_2PO_4$, 300 mM NaCl, pH 8.1, 1 kDa MWCO) with multiple buffer exchanges until no residual dye was visible in the dialysate. The labeled protein was concentrated to the working concentration using a 3 kDa MWCO Amicon centrifugal filter unit (Millipore) and stored on ice for immediate use, or aliquoted and flash-frozen for long-term storage at −80 °C

***Confocal Microscopy***

Crystals loaded with fluorescently labeled UBX-HD were imaged using a Nikon Eclipse Ti Yokagawa CSU-X1 spinning-disk confocal microscope equipped with an AndoriXon Ultra 879U EMCCD camera. Fluorescein signal was collected using 488 nm laser excitation and a 515 nm emission filter. Image acquisition and processing were performed using Nikon NIS-Elements software.

For the confocal microscope experiments reported in Fig. S12, Fig. S13, and Fig. S14, a large crystal was incubated for 24 hours in a drop with 8 µL volume, a 15 mM KCl, 4 mM $CaCl_2$, 10 mM Tris HCl, pH 7.4, and 10% glycerol buffer and a UBX-HD concentration (combined unlabeled and trace labeled) of 28 µM. Images were acquired using a 10x objective lens, and Z-stacks were collected using 2.5 µm step sizes for a total of 37 steps in all experiments. Laser power exposure time, and all other optical parameters were held constant across experiments to enable direct comparison of fluorescence intensity between crystals.

For the experiments reported in Fig. 2, fluorescein labelled UBX-HD was diluted in unlabeled UBX-HD protein (1:200). The final drop had a volume of 10 µL, a 15 mM KCl, 4 mM $CaCl_2$, 10 mM Tris HCl, pH 7.4, and 10% glycerol buffer and a UBX-HD concentration (combined unlabeled and trace labeled) of 23 µM. Images were acquired using a 20x objective lens, and Z-stacks were collected using 1 µm step sizes. Laser power exposure time, and all other optical parameters were held constant across imaging time points. The exception is the 24 hr z-stack, where laser power was reduced from 20% to 12% to avoid over-saturation.

***AlphaFold-3 modeling of guest proteins into the CC1$^{+10}$ lattice***

To evaluate potential guest installation sites in the porous CC1$^{+10}$ scaffold, we used AlphaFold-3 (AF3)(40) in combination with PyMOL (82) to model guest protein binding within the context of the full crystal lattice. The protocol below outlines the computational workflow used to assess guest compatibility with the CC1$^{+10}$ lattice. Using the AF3 server (https://alphafoldserver.com), we created prediction jobs that included: (i) the RepE54 scaffold protein sequence, (ii) the full CC1$^{+10}$ insert DNA sequence containing the intended guest-binding site, and (iii) the sequence of the DBD. For each guest family, we tested multiple registers by inserting the canonical DNA-binding motif (e.g., 5′-TAATTA-3′ for homeodomains) into different positions along the insert DNA.

For each structure prediction target, all five models output by AF3 were visually inspected for consistency in PyMOL. The AF3-predicted structures were then aligned in PyMOL to superimpose the AF3 RepE54 onto the corresponding RepE54 chain in a representative CC1$^{+10}$ asymmetric unit (the most similar available model). After saving the superposed AF3 model, we manually added the CRYST1 record from a representative experimental CC1$^{+10}$ PDB file to the AF3 model header. We then used a PyMOL supercell script (https://pymolwiki.org/index.php/Supercell) to expand the asymmetric unit into a full lattice. The resulting lattice environment was inspected to assess steric clashes, crystal packing compatibility, and accessibility of the guest protein to solvent channels. Non-clashing models were prioritized for experimental testing based on which sites yielded high AF3 confidence for guest binding to the target site (i.e. high pLDDT values for the guest).

To help identify guest DBD that are compatible with the scaffold crystals (i.e. not too large), and to facilitate the selection of guest binding sites, we have prepared Python scripts to automate sliding window superposition of candidate DBD:cognate-DNA models to scan all possible registers. The scripts (https://github.com/cdsnow/DBDscan) assess candidate installation sites on the basis of potential interactions with symmetry mates. Additionally, as shown in Figure S23, a web application version of the scripts (https://www.engr.colostate.edu/~cdasnow/guestscan.html) allows interested parties to assess where their DBD of interest might be installed within the $CC1^{+10}$ or $CC1^{+21}$ scaffolds.

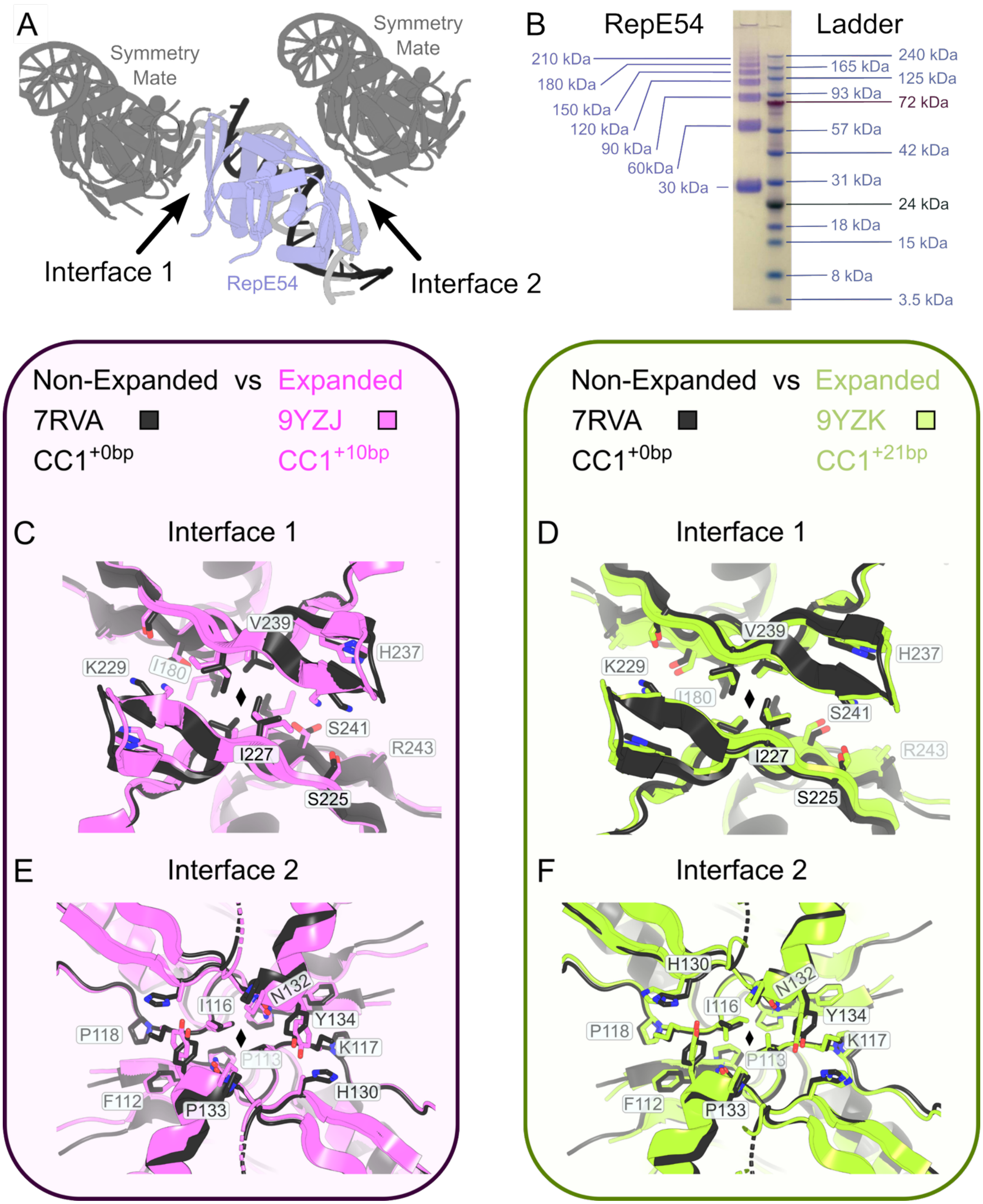

A
Symmetry Mate
Symmetry Mate
Interface 1
RepE54
Interface 2
B
RepE54
Ladder
210 kDa
180 kDa
150 kDa
120 kDa
90 kDa
60kDa
30 kDa
240 kDa
165 kDa
125 kDa
93 kDa
72 kDa
57 kDa
42 kDa
31 kDa
24 kDa
18 kDa
15 kDa
8 kDa
3.5 kDa
Non-Expanded vs Expanded
7RVA
9YZJ
CC1+0bp
CC1+10bp
C
Interface 1
V239
H237
K229
I180
S241
R243
I227
S225
E
Interface 2
N132
I116
Y134
P118
K117
P113
F112
P133
H130
Non-Expanded vs Expanded
7RVA
9YZK
CC1+0bp
CC1+21bp
D
Interface 1
V239
H237
K229
I180
S241
R243
I227
S225
F
Interface 2
H130
N132
I116
Y134
P118
K117
P113
F112
P133

**Figure S1. Conservation of protein-protein interfaces in expanded CC1 lattices**

**(A)** From PDB:7RVA, a depiction of RepE54 (blue) with cognate DNA (black/gray) with symmetry mates across two distinct protein-protein interfaces. **(B)** Denaturing conditions were robust: incubation at 90°C for 10 min in storage buffer (100 mM sodium citrate pH 6.2, 100 mM KCl, 10 mM $MgCl_2$, and 10% glycerol) mixed 3:1 with 4X lithium dodecyl sulfate (LDS) ThermoFisher #NP0008). However, subsequent PAGE at 250 V at 25°C for 45 min on a RepE54 aliquot revealed distinct oligomer bands up to 7-mers, suggesting a notable propensity for unbounded protein oligomerization, consistent with utility as a building block for scaffold crystal growth. **(C)** Interface 1 (aligned residues 175-222) was very similar (0.52Å $RMSD_{C\alpha}$ superposition) in the non-expanded lattice (PDB:7RVA, black) and the $CC1^{+10}$ lattice (PDB:9YZJ, pink) but did show sidechain changes in the interfacial hydrophobic core. Given the 2-fold axis (◆), each interfacial residue appears twice but is labeled only once for visual clarity. **(D)** Within the limitations of the lower-resolution (5.2Å) $CC1^{+21}$ lattice (PDB:9YZK, lime), Interface 1 (aligned residues 175-222) was still very similar (0.69Å $RMSD_{C\alpha}$ superposition) to the non-expanded lattice (PDB:7RVA, black). **(E)** Interface 2 (aligned residues 87-95 and 110-117) was very similar (0.48Å $RMSD_{C\alpha}$ superposition) in the non-expanded lattice (PDB:7RVA, black) and the $CC1^{+10}$ lattice (PDB:9YZJ, pink) **(F)** For PDB:9YZK (lime), relative to Interface 1, Interface 2 was again subtly but noticeably more similar (0.62Å $RMSD_{C\alpha}$ superposition) to the non-expanded lattice (PDB:7RVA, black). Figure prepared in PyMOL (82) and Inkscape.

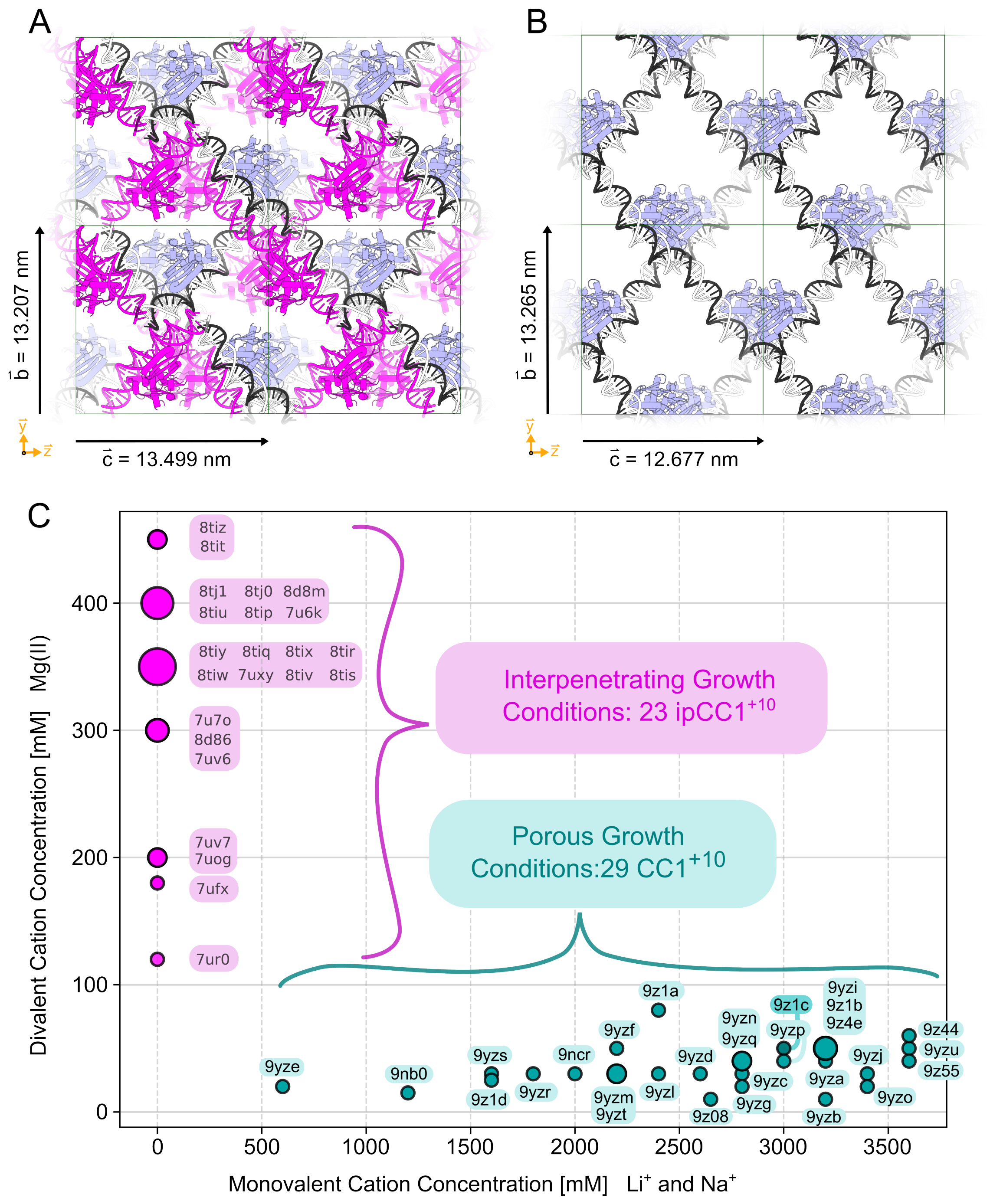
A
B
C
b = 13.207 nm
c = 13.499 nm
b = 13.265 nm
c = 12.677 nm
8tiz
8tit
8tj1 8tj0 8d8m
8tiu 8tip 7u6k
8tiy 8tiq 8tix 8tir
8tiw 7uxy 8tiv 8tis
7u7o
8d86
7uv6
7uv7
7uog
7ufx
7ur0
Interpenetrating Growth Conditions: 23 ipCC1+10
Porous Growth Conditions:29 CC1+10
9z1a
9yzi
9z1b
9z4e
9z1c
9yzn
9yzq
9yzp
9yzf
9z44
9yzu
9z55
9yze
9nb0
9yzs
9ncr
9yzd
9yzj
9yzc
9yza
9z1d
9yzr
9yzm
9yzt
9yzl
9z08
9yzg
9yzb
9yzo
Divalent Cation Concentration [mM] Mg(II)
Monovalent Cation Concentration [mM] Li+ and Na+
0
100
200
300
400
500
1000
1500
2000
2500
3000
3500

**Figure S2. Interpenetrating versus porous CC1 lattices**

**(A)** Four unit cells from the interpenetrating CC1 lattice (ipCC1$^{+10}$, I222 space group, PDB 7U6K), where one copy of the lattice (blue, white, black) is interwoven with another copy rotated by 180° (magenta). **(B)** Four unit cells from the porous CC1$^{+10}$ lattice (I121 space group) shown with the major solvent channels perpendicular to the image (PDB 9YZJ). Close packing of the DNA in the ipCC1 lattice occurs at a higher magnesium concentration. For example, the 7U6K sitting drop growth condition was 400 mM $MgCl_2$, 24% PEG 400, and 80 mM Tris HCl pH 8.0 while the 9YZJ growth condition for sitting drops was 30 mM magnesium acetate, 1.7 M $Li_2SO_4$, and 50 mM MES pH 6.5. **(C)** Comparison of the divalent and monovalent cation concentrations in the growth conditions for interpenetrating (magenta) and porous crystals (teal). Among porous CC1$^{+10}$ crystals (Table S2) the growth condition magnesium concentration spanned 10-80 mM. The magnesium concentration was much higher for our earlier 23 ipCC1$^{+10}$ PDB entries (spanning 120 mM to 450 mM Mg(II), but most often 300-400 mM). See Figure S3 for a molecular explanation of the role of Mg(II) in the interpenetrating crystal form.

These findings have parallels with other nanoscale DNA self-assembly processes. For example, the folding yield and quality of close-packed individual "3D" DNA origami objects is much more sensitive to the presence and quantity of divalent counterions than single-layer, "planar" origami (83) and the morphology of 3D crystals consisting of DNA origami can be programmed by Mg(II) concentration (84). Lastly, see our prior work (35) for an illustration of the idea that a crystal need not be 100% porous nor 100% interpenetrating (imagine an interpenetrating lattice where some regions lack one of the two sub-lattices. Figure prepared in PyMOL (82) and Inkscape.

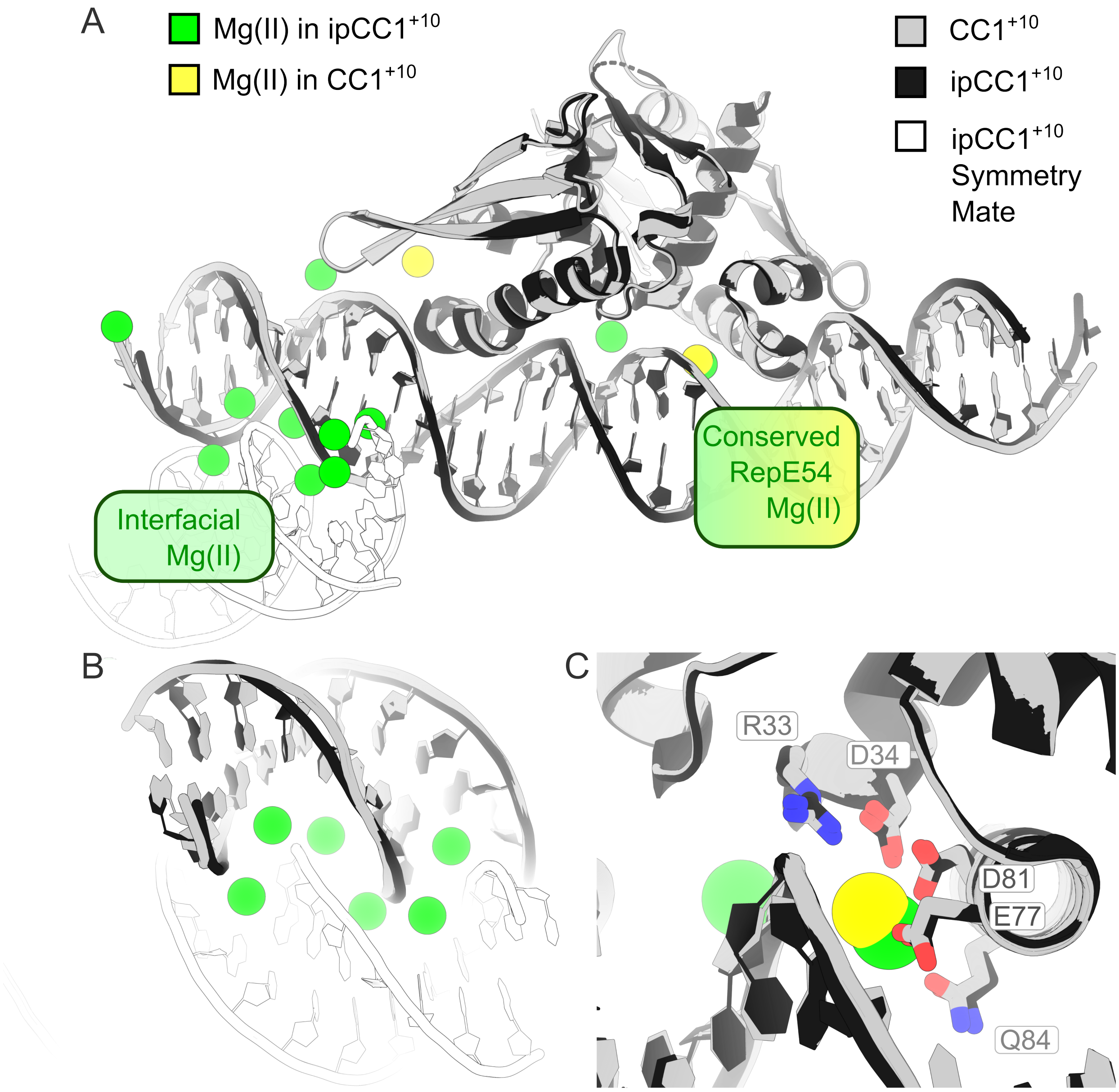

A
Mg(II) in ipCC1+10
Mg(II) in CC1+10
CC1+10
ipCC1+10
ipCC1+10 Symmetry Mate
Conserved RepE54 Mg(II)
Interfacial Mg(II)
B
C
R33
D34
D81
E77
Q84

**Figure S3. Comparison of CC1$^{+10}$ and ipCC1$^{+10}$ resolved ion coordinates**

A key difference between ipCC1 growth and CC1 growth was the reduction in magnesium concentration. **(A)** Overview, showing an entire asymmetric unit for CC1$^{+10}$ (gray, PDB:9NCR) superimposed onto ipCC1$^{+10}$ (black, PDB:7U6K), where we also show the ipCC1$^{+10}$ interpenetrating symmetry mate (white). The ipCC1$^{+10}$ magnesium ions (green) dramatically outnumber the CC1$^{+10}$ magnesium ions (yellow) and sodium ion (purple). **(B)** Close-up view of the interdigitating DNA in ipCC1$^{+10}$ , and six interfacial Mg(II). Thus, interfacial magnesium contacts appear to drive the close packing of DNA in the interpenetrating lattice. **(C)** We note that complete removal of divalent cations could hinder RepE54 binding, since one Mg(II) exists at the interface of RepE54 and the DNA, coordinated by R33, D34, E77, D81, and Q84. Figure prepared in PyMOL (82) and Inkscape.

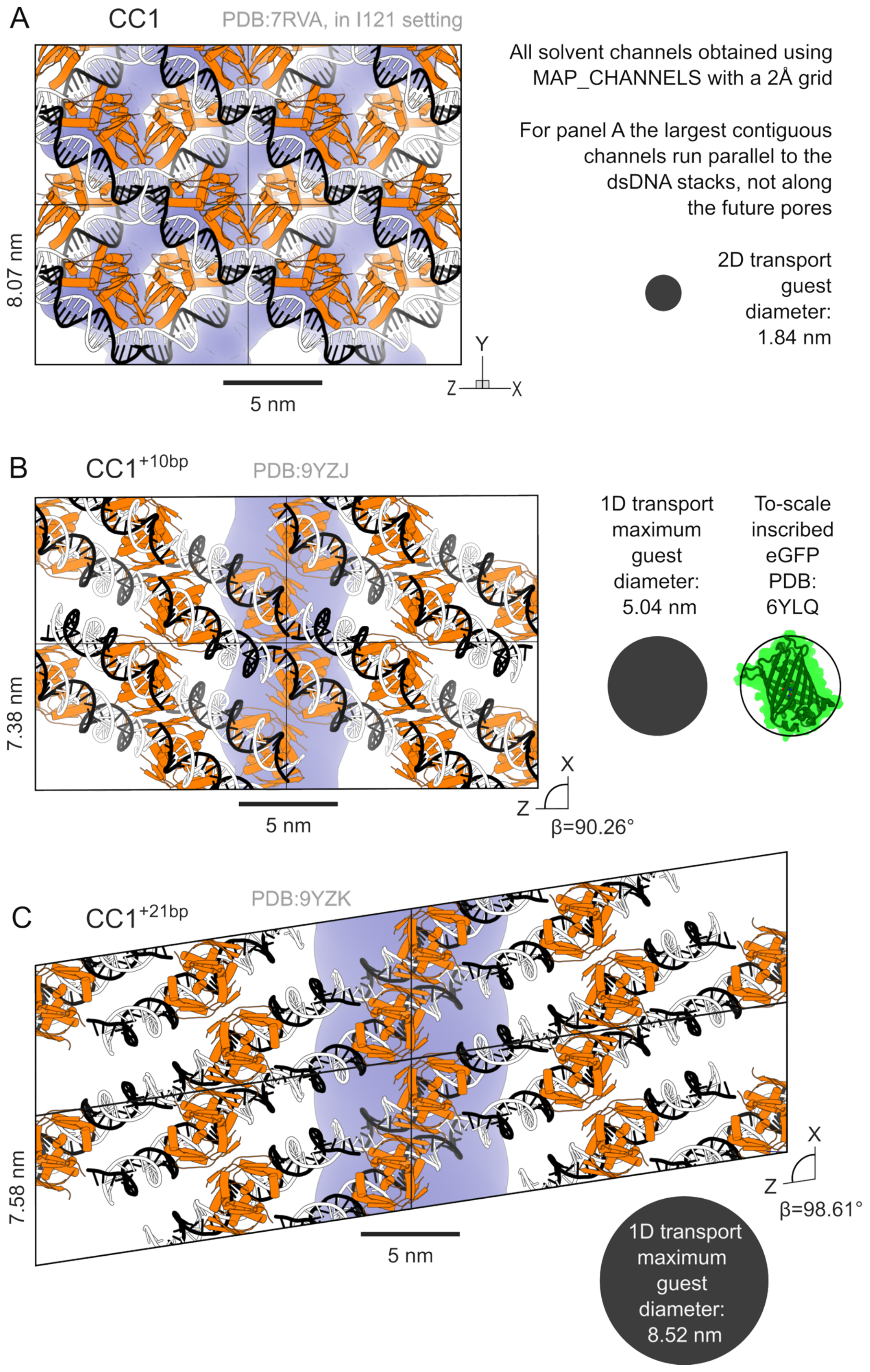
A
CC1
PDB:7RVA, in I121 setting
All solvent channels obtained using MAP_CHANNELS with a 2Å grid
For panel A the largest contiguous channels run parallel to the dsDNA stacks, not along the future pores
2D transport guest diameter: 1.84 nm
8.07 nm
5 nm
Y
Z
X
B
CC1+10bp
PDB:9YZJ
1D transport maximum guest diameter: 5.04 nm
To-scale inscribed eGFP PDB: 6YLQ
7.38 nm
5 nm
X
Z
β=90.26°
C
CC1+21bp
PDB:9YZK
7.58 nm
5 nm
X
Z
β=98.61°
1D transport maximum guest diameter: 8.52 nm

**Figure S4. MAP_CHANNELS analysis of CC1 family of crystals**

**(A)** MAP_CHANNELS (67) analysis of non-expanded CC1 (PDB 7RVA) using a 2Å grid revealed that the largest contiguous solvent channels ran along the coaxial dsDNA columns and were too small for macromolecular intra-crystal diffusion. **(B)** MAP_CHANNELS analysis of $CC1^{+10}$ (specifically PDB:9YZJ for this figure) using a 2 Å grid resulted in a calculated pore diameter of 5.04 nm. As a size comparison, we show to-scale eGFP (PDB:6YLQ) inscribed in a 5.04 nm circle, with an orientation selected to maximize the visible eGFP cross-section. **(C)** MAP_CHANNELS analysis of $CC1^{+21}$ using a 2 Å grid resulted in a calculated pore diameter of 8.52 nm (PDB 9YZK). See Table S8 for MAP_CHANNELS analysis of *all* new structures. Notably, the ability to tightly control the size of the solvent channels within the CC1 family of crystals may provide useful molecular sieve functionality (15). Figure prepared in PyMOL (82) and Inkscape.

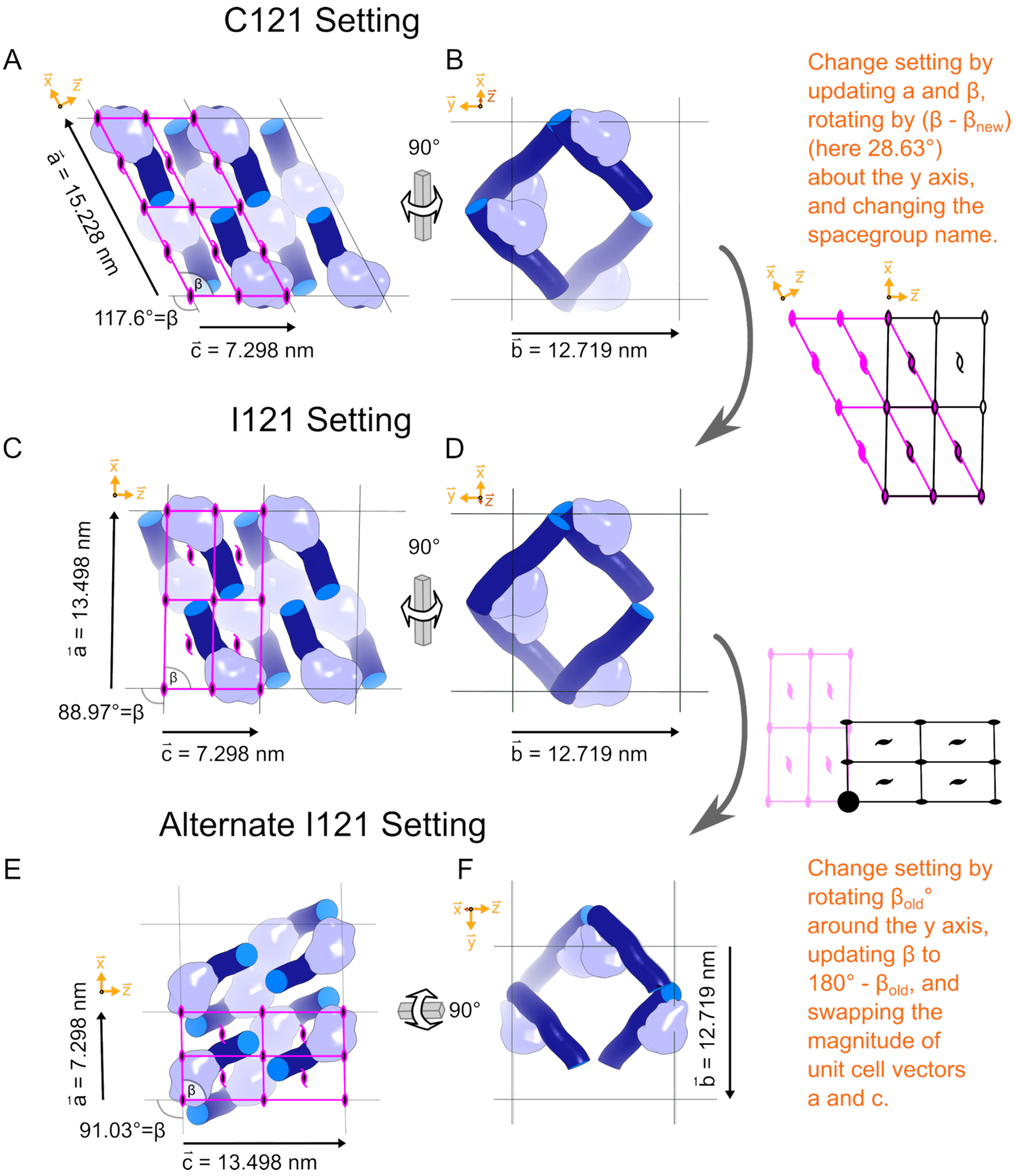
C121 Setting
A
B
a = 15.228 nm
117.6°=β
c = 7.298 nm
90°
b = 12.719 nm
Change setting by updating a and β, rotating by (β - βnew) (here 28.63°) about the y axis, and changing the spacegroup name.
I121 Setting
C
D
a = 13.498 nm
88.97°=β
c = 7.298 nm
90°
b = 12.719 nm
Alternate I121 Setting
E
F
a = 7.298 nm
91.03°=β
c = 13.498 nm
90°
b = 12.719 nm
Change setting by rotating βold° around the y axis, updating β to 180° - βold, and swapping the magnitude of unit cell vectors a and c.

**Figure S5. Equivalent descriptions of the CC1 expanded lattice.**

This lattice setting schematic uses a theoretical $CC1^{+10}$ model (expanded 7RVA). The monoclinic space group (#5) has several conventions. Typically, C121 is favored over I121, obtuse β angles are preferred to acute, and the smallest reduced cell is favored. In our case, the C121 setting **(A)** is convenient for depiction of the lattice on the page because the coaxial DNA columns vectors lie in the plane perpendicular to the z-axis (which is not coincident with unit cell vector *c*). Starting from this standard view, a 90° rotation **(B)** puts the y-axis into the horizontal direction, and directs the c vector towards the reader, revealing the nanopores parallel to the unit cell vector *c*. However, the I121 setting **(C)** is also useful because the β value is close to 90°, and the I121 setting facilitates comparison to the I222 space group (adopted by the former interpenetrating lattice, e.g. PDB:7U6K) where β = 90° exactly. Also, the unit cell vector *c* that is parallel to the major nanopores **(D)** is now nearly aligned to the Cartesian z-axis. Finally, since the near-90° β values for CC1 crystals may sometimes fall below 90° (as shown here), they can violate the obtuse β convention. Therefore, one alternate setting is to rotate about the unique monoclinic y-axis (with the 2-fold axes) so that what used to be the *c* vector becomes an *a* vector aligned to the x-axis. **(E)** In this alternate setting, the nanopores are parallel to the *a* vector, which is aligned to the x-axis. The standard depiction of monoclinic cells (i.e. in space group diagrams) aligns the +*c* vector to the horizontal, rather than aligning the +*a* vector to the vertical. Thus, a horizontal 90° rotation **(F)** reveals the nanopores, since the x-axis is nearly perpendicular to the image. Figure prepared in PyMOL (82) and Inkscape.

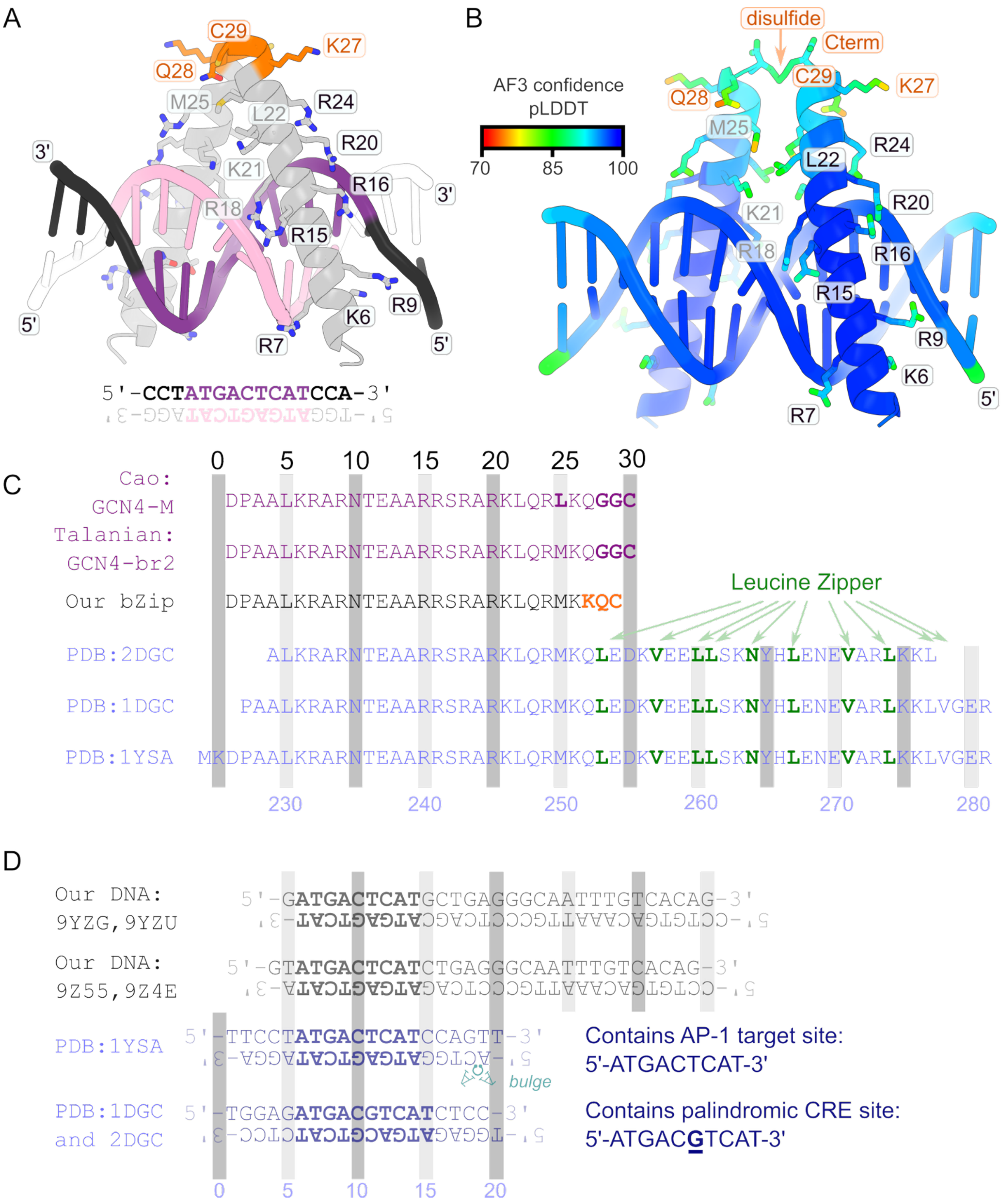

A
C29
K27
Q28
M25
L22
R24
R20
K21
R16
R18
R15
R9
K6
R7
3'
5'
5'-CCTATGACTCATCCA-3'
B
disulfide
Cterm
AF3 confidence
pLDDT
70
85
100
C
Cao:
GCN4-M
DPAALKRARNTEAARRSRARKLQRLKQGGC
Talanian:
GCN4-br2
DPAALKRARNTEAARRSRARKLQRMKQGGC
Our bZip
DPAALKRARNTEAARRSRARKLQRMKKQC
Leucine Zipper
PDB:2DGC
ALKRARNTEAARRSRARKLQRMKQLEDKVEELLSKNYHLENEVARLKKL
PDB:1DGC
PAALKRARNTEAARRSRARKLQRMKQLEDKVEELLSKNYHLENEVARLKKLVGER
PDB:1YSA
MKDPAALKRARNTEAARRSRARKLQRMKQLEDKVEELLSKNYHLENEVARLKKLVGER
0 5 10 15 20 25 30
230 240 250 260 270 280
D
Our DNA:
9YZG,9YZU
5'-GATGACTCATGCTGAGGGCAATTTGTCACAG-3'
Our DNA:
9Z55,9Z4E
5'-GTATGACTCATCTGAGGGCAATTTGTCACAG-3'
PDB:1YSA
5'-TTCCTATGACTCATCCAGTT-3'
bulge
PDB:1DGC
and 2DGC
5'-TGGAGATGACGTCATCTCC-3'
Contains AP-1 target site:
5'-ATGACTCAT-3'
Contains palindromic CRE site:
5'-ATGACGTCAT-3'
0 5 10 15 20

**Figure S6: Design of the bZip construct.**

Inspired by Cao *et al.* (85) and Talanian *et al*. (86), we designed a 29-a.a. peptide where a C-terminal disulfide bond is intended to replace the coiled-coil segment**. (A)** A homology model for our bZip construct was prepared in PyMOL by truncating PDB:1YSA and manually selecting the most common rotamer for the changed amino acids in the 3 C-terminal residues (orange). Each residue appears twice in the dimer but is labeled only once for visual clarity. **(B)** An AlphaFold3 (40) model for the same sequences. Confidence was very high (pLDDT > 90) in the DNA and DNA-binding region and was still high in the C-termini of the helices, including pLDDT > 87 for the disulfide bond. **(C)** Our bZip construct design was guided by AF3, but was also inspired by Cao *et al.* (85) variant GCN4-M and Talanian *et al.* (86) variant GCN4-br2 where a C-terminal suffix (GGC) enhanced binding affinity via disulfide bond formation to compensate for deletion of the leucine zipper region (green). **(D)** Our bZip capture DNA sequence was the AP-1 target sequence, added as an asymmetric expansion of the RepE54 scaffold sequence. Notably, the bottom strand 5' end in this schematic is the top strand residue 1 by our usual numbering and depiction convention (Table S9). Figure prepared in PyMOL (82) and Inkscape.

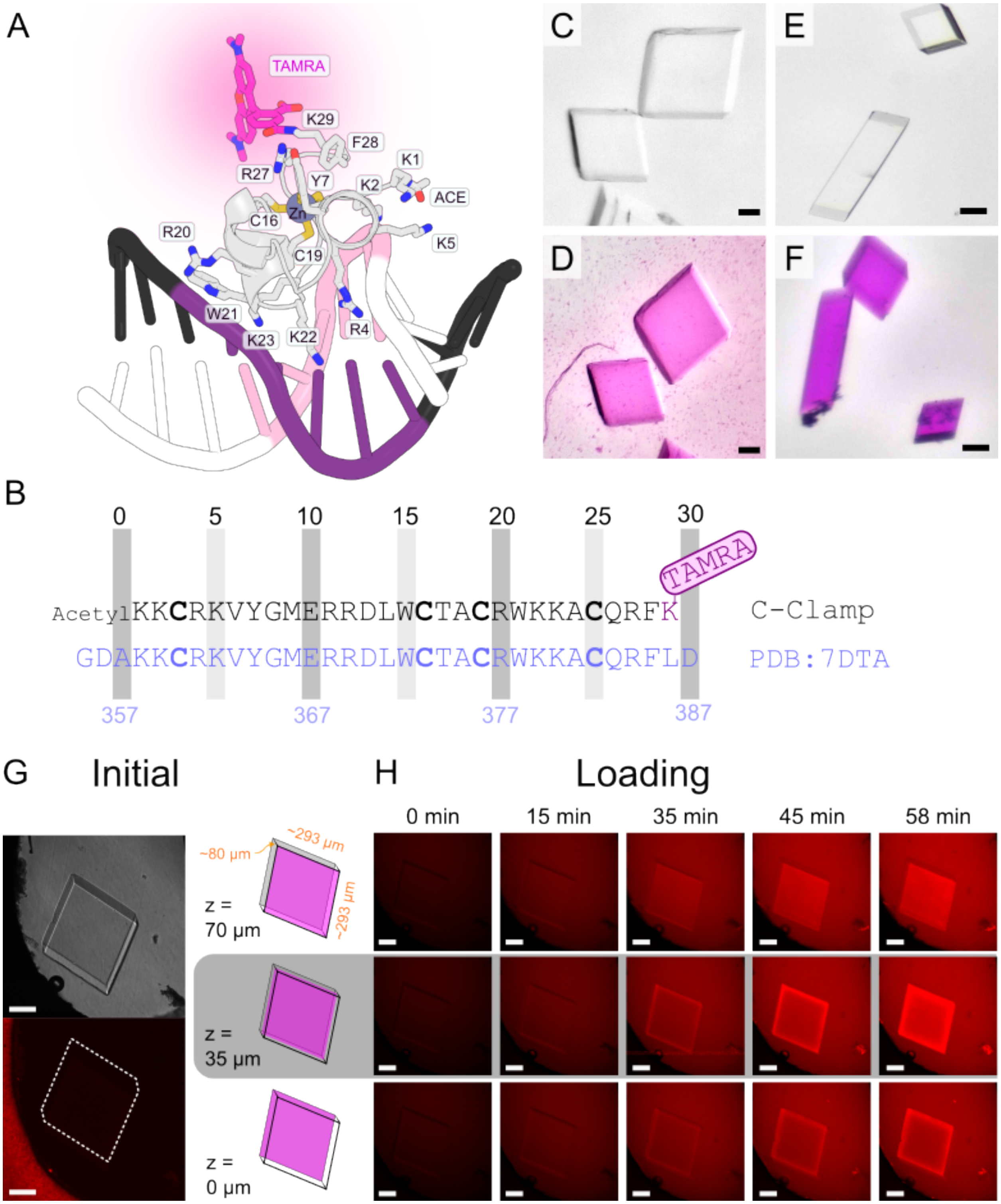
A
TAMRA
K29
F28
K1
R27
Y7
K2
ACE
Zn
C16
R20
K5
C19
W21
K23
K22
R4
C
E
D
F
B
0 5 10 15 20 25 30
TAMRA
AcetylKKCRKVYGMERRDLWCTACRWKKACQRFK
C-Clamp
GDAKKCRKVYGMERRDLWCTACRWKKACQRFLD
PDB:7DTA
357 367 377 387
G
Initial
~80 µm
~293 µm
~293 µm
z = 70 µm
z = 35 µm
z = 0 µm
H
Loading
0 min
15 min
35 min
45 min
58 min

**Figure S7. C-clamp and stereozoom microscope images of CC1$^{+10}$ adsorbing C-clamp**

(**A**) A C-clamp model was manually prepared from model 1 in the 7DTA NMR ensemble (Human HDBP1). We removed terminal residues, mutated Leu386Lys, and placed TAMRA (CCD FH8) to mimic K29 conjugation. (**B**) Sequence comparison: our C-clamp vs. PDB:7DTA. Zn-coordinating cysteines are shown in **bold**. A C-terminal Lys was added for TAMRA conjugation (magenta). Crosslinked (53 mg/mL EDC) CC1$^{+10}$ crystals (matching PDB:9YZA) **(C)** before and **(D)** after a 24 hr soak in 50 µM C-clamp-TAMRA in 200 µL of CC1 mother liquor (1.6 M Lithium sulfate, 40 mM magnesium acetate, 50 mM MES pH 6.5). Crosslinked (51.5 mg/mL EDC) CC1$^{+10}$ crystals (matching PDB:9YZA) before **(E)** and **(F)** after a 48 hr soak in 300 µM C-clamp-TAMRA in 200 µL of CC1 mother liquor (1.6 M Lithium sulfate, 40mM magnesium acetate, 50mM MES pH 6.5). (**G-H**) Early confocal microscopy imaging work (561 nm laser power 5%) with the larger crystal from panels C&D, prior to our investigation of the effects of salt concentration on guest capture, revealed C-clamp loading required more than an hour despite the small size of the guest. Notably, this moderately large crystal is near the edge of a drop which may slow transport. **(G)** Initial imaging prior to addition of C-clamp to the drop showed the parallelepiped shape with differential interference contrast imaging and negligible red fluorescence emission. **(H)** A time resolved z-stack ($\Delta z = 5$ µm, $\Delta t = 1$ min) with a schematic z-plane depiction for each row on the left.

Main text Figure 2 shows guest loading under conditions that lead to guest binding to specific DNA sequences. However, high occupancy target site binding is not required for guest uptake into the crystals. We included this C-clamp loading experiment to demonstrate this phenomenon. The C-clamp concentration for this experiment was 50 µM in a 5 µL loading solution of: 1.6 M Lithium sulfate, 40 mM magnesium acetate, and 50 mM MES pH 6.5. Under these high-salt conditions, X-ray diffraction reveals the scaffold crystal, but no evidence of site-specific C-clamp binding. Nonetheless, C-clamp uptake is clear. Strong adsorption (panels D, F, and H) *without* site-specific binding is an intriguing phenomenon that will require further study. Possibly, uptake is driven entropically where each guest protein taken into the crystal ultimately releases multiple cations to the higher-entropy bulk solvent.

Regardless of the mechanism, C-clamp adsorption to the crystal was obvious, and followed roughly the same pattern observed in main text Figure 2. Specifically, the top of the crystal adsorbed guest rapidly and uniformly, which makes sense for the top facet of the crystal having the large solvent channel apertures. An interior plane and a plane closest to the slide show bright edges and a slower guest increase in the interior, especially in the bottom plane. C-clamp should be small enough to diffuse through the lateral pores (see Fig. S11). Scale bars are 100 µm. Figure prepared in PyMOL (82) and Inkscape.

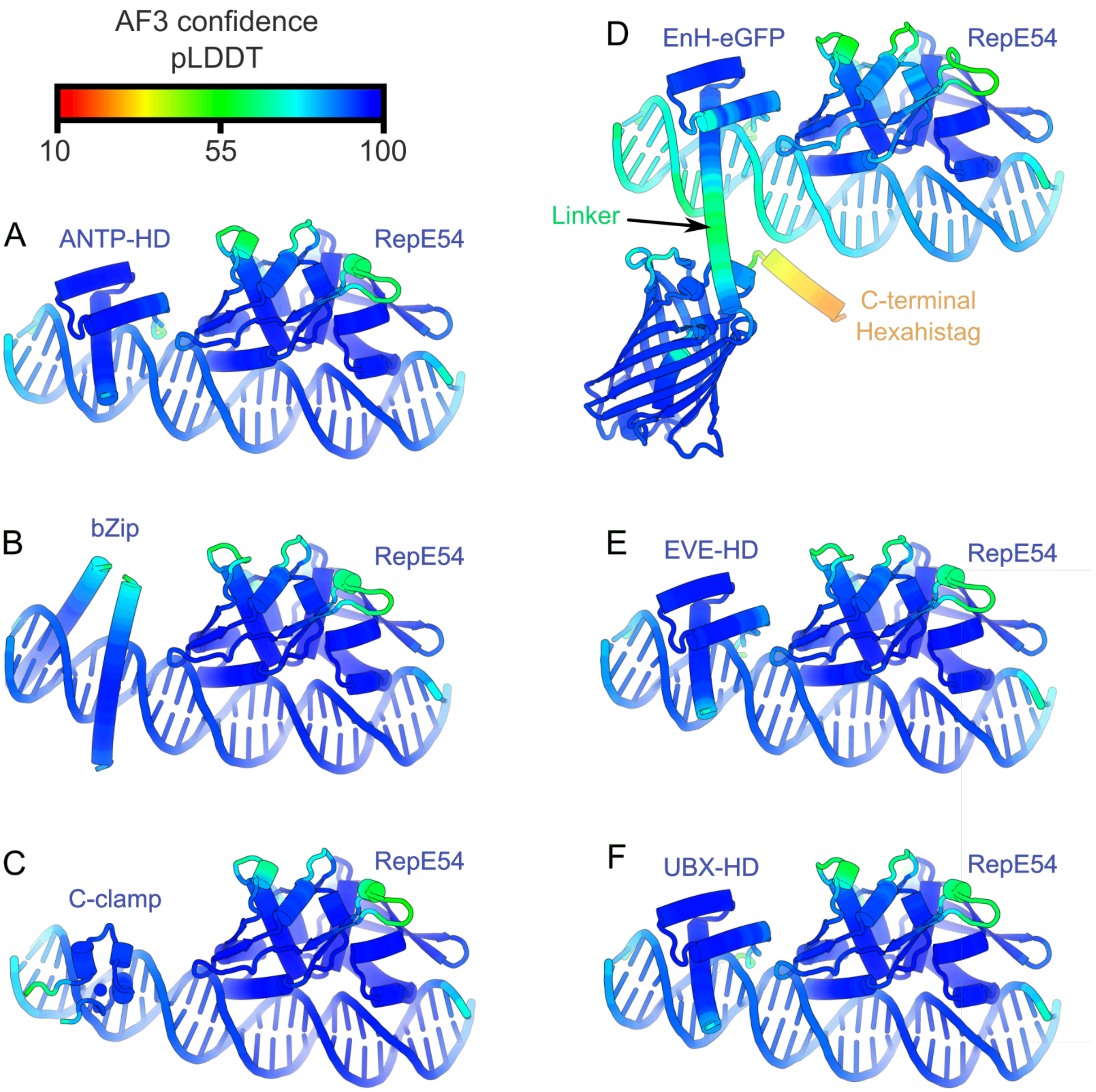
AF3 confidence
pLDDT
10
55
100
A
ANTP-HD
RepE54
B
bZip
RepE54
C
C-clamp
RepE54
D
EnH-eGFP
RepE54
Linker
C-terminal
Hexahistag
E
EVE-HD
RepE54
F
UBX-HD
RepE54

**Figure S8. AlphaFold-3 structure prediction of guest proteins bound to $CC1^{+10}$**

The models depicted here are examples of AF3 (40) predicted structures of $CC1^{+10}$ asymmetric units / building blocks colored by pLDDT. **(A)** ANTP-HD bound to $CC1^{+10}$ with insert 5' TGATGAGCAG 3'. **(B)** bZip bound to $CC1^{+10}$ with insert 5' ATGAGTCATA 3'. **(C)** C-clamp bound to $CC1^{+10}$ with insert 5' CCCGGCCGGA 3'. **(D)** EnH-eGFP bound to $CC1^{+10}$ with insert 5' TGATGAGCAG 3'. **(E)** EVE-HD bound to $CC1^{+10}$ with insert 5' TGATGAGCAG 3'. **(F)** UBX-HD bound to $CC1^{+10}$ with insert 5' TGATGAGCAG 3'. To check for compatibility with the $CC1^{+10}$ lattice, each such model was aligned to the most closely related crystal structure for empty $CC1^{+10}$. See Supplemental Methods: Alphafold3 modeling. Figure prepared in PyMOL (82) and Inkscape.

As a side note, the use of TGATGA as a recognition sequence for the homeodomains was motivated by ITC data collected by the Spratt laboratory (not shown) having been originally inspired by SOLEXA data from FlyBase (https://flybase.org) (87).

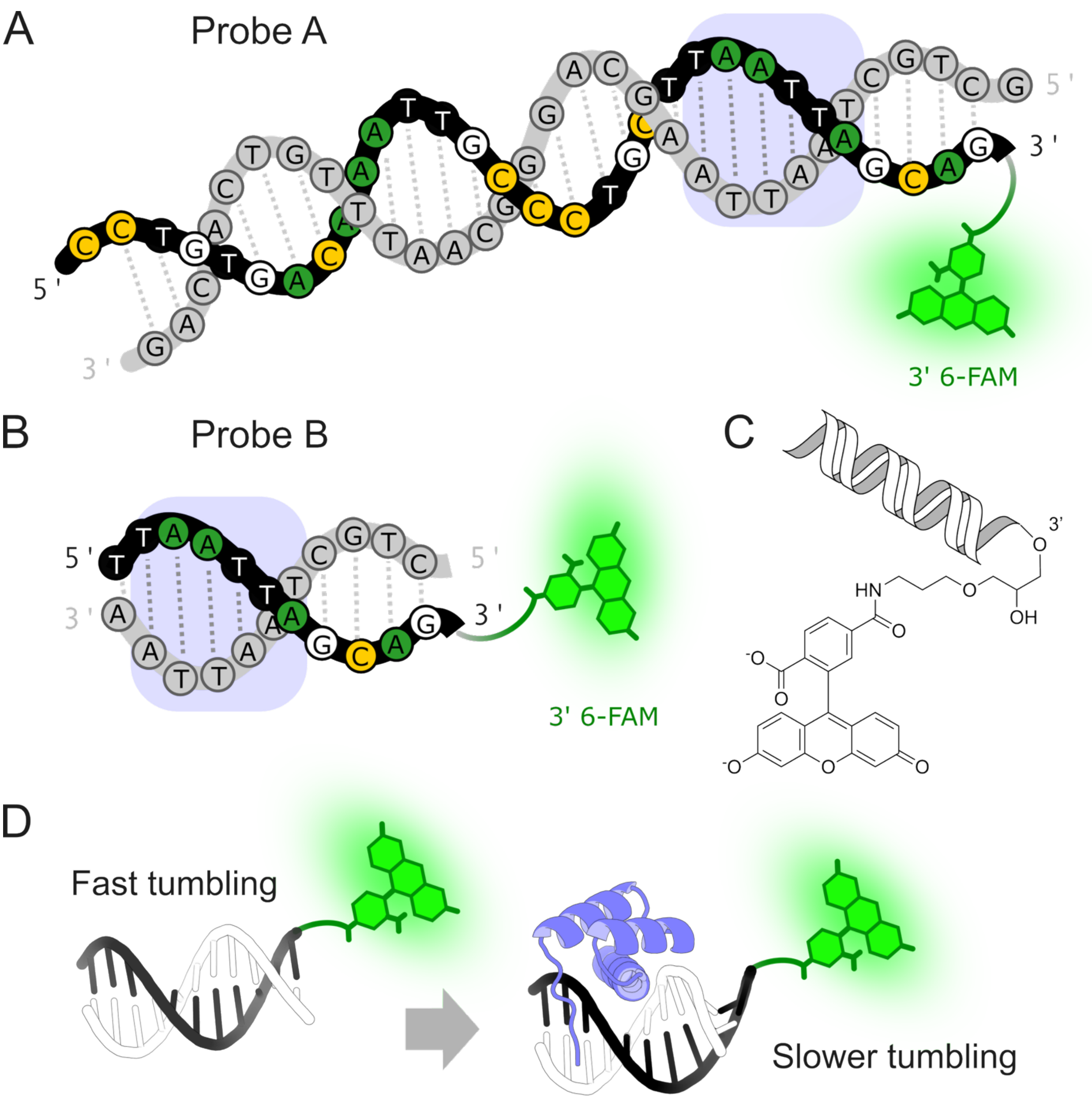

A
Probe A
5 '
3 '
3 '
5 '
3' 6-FAM
B
Probe B
5 '
3 '
5 '
3 '
3' 6-FAM
C
D
Fast tumbling
Slower tumbling

**Figure S9. Fluorescence polarization probe schematics.**

(**A**) A full-length fluorescence polarization probe A (also used for $CC1^{+10}$ crystallization trials) with a 3’ 6-FAM conjugation (Integrated DNA Technologies). The location of a canonical sequence for homeodomain binding (5’ TAATTA 3’) is marked with a blue backdrop. (**B**) Truncated fluorescence polarization probe B. **(C)** Depiction of the fluorophore chemical structure and linker to the 3’ OH of DNA **(D)** Schematic illustration of EVE-HD binding to Probe B. Figure prepared in Inkscape, ChemDraw Professional (Version 25.5.0.6237, Revvity Signals Software, Inc.), and PyMOL (82).

The sequences below show the target homeodomain site (bold & underlined). Probe A also has the 5’-AATT-3’ R1 site within the RepE54 binding footprint (bold).

Probe A:<br>
5'-CCTGTGACA**AATT**GCCCTGCT**TAATTA**GCAG/36-FAM/-3'<br>
3′-GACACTGT**TTAA**CGGGACGA**ATTAAT**CGTCG-5′

Probe B:<br>
5′-T**TAATTA**GCAG/36-FAM/-3'<br>
3′-A**ATTAAT**CGTC-5′

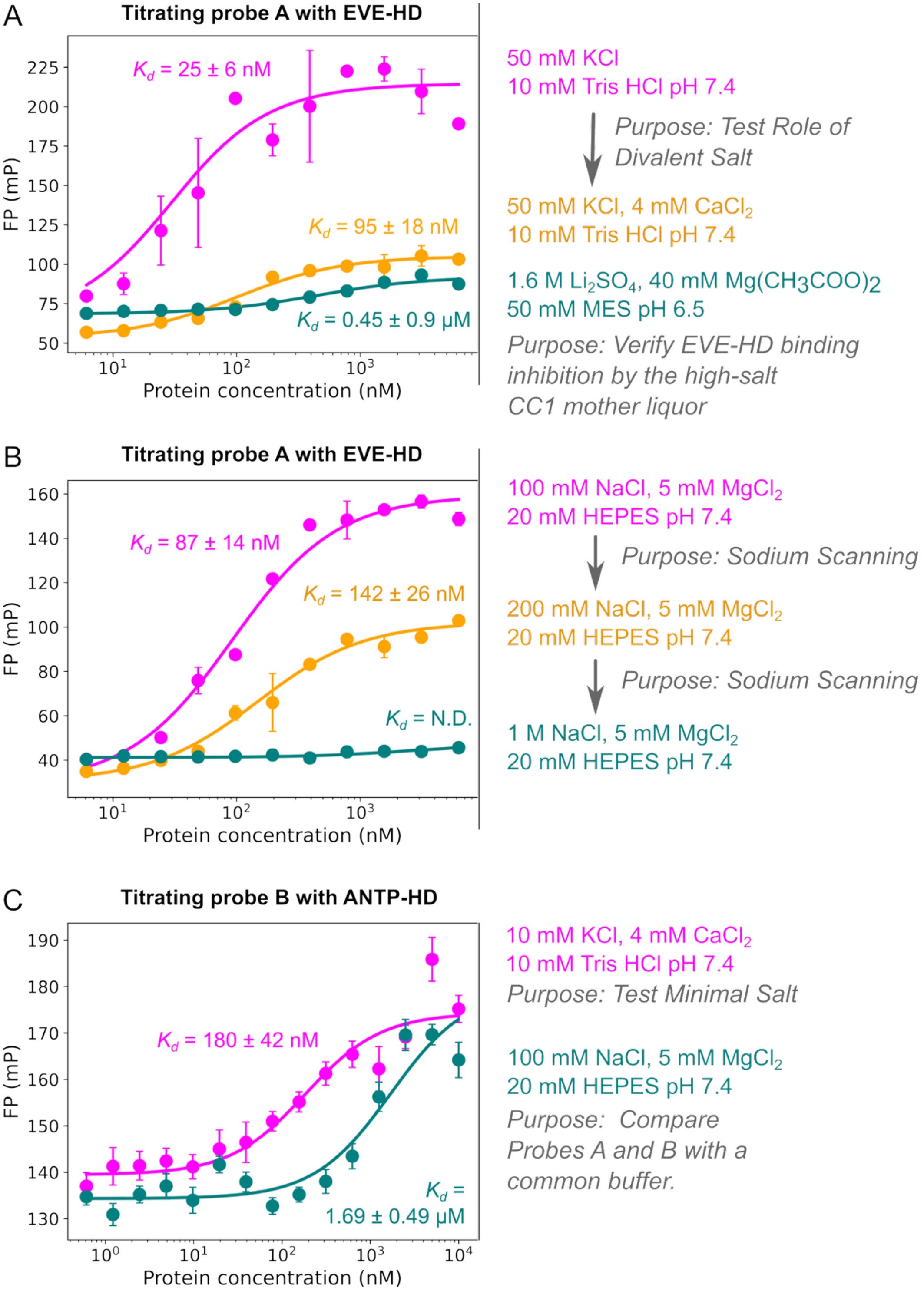


D **Morrison Quadratic Binding Model:**

$$FP = B + (T - B) \times \frac{(K_d + P_t + L_t) - \sqrt{(K_d + P_t + L_t)^2 - 4P_tL_t}}{2L_t}$$

| **Fixed parameters:** | **Fit parameters:** |
|---|---|
| *FP*: Anisotropy (mP) | *B*: Free Probe Signal (mP) |
| $P_t$: Total Protein Concentration (nM) | *T*: Bound Probe Signal (mP) |
| $L_t$: Total Probe Concentration = 10 nM | $K_d$: Dissociation Constant (nM) |

**Figure S10. Fluorescence polarization binding condition scouting data and fits**

We call these experiments "scouting" because the goal was to quickly find at least one salt regime consistent with DBD binding, rather than a systematic survey of guest binding as a function of solution conditions. Confidently ruling out (or building confidence in) alternate stoichiometry (e.g. 2:1 or 1:2 protein:DNA) would require less noisy experimental data, an effort counter to the scouting goal. Therefore, all fits used a 1:1 binding model. Each fit was suitable for semi-quantitative assessment of binding, with $R^2$ values above 0.9. (**A**) Fluorescence polarization fit of EVE-HD binding to the longer probe A (Fig. S9A) under three solvent conditions. Points are the mean of 2 replicates, and the error bars are the standard error of the mean (SEM) for better comparison with the data below. (**B**) Fluorescence polarization fit of EVE-HD binding to probe A (Fig. S9A) under three solvent conditions (each with 2 replicates) with varying sodium chloride concentrations. (**C**) Fluorescence polarization fit of a second homeodomain, ANTP-HD binding to probe B (Fig. S9B) under two solvent conditions (each with 8 replicates). (**D**) To account for depletion effects, we avoid the assumption that the total protein concentration equals the free protein concentration by using the quadratic (Morrison) equation. The Python scripts (https://github.com/cdsnow/FPfitting) fits the protein–DNA complex concentration at each titration point, incorporating the fixed final 10 nM concentration of fluorescent duplex to account for ligand depletion. The FP data can be found in the same repository. Nonlinear regression weighted by the inverse variance was used to simultaneously optimize $K_d$, the baseline polarization, and the bound-state polarization. For comparison, the scripts also perform simple Langmuir fitting. The values are quite close as shown in the tabulated results below and the conclusions drawn remain the same. Figure prepared with matplotlib and Inkscape.

| | | Quadratic $K_d$ (nM) | Langmuir $K_d$ (nM) |
|---|---|---|---|
| A | 50 mM KCl, 10 mM Tris HCl pH 7.4 | 25 ± 6 | 31 ± 6 |
| A | 50 mM KCl, 4 mM $CaCl_2$, 10 mM Tris HCl pH 7.4 | 95 ± 18 | 101 ± 19 |
| A | 1.6 M $Li_2SO_4$, 40 mM $Mg(CH_3COO)_2$, 50 mM MES pH 6.5 | 446 ± 86 | 452 ± 86 |
| B | 100 mM NaCl, 5 mM $MgCl_2$, 20 mM HEPES pH 7.4 | 87 ± 14 | 93 ± 14 |
| B | 200 mM NaCl, 5 mM $MgCl_2$, 20 mM HEPES pH 7.4 | 142 ± 26 | 148 ± 26 |
| C | 10 mM KCl, 4 mM $CaCl_2$, 10 mM Tris HCl pH 7.4 | 180 ± 42 | 185 ± 43 |
| C | 100 mM NaCl, 5 mM $MgCl_2$, 20 mM HEPES pH 7.4 | 1688 ± 488 | 1699 ± 490 |

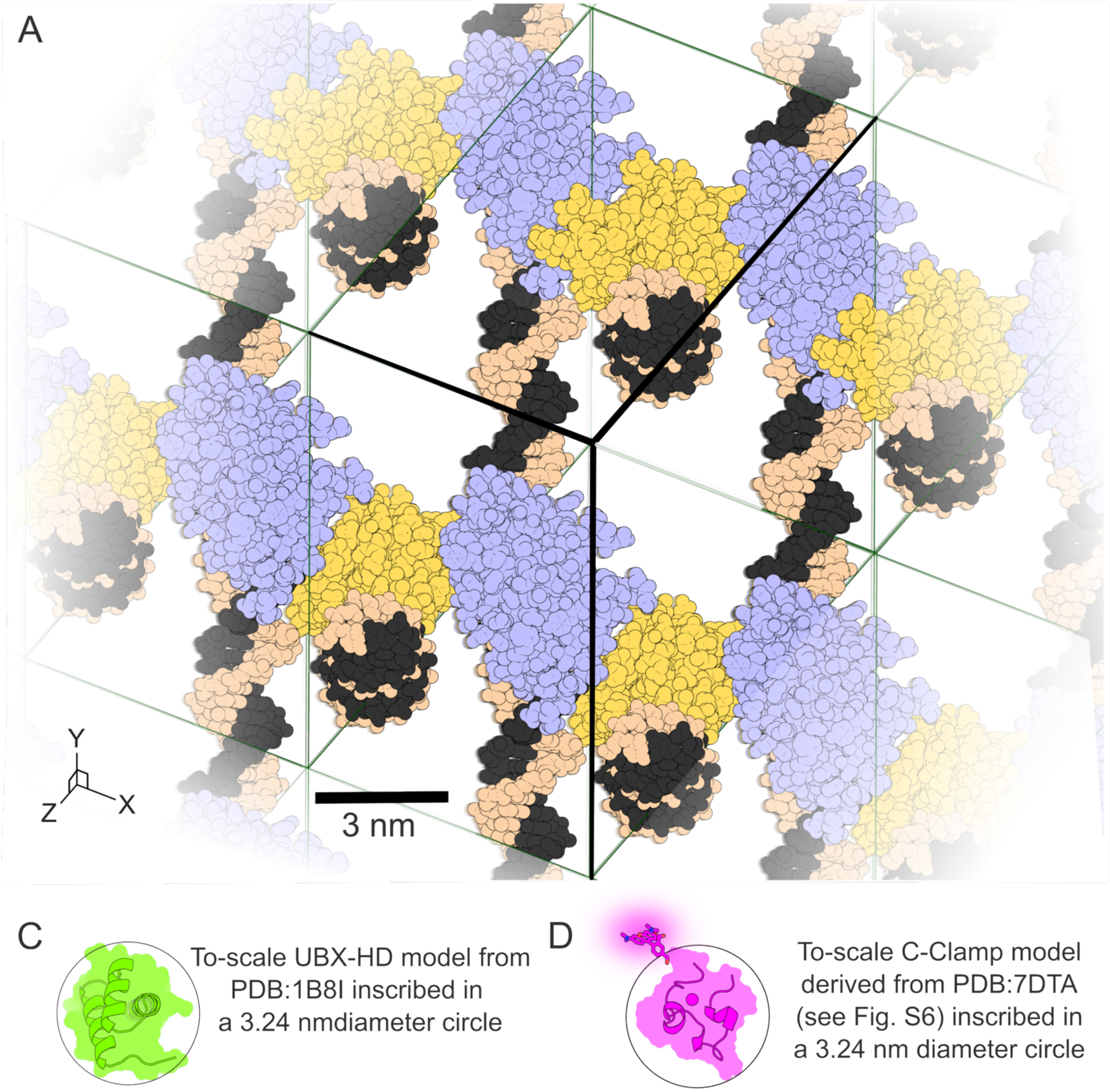


**Figure S11. Size comparison: lateral solvent channels in CC1$^{+10}$ versus homeodomain**

The clearest view of lateral solvent channels is obtained by orienting the viewpoint to look down the ***–a –b –c*** vector. **(A)** Space-filling side-view of a representative empty CC1$^{+10}$ (PDB:9YZJ) with a 3 nm scale bar and neighboring unit cells shading white. DNA-chains (black and tan) are oriented vertically or perpendicular to the screen. RepE54 monomers are colored alternately light blue or yellow. Notably, MAP_CHANNELS (67) analysis predicts that guests smaller than 3.24 nm diameter could diffuse through this crystal via channels other than the major 5 nm pores. **(B)** The 3.24 nm diameter limit is close to the size of homeodomains. Here we show that UBX-HD (PDB:1B8I), the protein used for the confocal loading experiment in main text Fig. 2, is comparable in size to a 3.24 nm circle. We hypothesize that lateral diffusion contributes to the Fig. 2 UBX-HD loading pattern, albeit at transport rates slower than diffusion along the major solvent channels. **(C)** The 3.24 nm limit should also be large enough for C-clamp diffusion. A C-clamp model (Fig. S6) is pictured here inscribed within a 3.24 nm circle (showing one possible placement for the flexible C-terminal TAMRA fluorophore tag). Supporting this assessment, confocal data for C-clamp loading (Fig. S6) is consistent with lateral diffusion that is slower than uptake via the major pores. Figure prepared in PyMOL (82) and Inkscape.

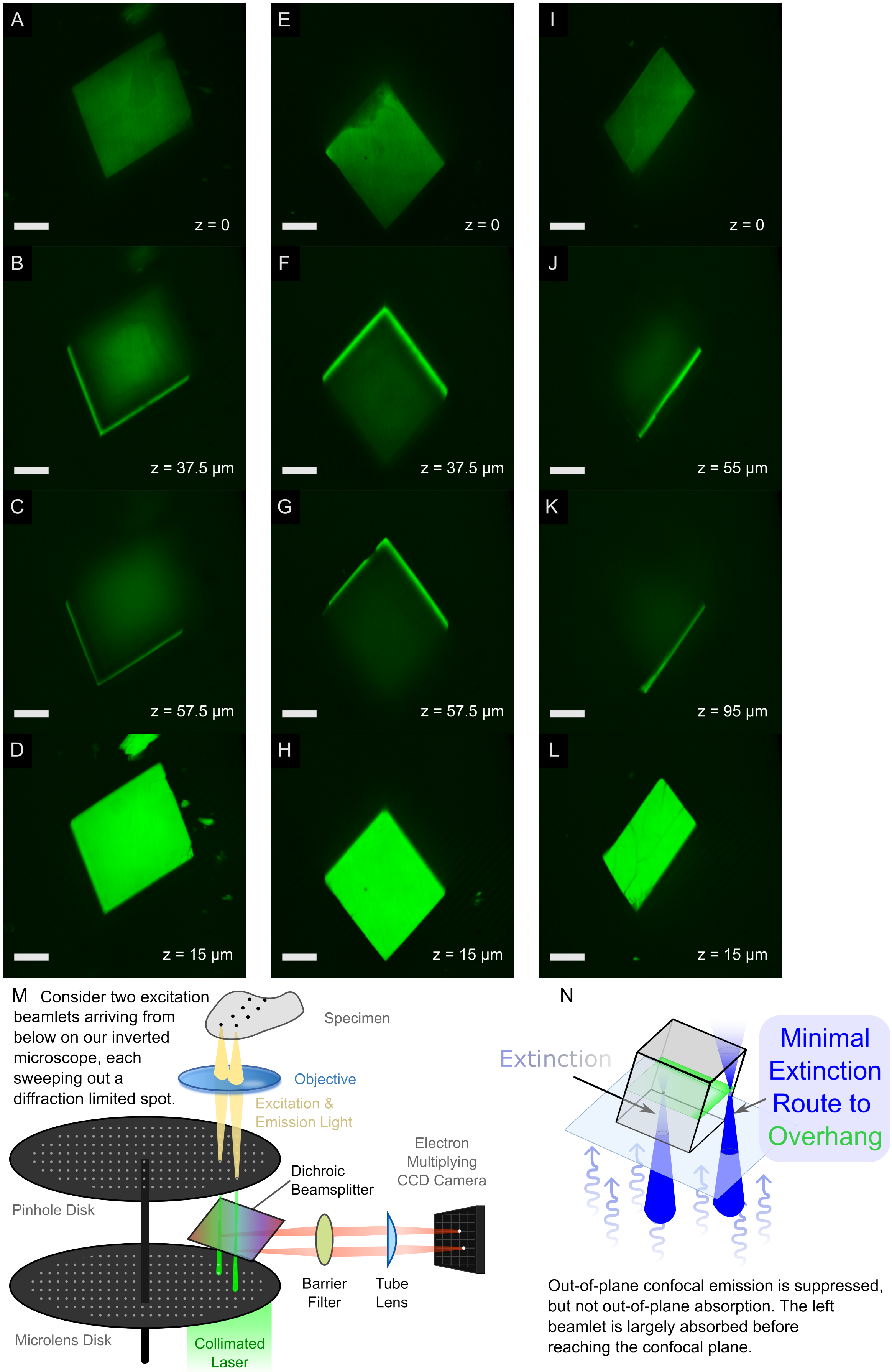
A
z = 0
B
z = 37.5 μm
C
z = 57.5 μm
D
z = 15 μm
E
z = 0
F
z = 37.5 μm
G
z = 57.5 μm
H
z = 15 μm
I
z = 0
J
z = 55 μm
K
z = 95 μm
L
z = 15 μm
M
Consider two excitation beamlets arriving from below on our inverted microscope, each sweeping out a diffraction limited spot.
Specimen
Objective
Excitation & Emission Light
Electron Multiplying CCD Camera
Dichroic Beamsplitter
Pinhole Disk
Barrier Filter
Tube Lens
Microlens Disk
Collimated Laser
N
Extinction
Minimal Extinction Route to Overhang
Out-of-plane confocal emission is suppressed, but not out-of-plane absorption. The left beamlet is largely absorbed before reaching the confocal plane.

**Figure S12. Multiple confocal views of the same $CC1^{+10}$ crystal loaded with UBX-HD trace-labeled with NHS-fluorescein**

To rule out heterogenous loading of guest protein we obtained z-stack confocal images for multiple orientations of the same crystal. Scale bars are 50 µm. (**A–D**) Optical sections of a $CC1^{+10}$ crystal (same crystal as Fig. S13 and Fig. S14) loaded with 80 µM UBX-HD trace-labeled with NHS-fluorescein. (**A**) Bottom z-slice closest to the microscope slide. (**B**) Mid-plane (z=37.5 µm) showing reduced fluorescence emission in the crystal interior, but strong emission from the two overhanging edges (lacking intervening crystal to absorb the excitation). (**C**) Z-slice (z=57.5 µm) that is even farther from the slide showing more pronounced extinction in the crystal interior. (**D**) Z-slice 10 µm above the bottom face corresponding to the brightest plane within the z-stack. **(E–H**) and (**I–L**), the same type of z-slices after flipping the crystal 180° and 70° respectively (see Fig.S14). (**M**) Schematic representation of our Yokagawa CSU-X1 spinning disc spinning disk confocal, with beamlets entering the sample from below. (**N**) Schematic representation of a parallelepiped crystal resting on a microscope slide, showing how the excitation light path length through crystalline material varies depending on the location, with overhanging edges experiencing minimal extinction.

These different views of the same crystal provide strong evidence that the striking observed fluorescence emission z-stack patterns were primarily due to variable excitation extinction. In panel **A**, a small defect notch in the crystal is visible just to the right of the top corner. The brightest plane in the crystal (panel **D**) was 10 microns above this defect. After the crystal was flipped 180°, this same small defect is now visible in panel **G**, just to the left of the top corner at a new height of 42.5 microns above the slide. In the new orientation, the notch is part of an overhang and is therefore illuminated despite most of the crystal being dim. In the second orientation, the bright uniform layer (panel **H**) was again 10 microns above the slide. This layer was 47.5 microns above the slide in the original orientation (similar to panel **C**) and was therefore mostly dim other than the two overhanging edges. *Thus, the bright zones in the crystal were dominated by proximity to unimpeded illumination rather than heterogeneous loading.*

As a final proof, the crystal was placed onto one of the smaller planes s. Again, the bright z-plane was parallel to the slide and 10 microns above. In this orientation, only one facet had a visible slight overhang. No matter the crystal orientation, the well-illuminated z-planes (without visible extinction effects near the slide) appeared to be uniformly loaded.

This crystal differs from the main text Fig. 2 crystal in two critical ways. First, the crystal here is larger (maximum side length of ~293 µm rather than ~81 µm). Second, for this figure we were loading with the trace labeled UBX-HD sample (the reaction product from a 1:5 dye:protein conjugation stoichiometry), but the result was an overly high fluorophore concentration within the crystals leading to overly prominent extinction effects. In contrast, the trace labeled UBX-HD in Fig. 2 was further diluted 1:200 with unlabeled UBX-HD.

When loading fluorescent guest molecules that adsorb to a host crystal, the photophysical effects of reaching a high local concentration inside the crystals may become evident. There are 4 copies of the asymmetric unit within each I121 unit cell for the $CC1^{+10}$ host crystals, and the unit cell volumes are 1186-1351 $nm^3$. Thus, installing one guest per asymmetric unit would lead to an intra-crystal local concentration of 4.9-5.6 mM (and we typically observe two guest homeodomains per asymmetric unit for an implied concentration of ~10 mM plus any additional guests that are inside the crystal but unbound). We hypothesize that in future work it might be possible to use the extinction effect to calculate the concentration of fluorophore within the crystal. Specifically, one could quantitatively analyze the optical properties of this system and using the brightly lit edges as an internal control, fit the internal concentration that can predict intensity versus depth in the crystal using Beer's law and known properties of the fluorophore. To enable downstream analysis, we deposited our confocal data into a Zenodo repository: 10.5281/zenodo.18328922. Figure prepared with NIS Elements, FIJI, and Inkscape (panels **M-N)**.

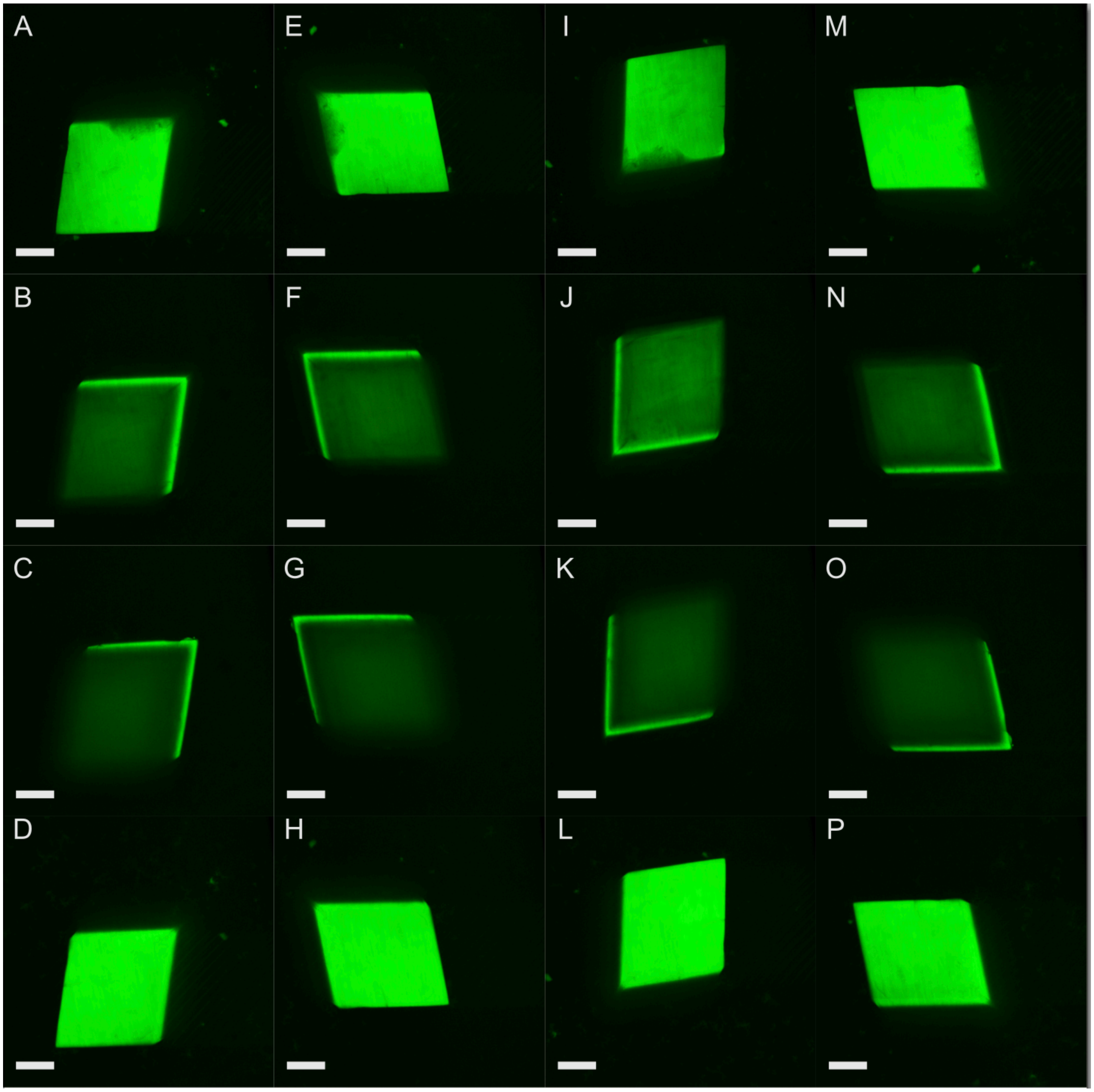


**Figure S13. CC1$^{+10}$ confocal loading of UBX-HD trace-labeled with NHS-fluorescein**

Our inverted confocal microscope (Nikon Eclipse Ti spinning-disk confocal microscope with a Yokagawa CSU-X1 quad dichroic 10kRPM spinning disc) illuminates samples from below. Crystals with high internal fluorophore concentrations can exhibit extinction effects (Fig S12) where emission intensity is reduced due to excitation absorption by intervening layers. Facets with overhangs were bright regardless of the rotation. (**A–D**) Optical sections of a CC1$^{+10}$ crystal (identical DNA sequence to 9YZF.pdb) loaded with 80 μM UBX-HD trace-labeled with NHS-fluorescein. (**A**) Bottom optical section. (**B**) Mid-plane section showing reduced fluorescence emission in the crystal interior, but strong emission from the two overhanging edges (lacking intervening crystal to absorb the excitation). (**C**) Z-slice (z=45 μm) that is even farther from the microscope slide showing more pronounced extinction in the crystal interior. (**D**) Z-slice 10 μm above the bottom face corresponding to the brightest plane within the z-stack. (**E–H**), (**I–L**), and (**M-P**) To rule out crystal orientation edge illumination effects, we collected z-stacks after rotating the crystal 90°, 180°, and 270° counter-clockwise around an axis perpendicular to the slide. Scale bars are 50 microns. Figure prepared with NIS Elements, FIJI, and Inkscape.

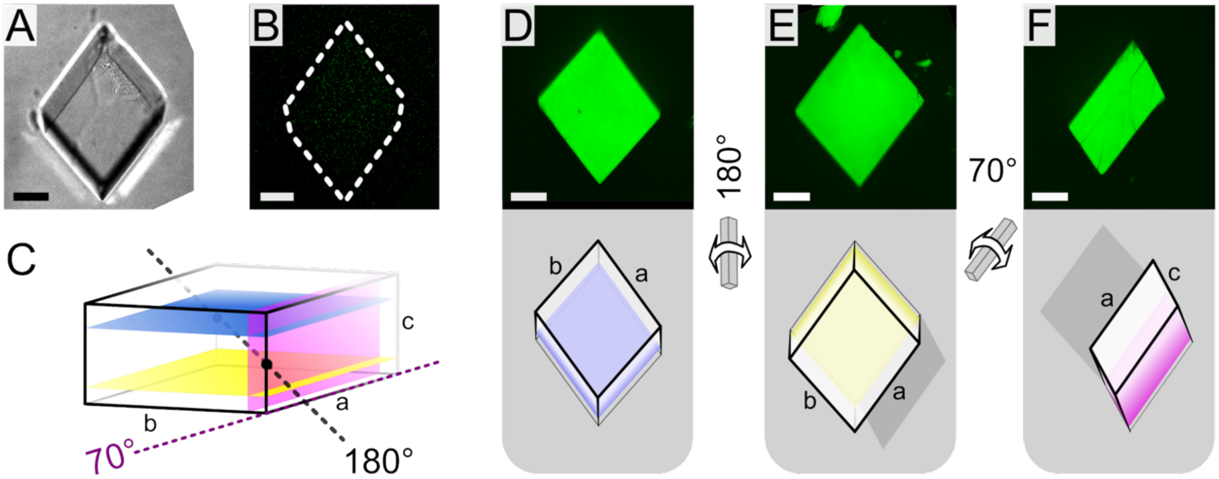


**Figure S14. Confocal Microscopy Images of CC1$^{+10}$ crystal loaded with UBX-HD trace-labeled with NHS-fluorescein**

This figure summarizes the confocal data shown in Fig. S12. **(A)** Unloaded CC1$^{+10}$ crystal with asymmetric insert 5'TAATTAGGCCG 3' via differential interference contrast. **(B)** Unloaded CC1$^{+10}$ crystal with asymmetric insert 5'TAATTAGGCCG 3' under 488nm laser. **(C)** Our crystal measurements gave a triclinic crystal shape of a = 152 µm, b = 136 µm, c = 73µm, α = 66°, β = 63°, γ = 78°. Schematic representation of the same CC1$^{+10}$ crystal, depicting 3 internal planes imaged in subsequent panels. **(D)** Highest fluorescence intensity plane and schematic of one crystal orientation. **(E)** Highest fluorescence intensity plane after a 180° crystal flip. (**F**) Highest fluorescence intensity plane and schematic of the same crystal resting on a smaller side facet. Scalebars are 50 microns. Figure prepared with FIJI (88) and Inkscape.

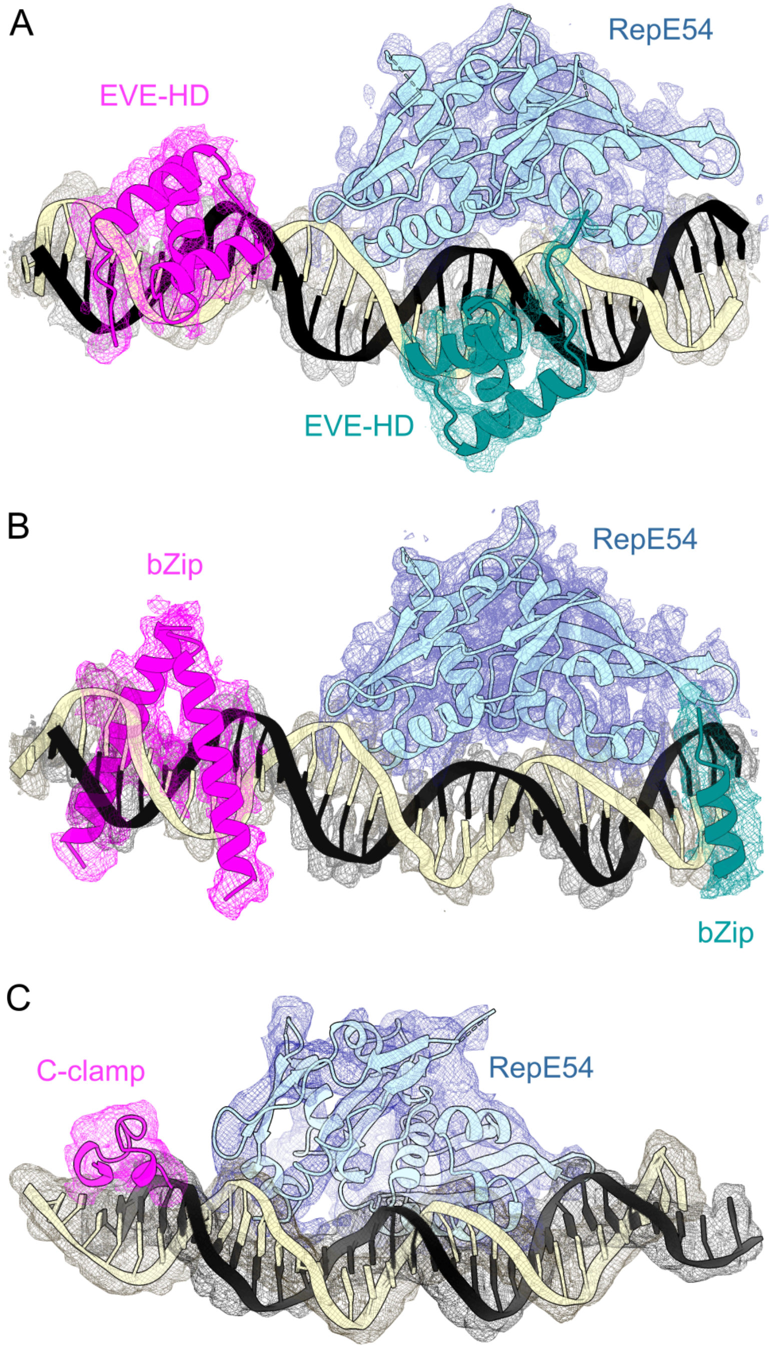


**Figure S15. Representative PHENIX omit maps of each guest structure motif**

Each panel is a PHENIX simulated annealing omit map (46). **(A)** 9YZM.pdb with omitted EVE-HD shown in magenta & teal. **(B)** 9YZU.pdb with omitted bZip shown in magenta & teal. **(C)** 9Z44.pdb with omitted C-clamp shown in magenta. All meshes individually contoured at 1.0 using ChimeraX (sdLevel 1.0). Omit maps of this type are the standard method to verify that electron density map features are not the result of model bias (45–47). Omit maps confirmed that the guest molecule density was well supported by the data. Figure prepared with scripted ChimeraX (89) and Inkscape.

Notably, “Bonus density” was not unique to the homeodomain family. For the bZip guest, which interacts exclusively with DNA major grooves (90), we likewise observed additional electron density at an unanticipated register (with no notable overlap with the canonical bZip recognition sequence), although in this case the canonical target site was better resolved (Fig. 3E). The partial density was consistent with a bZip helix segment fragment but lacked resolvable side chains and was not obviously stabilized by neighboring lattice contacts. We hypothesize that this site is a low-affinity complex that was only occupied due to mass action (during loading the initial bZip concentration outside the crystal was 33 μM in a 200 μL drop).

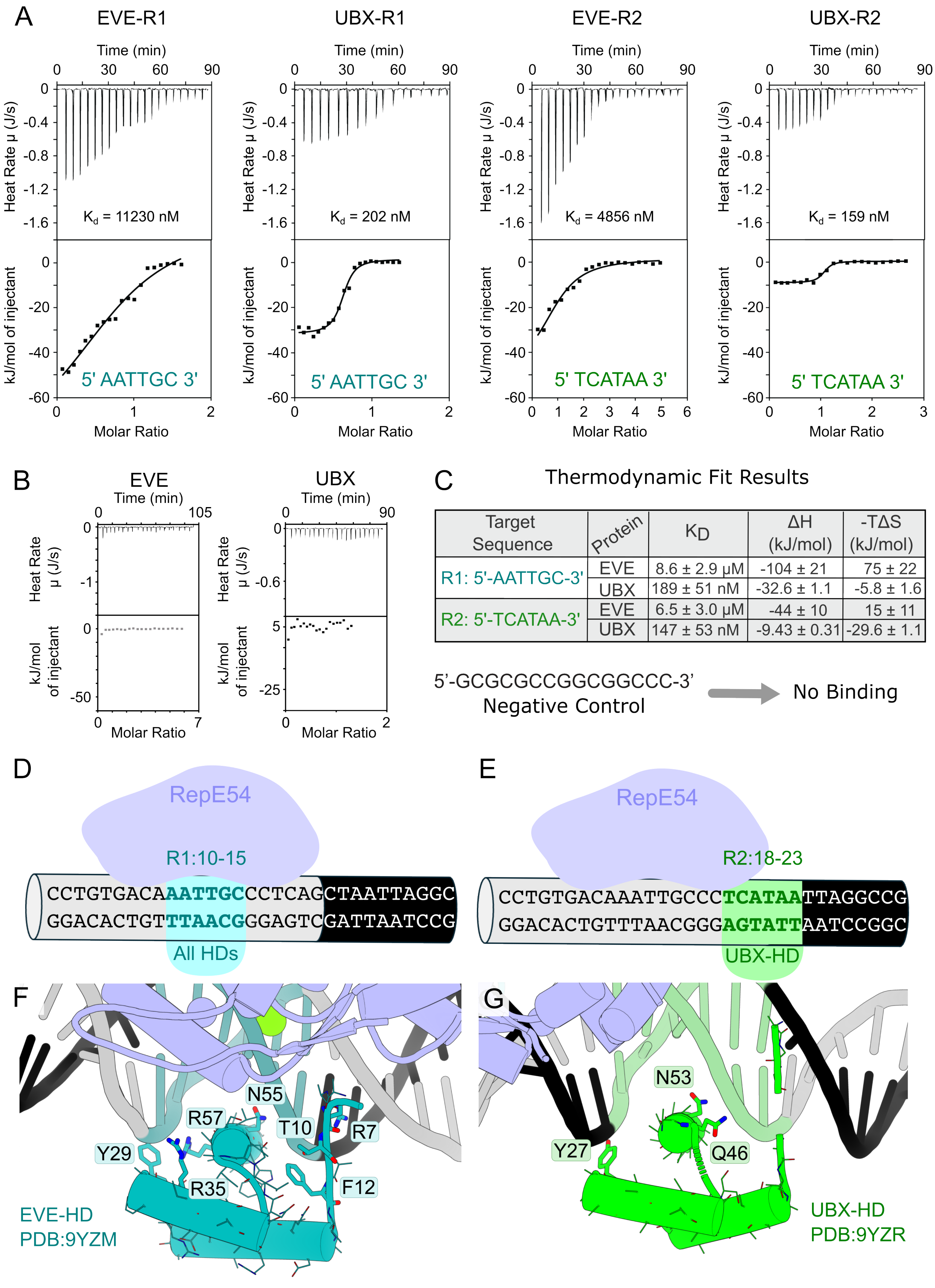
A
EVE-R1
UBX-R1
EVE-R2
UBX-R2
Time (min)
Heat Rate μ (J/s)
kJ/mol of injectant
Molar Ratio
Kd = 11230 nM
Kd = 202 nM
Kd = 4856 nM
Kd = 159 nM
5' AATTGC 3'
5' AATTGC 3'
5' TCATAA 3'
5' TCATAA 3'
B
EVE
UBX
Time (min)
Heat Rate μ (J/s)
kJ/mol of injectant
Molar Ratio
C
Thermodynamic Fit Results
Target Sequence | Protein | KD | ΔH (kJ/mol) | -TΔS (kJ/mol)
R1: 5'-AATTGC-3' | EVE | 8.6 ± 2.9 μM | -104 ± 21 | 75 ± 22
UBX | 189 ± 51 nM | -32.6 ± 1.1 | -5.8 ± 1.6
R2: 5'-TCATAA-3' | EVE | 6.5 ± 3.0 μM | -44 ± 10 | 15 ± 11
UBX | 147 ± 53 nM | -9.43 ± 0.31 | -29.6 ± 1.1
5'-GCGCGCCGGCGGCCC-3'
Negative Control
No Binding
D
RepE54
R1:10-15
CCTGTGACAAATTGCCCTCAGCTAATTAGGC
GGACACTGTTTAACGGGAGTCGATTAATCCG
All HDs
E
RepE54
R2:18-23
CCTGTGACAAATTGCCCTCATAATTAGGCCG
GGACACTGTTTAACGGGAGTATTAATCCGGC
UBX-HD
F
N55
R57
T10
R7
Y29
R35
F12
EVE-HD
PDB:9YZM
G
N53
Y27
Q46
UBX-HD
PDB:9YZR

**Figure S16. ITC measurements of non-canonical homeodomain binding sites and structures thereof** (**A)** Isothermal titration calorimetry (ITC) results of EVE-HD and UBX-HD binding to R1 (5' GCGCGCAATTGCGCCC 3') and R2 (5' GCGCGCTCATAAGCCC 3') sequences. **(B)** ITC results with a negative control duplex. **(C)** Fits for thermodynamic parameters were obtained with Python scripts (see Supplementary Methods and https://github.com/cdsnow/ITCfitting). (**D**) Schematic representation of R1 binding site along the full CC1$^{+10}$ DNA sequence from 9YZM.pdb and a PyMOL illustration of EVE-HD bound to the R1 site (teal). (**E**) Schematic representation of R2 binding site along the full CC1$^{+10}$ DNA sequence from 9YZR.pdb and a PyMOL illustration of UBX-HD bound to the R2 site (green). Figure prepared in Excel, Inkscape, and PyMOL (82). **(F-G)** Close views of the corresponding PDB models 9YZM and 9YZR.

While UBX-HD has previously been reported to have significant affinity for non-canonical target sequences (91,92)(92), it is nonetheless remarkable that UBX-HD preferred the R2 site rather than the intended target site R:21-26. In this case, the target and off target sites overlap by 3 bp, and we predict that binding is mutually exclusive due to steric clashes. Therefore, R2 wins a binding competition over the target site. This is striking because some of the candidate driving forces identified for R1 binding (main text Discussion: Scaffold Binding Driving Forces) do not apply to UBX-HD binding R2. There are no obvious helpful lattice contacts, and the N-terminus is not placed to take advantage of RepE54-induced minor groove compression depicted in Fig. S21.

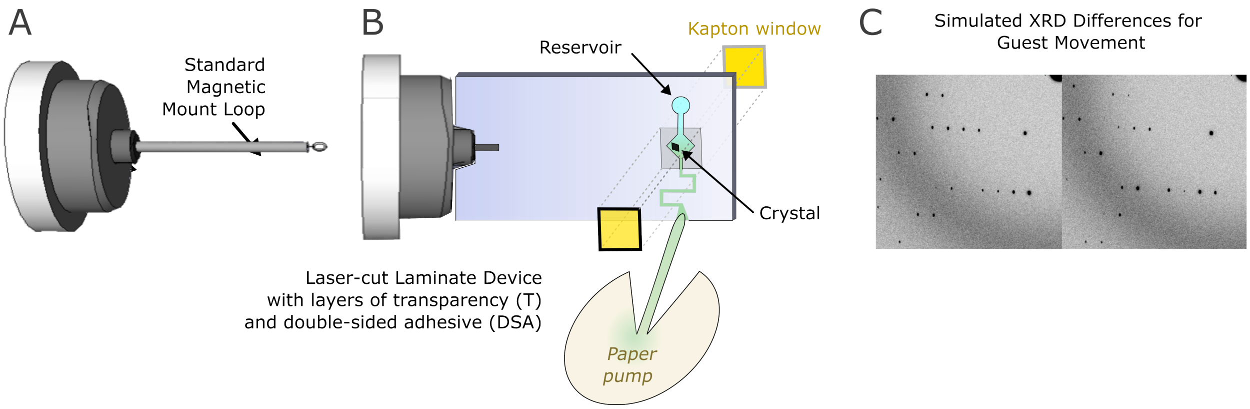


**Figure S17. Future Non-Frozen (Room Temperature) X-ray Diffraction Opportunities**

The demonstrated ability to sequentially (i) grow optimized scaffold crystals, (ii) render them stable via chemical ligation, and (iii) change solution condition while preserving X-ray diffraction opens the door to new methods such as repeatedly changing solution conditions or guest molecule titration. For example, rather than using a standard cryo-loop **(A)** or inserting such a loop into a room-temperature X-ray diffraction sleeve, simple and economical laminate microfluidic devices can be fabricated using double-sided adhesive and transparency (93). As shown in the device schematic **(B)**, one or more host crystals could be placed into a central well (open for pipetting and looping) before the front and back are closed with Kapton film (yellow) a thin transparent polymer with minimal X-ray scattering (94). A reservoir port would be open on one side for users to pipette new solution conditions (cyan). To slowly and gently pump the old solution conditions (green) out of the microfluidic device without disturbing the crystal(s) an open side port could allow insertion of a paper pump (filter paper that steadily adsorbs fluid via capillary action). If necessary, a low-melt agarose gel in the crystal chamber could help prevent crystal motion while still permitting timely solution exchange steps. Ultimately, multiple crystals could be immobilized within devices of this type allowing switching to new crystals when radiation damage accumulates. **(C)** Solution changes that drive organized movement of guest macromolecules within porous crystal scaffolds should result in observable changes to the XRD intensities. We illustrate this with simulated XRD from the MLFSOM program (95). In this *in silico* proof of concept test, adoption of a new coherent guest GFP position within a static porous crystal resulted in clear visible intensity changes to the Bragg reflections.

We hypothesize that XRD changes could be used to monitor guest DBD binding isotherms via XRD or to identify solution conditions that trigger significant movements of guests within the crystal.

There are literature precedents for these ideas, from research into fragment screening, serial crystallography, and time-resolved X-ray free electron laser strategies (96,97). Gicquel *et al*. (98) and Junius *et al*. (99) described the use of low background devices for *in situ* crystal X-ray diffraction to solve structures for crystals grown in chip. In contrast, Lyubimov *et al*. captured pre-existing microcrystals at fixed addressable locations within a trap array (100). Other work has emphasized the ability of microfluidic platforms to probe ligand soaking, buffer exchange, and weak binding events through controlled fluid delivery and room-temperature diffraction readout. For example, Sui *et al*. used capillary-based microfluidics to accelerate small-molecule binder screening via *in situ* soaking (101). Maeka used another microfluidic crystal array device to determine 8 trypsin-ligand complex structures (102).

In summary, microfluidic devices permit continuous exposure of immobilized crystals to ligand solutions, enabling rapid preparation of protein–ligand complexes and facilitating quantitative electron-density detection of weak binders in high-throughput screening formats. These reports reinforce the feasibility of dynamically perturbing solution conditions while directly observing diffraction consequences. Our scaffold crystal technology allows macromolecular guest uptake. Combining these methods with the macromolecular guest soaking enabled by our scaffold crystal technology should enable interrogation of the status of guest macromolecules (e.g. disordered, coherent) throughout the crystal interior in near real-time. Critically, it should also be possible to use small molecule fragment screening in a more traditional way, observing how different drug candidates interact with the DNA and/or the DNA-binding protein (103). For example, since the DBD is free to leave, drug discovery screening could trigger DBD dissociation while leaving the host crystal intact. Alternately, for one or more DBD installation registers we could observe how the addition of small molecules changes the DBD shape, a capacity that relies on the lack of constraints on DBD orientation in the crystal beyond its binding interface with the cognate DNA. Figure prepared in Inkscape.

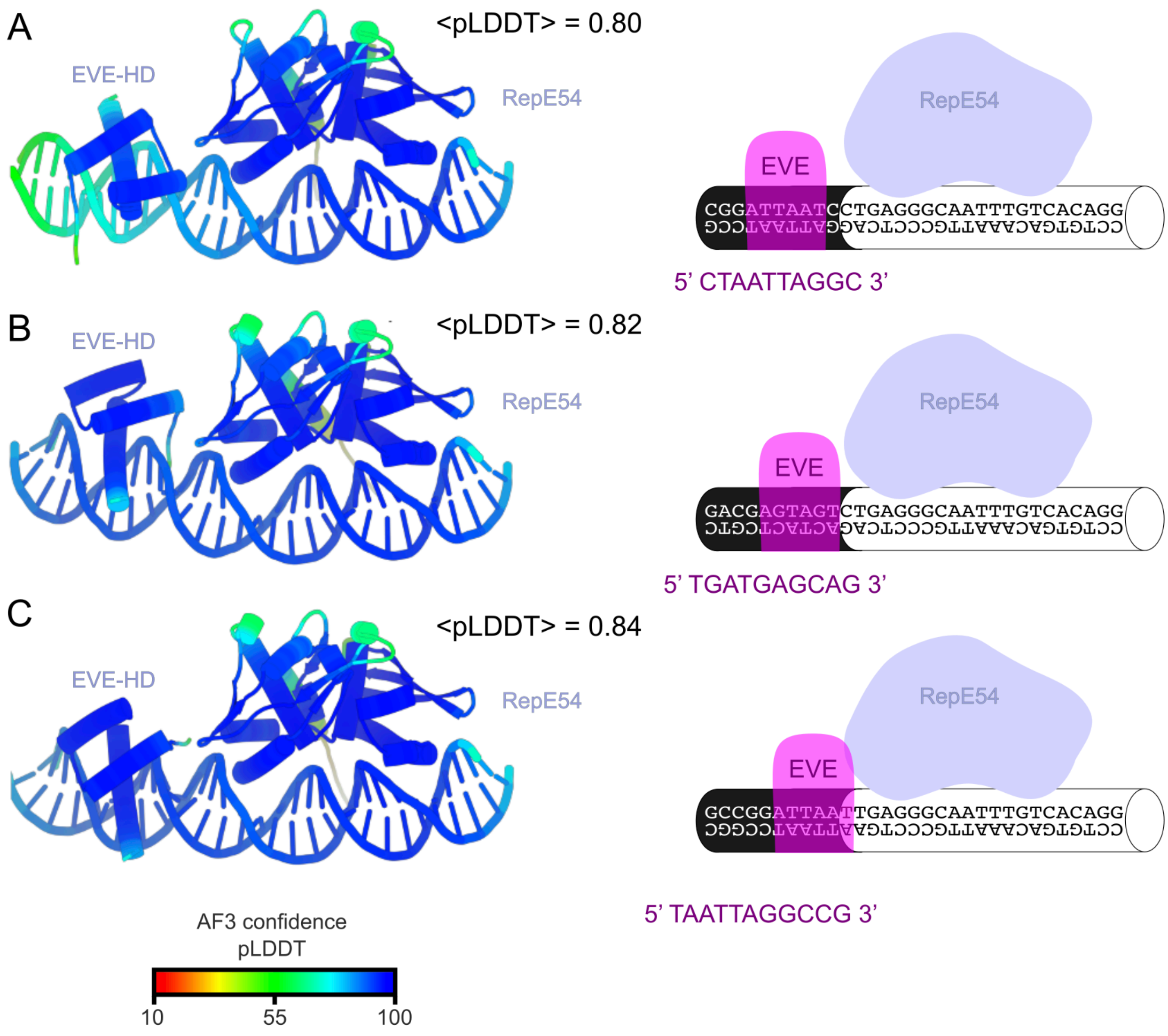


**Figure S18. AlphaFold-3 prediction of EVE-HD bound to $CC1^{+10}$ with shifting binding sites**

All depicted models are AF3 predictions colored by pLDDT. We report the mean pLDDT for the complete model. (**A**) $CC1^{+10}$ with insert 5’ CTAATTAGGC 3’ and EVE-HD. (**B**) $CC1^{+10}$ with insert 5’ TGATGAGCAG 3’ and EVE-HD. (**C**) $CC1^{+10}$ with insert 5’ TAATTAGGCCG 3’ and EVE-HD. *These predictions correspond to the crystal structures we obtained in Fig. 4*. A prediction-vs-experiment superposition is depicted in Fig. S27. Figure prepared in PyMOL (82) and Inkscape.

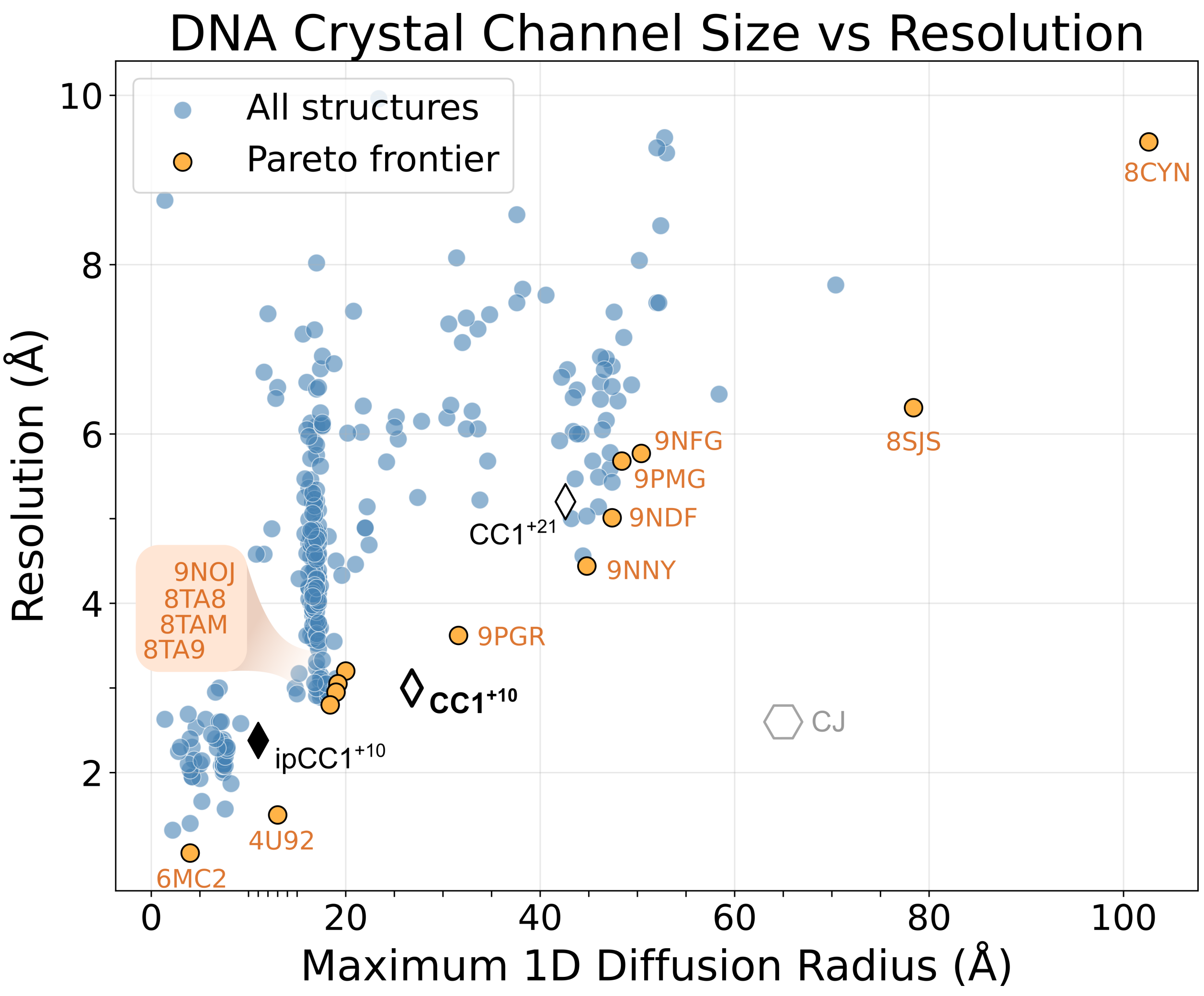


**Figure S19. Trends in Pore Size and XRD Resolution Limit**

For this meta-analysis, we collected 302 PDB entries via manual search of crystals deposited from laboratories known to work on designed DNA crystals. Crystals at the Pareto frontier (for high MAP_CHANNELs guest diffusion radius and low diffraction resolution limit) were inspected to ensure pertinence. In principle, guests with 20.5 nm diameter could diffuse into the remarkable 8CYN lattice. That structure also features long accessible DNA struts suitable for guests that seek B-DNA. This is also largely true of 8SJS, another $P6_3$ crystals from Sha and coworkers (104). 8SJS should permit diffusion for guests up to 15.7 nm diameter. At the other end of the plot, like our earlier interpenetrating ipCC1$^{+10}$ crystals wherein we could observe the small molecule netropsin binding (35), several pure DNA crystals have yielded small molecule guest structures (netropsin, Hoechst, and DAPI in 8TAB, 8TAM, and 8TA9, respectively) (105). The highest-resolution DNA crystals captured in this survey feature quadruplexes (and small solvent channels). Note: the solvent channel calculation for 4U92 is not correct since that model excludes a number of disordered base pairs (106).

Suitability as a programmable scaffold depends on properties beyond pore size and diffraction limit. The freedom to encode arbitrary guest installation sequences will depend on the size of the dsDNA regions that are sterically accessible and not otherwise constrained by symmetry considerations or proximal DNA crossovers. For example, our asymmetric $CC1^{+10}$ building blocks have the advantage compared to our symmetric $CC1^{+10}$ building blocks in that the asymmetric design places the nick sites between stacked dsDNA far from the center of the accessible struts. Nicks placed in the middle of the ideal guest binding struts may not be an issue given continued progress towards high-yield in-crystal ligation of those sites.

There is an obvious trend: more porous crystals tend to have inferior X-ray diffraction resolution. This is typical for macromolecular crystals (107–109). It will be intriguing to discover how the diffraction quality for these scaffold crystals (co-crystals and DNA crystals alike) changes with further optimizations, and to assess the effect of guest installation on the diffraction limit. For example, the rigidity and resolution limit for $CC1^{+21}$ crystals (8.5 nm diameter pores, 90% solvent) may improve upon installation of large, well-ordered guest proteins. To underscore the feasibility of reaching sub-3Å diffraction for highly porous crystals, we show an outgroup example (open hexagon symbol); the best diffraction limit we have obtained in our “CJ” porous protein crystal system (P622: PDB:5W17) is 2.6Å despite 13 nm diameter pores.

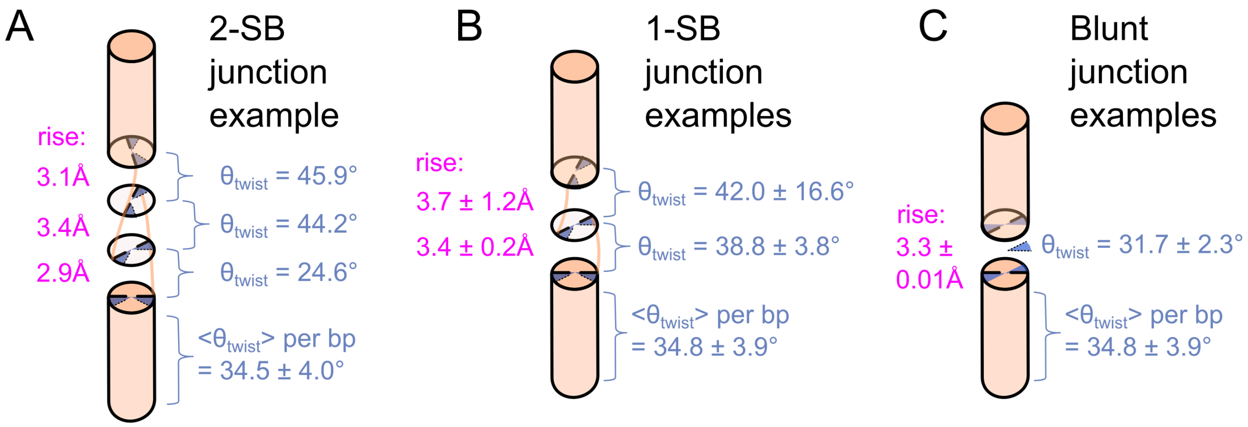


**Figure S20. Schematic of $CC1^{+10}$ coaxial DNA:DNA interface geometries**

We used the program x3DNA (110) to extract DNA geometry parameters including rise (magenta) and twist (blue). **(A)** $CC1^{+10}$ two sticky-base interface analysis of one model (PDB 9YZB). **(B)** $CC1^{+10}$ one sticky-base interface x3DNA analysis of eight one-sticky-base $CC1^{+10}$ structures (PDB 9YZA, 9NCR, 9NB0, 9YZC, 9YZD, 9YZE, 9YZF, and 9YZG). The data may be sufficient to note a tendency to slightly overwind DNA at the interface since the twist exceeds the average value for baseline B-DNA. **(C)** $CC1^{+10}$ blunt end interface x3DNA analysis of two blunt-end $CC1^{+10}$ structures (PDB 9YZI and 9YZJ). Notably, successful growth of blunt-end CC1 lattices may allow more tunable porous scaffolds via future growth of a wider variety of expansion sizes (i.e. beyond +10 and +21 bp).

The blunt variants were of particular interest because they allowed us to probe whether removing sticky overhangs would alter the helical twist across the interface, revealing otherwise hidden geometric strain introduced by overhang pairing. However, the average twist across the two blunt-ended examples was 31.7°, which was only 2.1° less than the twist values observed for sticky-end interfaces (33.8° ± 3.8°). Per Grupa *et al*. (111), unconstrained blunt end dsDNA blocks tend to have a preferred twist of 20°, suggesting that our observed blunt end twist is instead enforced by the lattice. There is a precedent for lattice-enforced DNA conformations (112). Figure prepared in Inkscape.

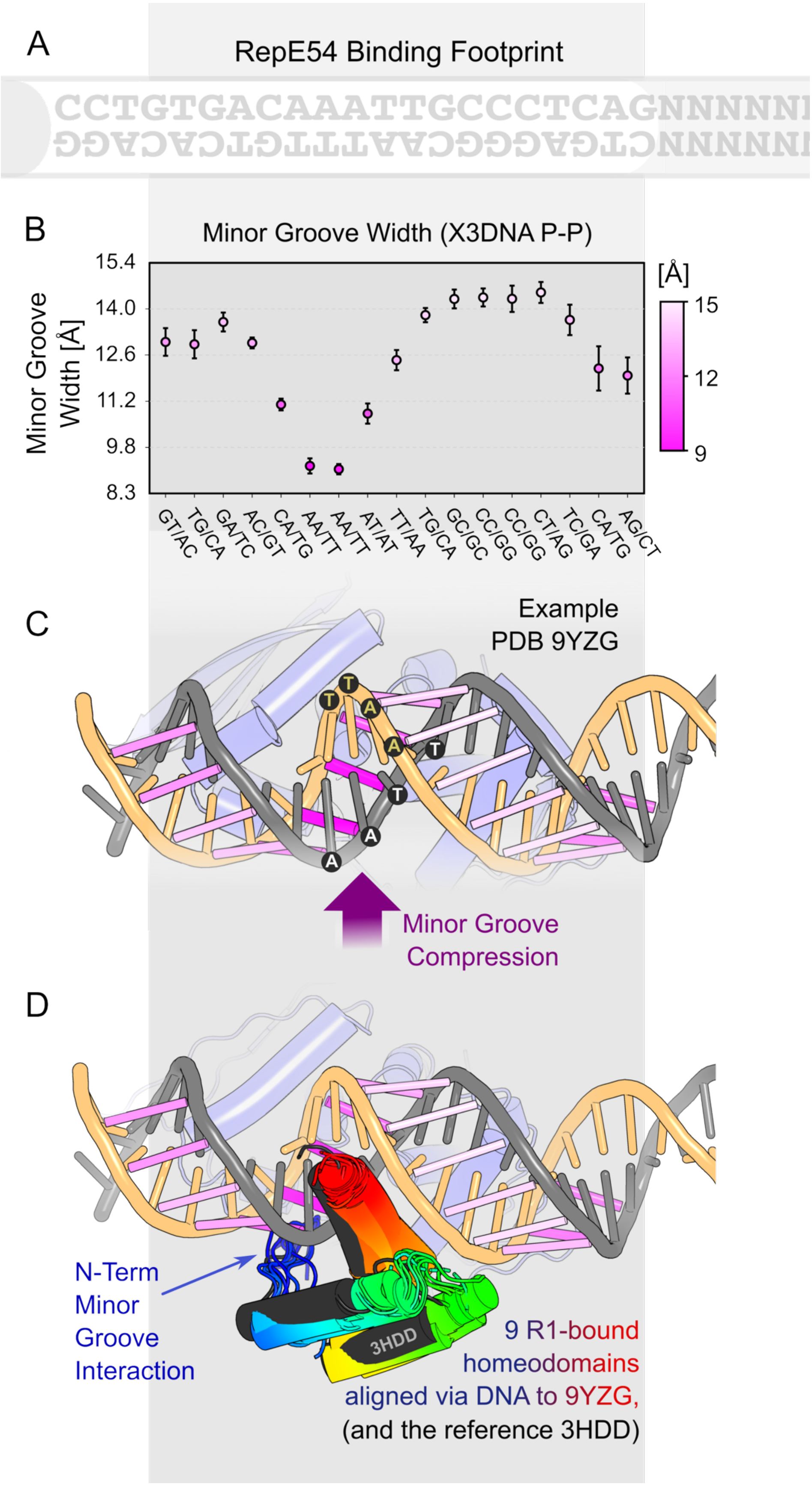

A
RepE54 Binding Footprint
CCTGTGACAAATTGCCCTCAGNNNNNN
B
Minor Groove Width (X3DNA P-P)
Minor Groove Width [Å]
15.4
14.0
12.6
11.2
9.8
8.3
[Å]
15
12
9
GT/AC
TG/CA
GA/TC
AC/GT
CA/TG
AA/TT
AA/TT
AT/AT
TT/AA
TG/CA
GC/GC
CC/GG
CC/GG
CT/AG
TC/GA
CA/TG
AG/CT
C
Example
PDB 9YZG
Minor Groove
Compression
D
N-Term
Minor
Groove
Interaction
3HDD
9 R1-bound
homeodomains
aligned via DNA to 9YZG,
(and the reference 3HDD)

**Figure S21. Minor groove compression of CC1$^{+10}$ associated with RepE54 binding**

**(A)** We typically conserved a 21 bp RepE54 binding window. **(B)** X3DNA (110) computed minor groove widths (mean and standard deviation) were tabulated for 17 CC1$^{+10}$ models that lack guest molecules: 9NB0, 9NCR, 9YZA, 9YZB, 9YZC, 9YZD, 9YDE, 9YDF, 9YDG, 9YDI, 9YZJ, 9Z08, 9Z1A, 9Z1B, 9Z1C, 9Z1D, 9Z55. **(C)** The X3DNA minor groove width is an average of two P-P distances flanking each base pair step. Using 9YZG as a representative example model, we depict each averaged vector using a variable width/color cylinder (increased magenta saturation and radius for short distances). The central 5'-AATT-3' sequence is labeled. As expected (113), our observed narrow minor groove coincides with our AT-rich region. **(D)** Nine additional CC1$^{+10}$ models have a homeodomain bound to the R1 register (9YZL, 9YZM, 9YZN, 9YZO, 9YZP, 9YZQ, 9YZR, 9YZS, 9YZT). To depict the position of those homeodomains relative to the compressed minor groove, we align each homeodomain-bound model (PyMOL align of the DNA chains) to the same 9YZG reference model and show only the R1: bound homeodomain. The C-terminal helices (red) bind the major groove opposite RepE54, and the varying homeodomain N-termini (blue) can insert an arginine into the compressed minor groove (113), a groove that is only partially obstructed by RepE54 (light blue). We also aligned the 5'-TAATTA-3' duplex in EnH-HD model 3HDD to the R1 site 5'-AATT-3' duplex and show the EnH-HD from 3HDD in black. Figure prepared in Inkscape, matplotlib, and PyMOL (82).

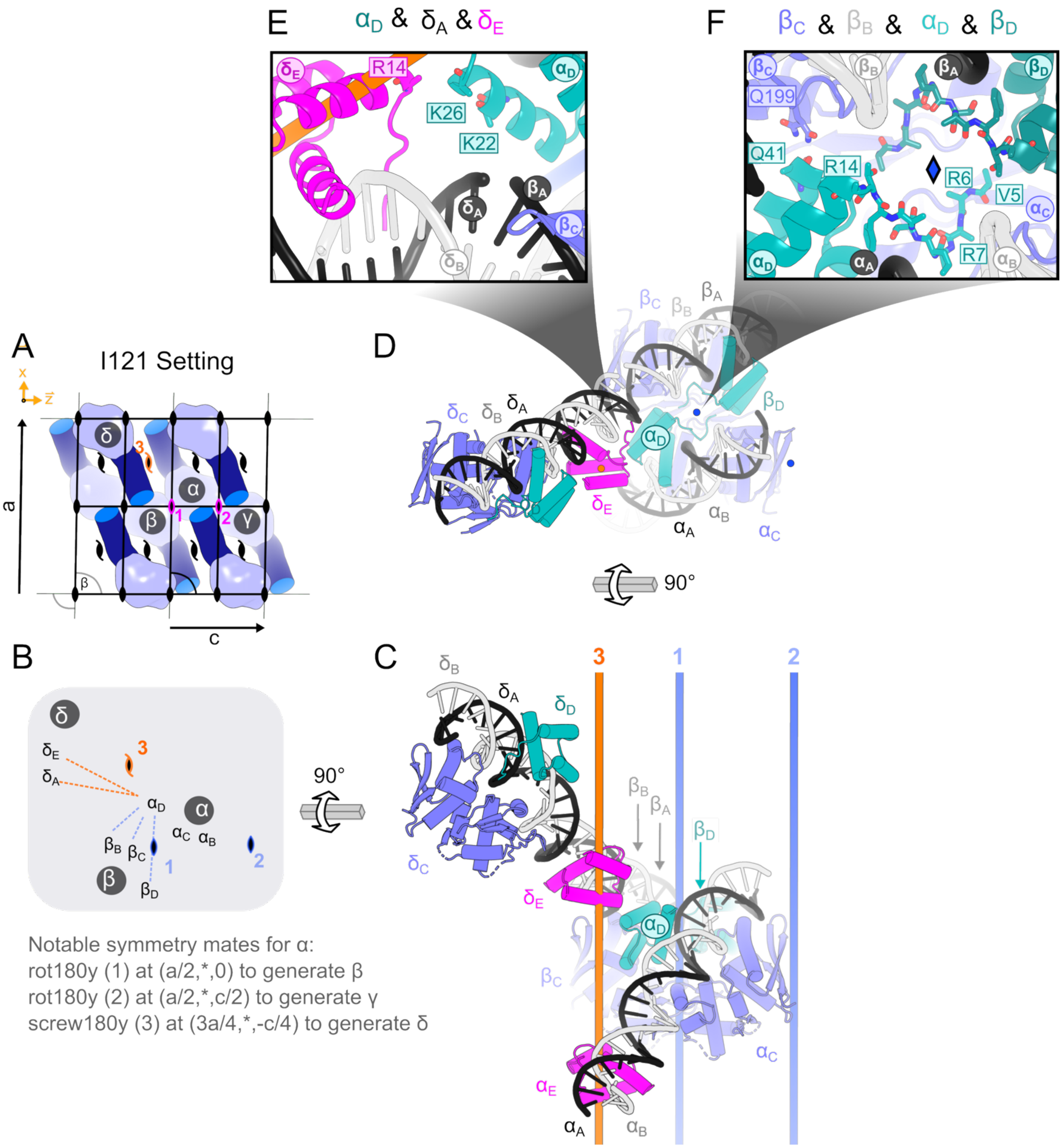


**Figure S22. Notable symmetry neighbors for the CC1 lattice (PDB entry 9YZM)**

Tabulating interactions with symmetry mates can be a dry endeavor. Here we complete one such analysis for a representative homeodomain bound structure. **(A)** For one asymmetric unit / building block, **α**, there are three neighbors that interact with the guest homeodomains. Copies **β** and **γ** are related to **α** via 2-fold symmetry axes (#1 and #2, magenta). In contrast, copy **δ** is related to **α** via a 2-fold screw axis (#3, orange). In this orientation, the major pores run horizontally (along the unit cell vector c). **(B)** A simplified diagram shows dashed lines between the constituent chains (A,B,C,D,E) within copy **α** and its neighbors, switching the color of the 2-fold axes to blue. **(C)** After a 90° rotation, each symmetry copy is depicted with two DNA chains (light gray, chain A; black, chain B), one RepE54 protein (blue, chain C), a homeodomain at the unexpected binding site opposite RepE54 (teal, chain D), and a homeodomain at the intended site (magenta, chain E). In this orientation, the 2-fold axes are depicted as blue cylinders and the 2-fold screw axis is depicted as an orange cylinder. **(D)** After another 90° rotation, the axes are once more perpendicular to the page. **(E)** This close view shows interactions of the homeodomain $\boldsymbol{\alpha_D}$. Specifically, the $\boldsymbol{\alpha_D}$ and $\boldsymbol{\delta_E}$ guest-guest interaction has a minimum distance of 3.8Å. Arg14 in $\boldsymbol{\delta_E}$ is projecting towards $\boldsymbol{\alpha_D}$. Arg14 is not resolved, but could potentially find a favorable interaction with the exposed carbonyls (helix dipole) of residues 25 and 26. The $\boldsymbol{\delta_E}$ helix dipole (residues 14-26) is also oriented favorably with respect to the $\boldsymbol{\alpha_D}$ helix dipole (residues 14-26). With a β-carbon to phosphate oxygen distances of 7.6-7.7Å, the unresolved Lys22 in $\boldsymbol{\alpha_D}$ is likely interacting favorably with the DNA in $\boldsymbol{\delta_A}$ and $\boldsymbol{\beta_A}$. **(F)** This close view shows other interactions of the homeodomain $\boldsymbol{\alpha_D}$ which is very close (3.0Å) to its symmetry neighbor across the 2-fold axis (blue diamond). The nature of the interaction is not clear, and residues prior to Val5 are not resolved. The N-terminal tail of $\boldsymbol{\alpha_D}$ is also likely interacting with its own scaffold RepE54 $\boldsymbol{\alpha_C}$ (distance 4.8Å), via steric occlusion and interactions with disordered Arg7 and Tyr8. The $\boldsymbol{\alpha_D}$ helix dipole (residues 14-26) is also close to (5.3Å) and oriented favorably (positive N-terminal end) with respect to the negative DNA backbone ($\boldsymbol{\beta_B}$). The unresolved Arg14 could also interact favorably with the $\boldsymbol{\beta_B}$ DNA. Finally, there is a direct interaction (2.8Å) between the C-terminal end of the $\boldsymbol{\alpha_D}$ alpha helix 32-41 with multi-conformation Gln199 on RepE54 ($\boldsymbol{\beta_C}$).

In summary, there are a variety of intermolecular contacts that may contribute to the stabilization of both the target binding site (magenta) and bonus site (teal). Interfacial helix capping for Gln199 seems like a good candidate for an important stabilizing contact, but this is undercut by the multiple Gln199 conformations. Speculation as to the relative importance of the other noted interfacial interactions is challenging in part because of disorder of both the sidechains and backbone. We intend to revisit analysis of these contacts after further scaffold optimization and resolution improvements. The observed interactions represent opportunities for structure-guided engineering. For example, R1 site homeodomain binding is well positioned to rigidify the scaffold crystal, and redesigning the disordered interfacial N-terminus could enhance this effect. Figure prepared in PyMOL (82) and Inkscape.

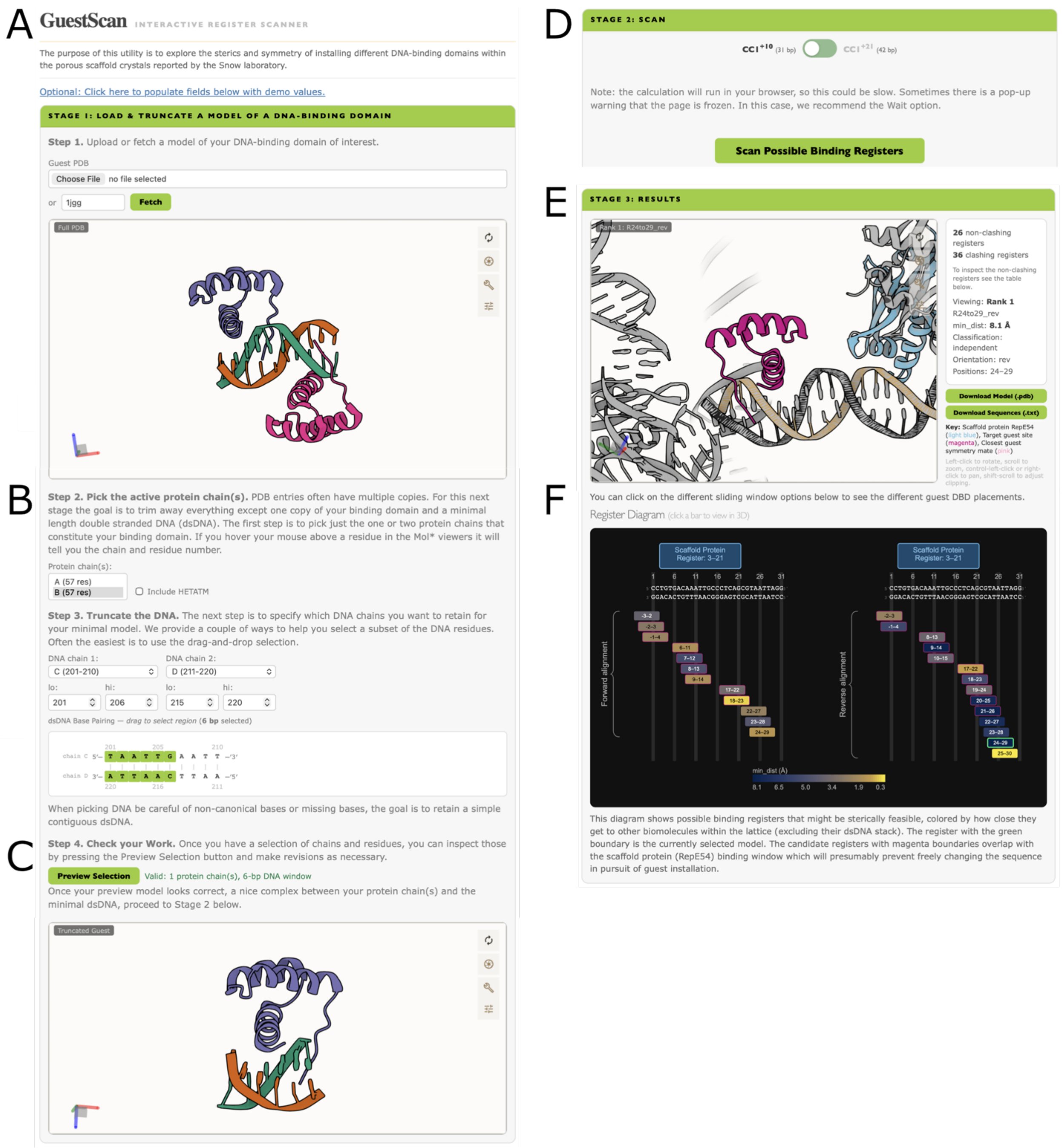


**Figure S23. Selecting Installation Sites**

We have developed open-source Python code to identify and classify candidate installation sites for DNA-binding domains (DBD) within the CC1$^{+10}$ and CC1$^{+21}$ scaffold crystals (https://github.com/cdsnow/DBDscan). An interactive web interface is also available (https://www.engr.colostate.edu/~cdasnow/guestscan.html). **(A)** A PDB model containing the DBD of interest is uploaded or retrieved using a PDB code. Many structures, including the demonstration model PDB 1JGG, contain multiple chains. **(B)** Therefore, tools are provided to (i) select the relevant protein chains and (ii) trim the bound dsDNA to the minimal footprint required for binding. **(C)** The preview window uses the fully interactive Mol* viewer (114). **(D)** The selected DBD-DNA complex is scanned through the $CC1^{+10}$ (PDB 9YZJ) or $CC1^{+21}$ (PDB 9YZK) lattice using a sliding-window procedure. Each scan position is generated by optimal superposition of the C1' atoms of the guest DNA onto each possible size-matched dsDNA window in the scaffold crystal. The calculations run in-browser and typically take several seconds. **(E)** After the scan completes, the placement that maximizes the distance to the nearest neighbor (excluding DNA chains in the dsDNA stack that harbors the installation site) is displayed. The guest DBD is shown in magenta, the scaffold protein in the same asymmetric unit in blue, and symmetry mates in gray. For installation sites near crystallographic 2-fold axes, the closest neighbor may be a symmetry-related guest molecule; the closest of these is shown in pink. Buttons allow downloading the current model, or a text file listing the DNA sequences produced by inserting the guest DNA footprint into the scaffold DNA sequence. **(F)** A summary diagram reports the results for all scan positions. Each window is colored by min_dist, defined as the minimum distance between the guest DBD and any lattice molecule other than the dsDNA stack containing the binding site. Steric feasibility is evaluated using only backbone "core atoms" (N, $C_\alpha$, C, O, $C_\beta$; and for proline also $C_\gamma$ and $C_\delta$), since side-chain clashes may be relieved by rotamer changes. Positions with core-atom overlaps < 2Å are excluded as steric clashes. Sites where the guest DNA footprint overlaps the RepE54 scaffold protein are outlined in magenta, indicating limited freedom to modify the DNA sequence without disrupting RepE54 binding.

Notably, each register generates two candidate DBD placements (forward and reverse) depending on which DNA strand in the guest model is superimposed onto the top strand in the scaffold model. Selecting a window updates the Mol* view above, allowing rapid inspection of candidate installation sites. Depending on the application, installation sites may be chosen either to minimize contacts beyond the cognate DNA interaction or to introduce potentially stabilizing contacts with symmetry mates.

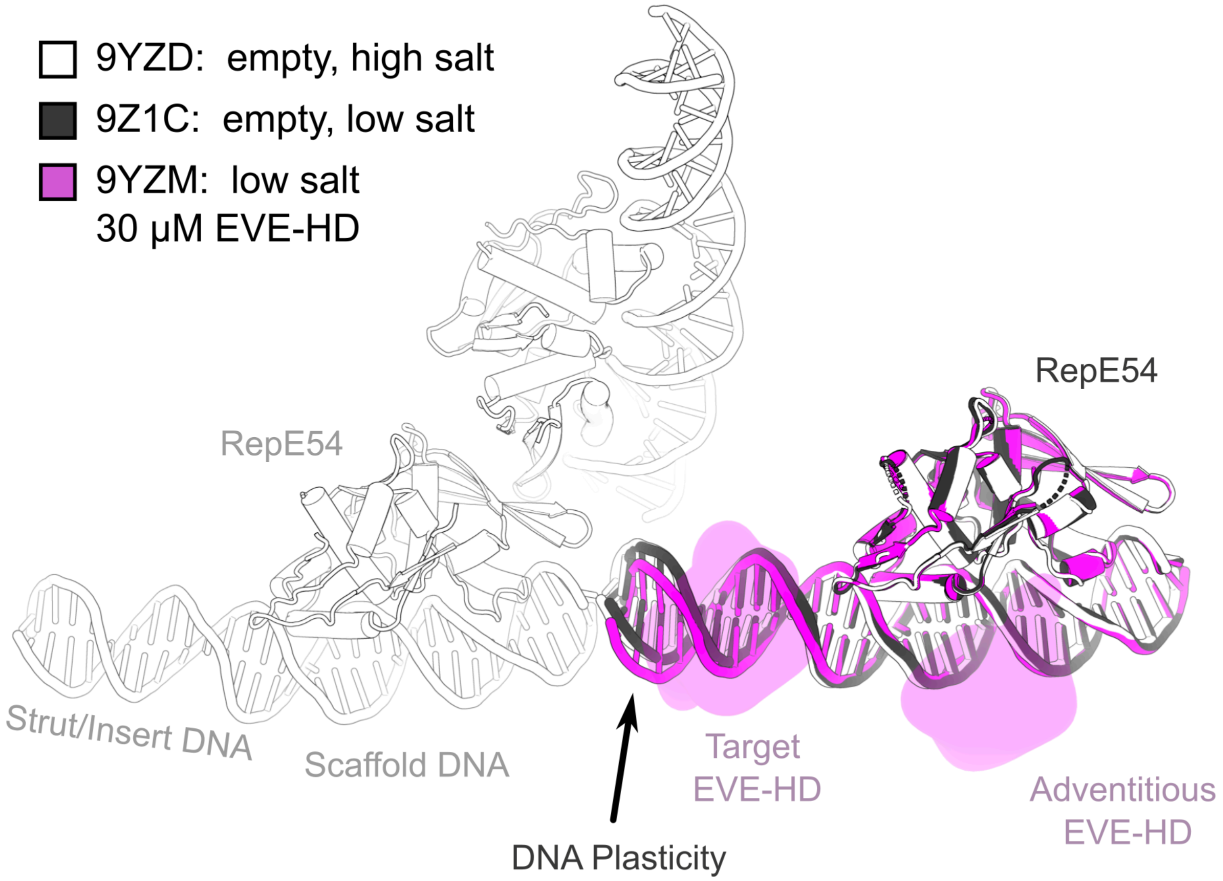


**Figure S24. DNA cantilever comparison of CC1$^{+10}$ variants with identical DNA inserts**

CC1$^{+10}$ building blocks with insert 5’ CTAATTAGGC 3’ and XRD collected under three different solvent conditions:

**White)** 30 mM Magnesium acetate, 1.3 M Lithium sulfate, 50 mM MES, pH 6.5 (PDB 9YZD).

**Black)** 50 mM KCl, 4 mM $CaCl_2$, 10% glycerol, 10 mM Tris HCl, pH 7.4 (PDB 9Z1C).

**Magenta)** 50 mM KCl, 4 mM $CaCl_2$, 10% glycerol, 10 mM Tris HCl pH 7.4 and loaded with 30 µM EVE-HD (PDB 9YZM).

Using PyMOL align, the models from 9Z1C (black) and 9YZM (magenta) were aligned via their RepE54 domain onto 9YZD (white). All biomolecules are shown as cartoons other than the guest EVE-HD domains from 9YZM which were depicted (for visual clarity) as partially transparent smoothed gaussian surfaces (gaussian_resolution = 4). Figure prepared in PyMOL (82) and Inkscape.

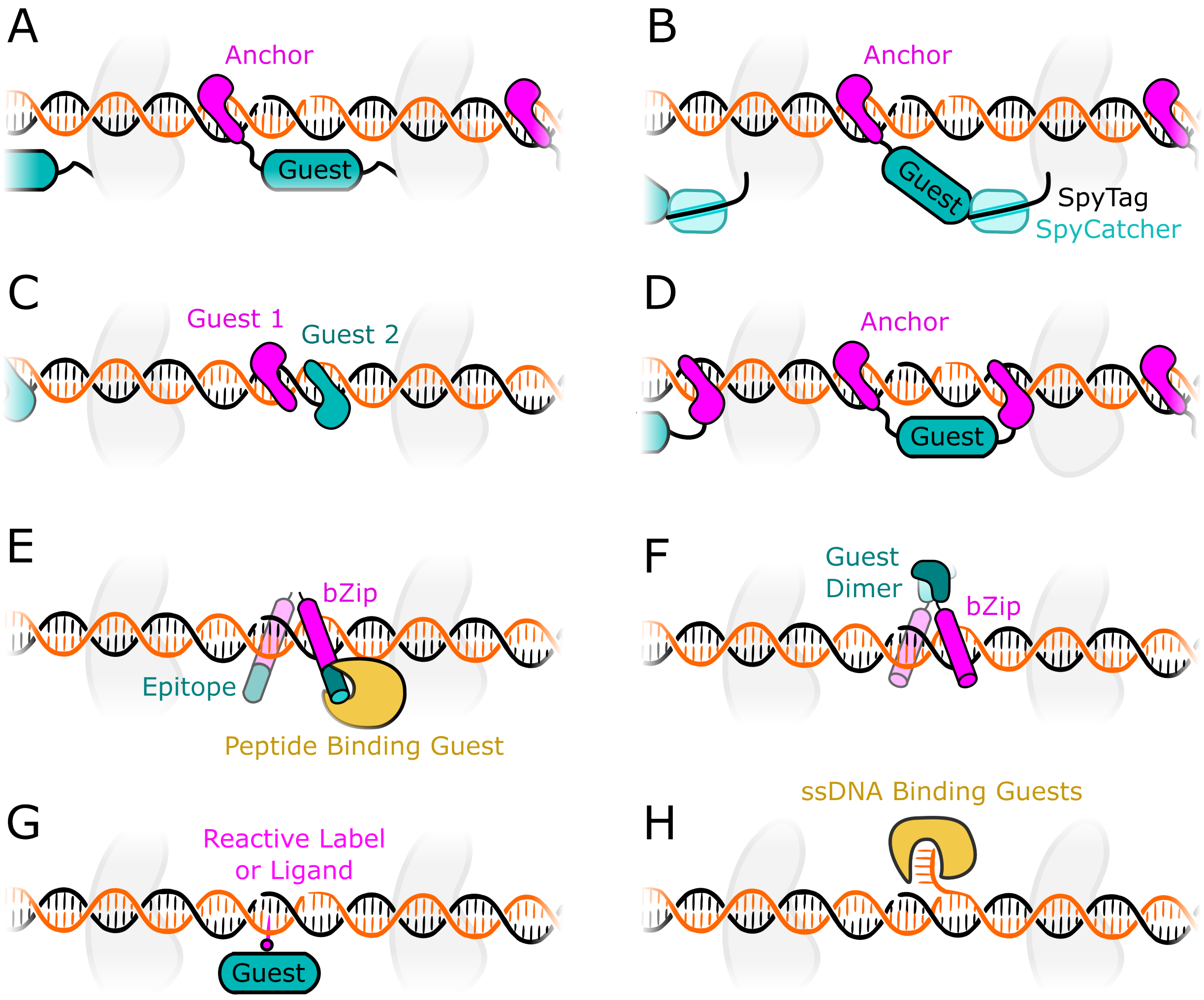


**Figure S25. Bionanotechnology Extensions to the Scaffold Crystal Platform Technology.** While the current report demonstrates how either programmed or adventitious binding can be used to install DNA-binding domains, this scaffold family should permit additional control over guest orientation via a variety of approaches, motivating the analogies to a 3D molecular pegboard or additive manufacturing on the molecular scale. Ultimately, these may be sufficient to induce order in guest domains that lack intrinsic DNA binding affinity or specificity. Notably, while adventitious binding sites do not prevent programmed guest installation for structural biology, adventitious sites could interfere with the pursuit of ultimate control. Therefore, future work will focus on eliminating off-target installation. One key concept is to move beyond single attachment sites to dual anchoring strategies. **(A)** Genetic fusion of the guest protein to the scaffold protein RepE54 and a DBD anchor. While sizable changes to the scaffold protein risk degrading expression, stability, and crystallization, we have successfully grown $CC1^{+10}$ crystals in preliminary work where an entire homeodomain was genetically fused to RepE54. Rapidly growing crystals where the anchor binding site is moved could be a systematic way to reduce variability in the guest binding position. **(B)** To limit the perturbation to the scaffold protein expression, stability, and crystallization propensity, we can engineer a RepE54 variant that offers a covalent installation motif like SpyTag (115). We have grown $CC1^{+10}$ crystals containing RepE54-ST in preliminary results. In principle, guest proteins of interest could be expressed as a genetic fusion to SpyCatcher and covalently captured by the scaffold protein SpyTag in solution or *in crystallo*. **(C)** For small guests or larger struts (e.g. $CC1^{+21}$), it should be possible to study the interactions between the guests, interactions that could provide additional stabilizing contacts to reduce disorder. The feasibility of this strategy is boosted by the proof of concept observed multiple times in this report of the same DBD bound to two registers. **(D)** Another route to capitalize on the existing guest DNA binding domains is to fuse guests of interest to N- and C-terminal DBD. We envision a modular system that can vary both the protein linkers as well as the DBD binding locations on the scaffold crystal, providing multiple methods to “tighten the straps”. **(E)** Existing DBD (e.g. homeodomains or bZip as shown) can be augmented to display bait motifs for additional guest molecules, thereby converting into structure determination “adaptors”. For example, adaptor domains could display helical peptide motifs (e.g. Sun Tag, ALFA tag) suitable to recruit additional guest proteins (e.g. peptide-binding scFv constructs). **(F)** Successful installation of the bZip parallel homodimer provides an interesting avenue to potentially pursue scaffold-assisted structure determination of homodimer guest domains. A simple expedient would be to add the bZip domains as a N- or C-terminal fusion, optionally with computational protein design of the linker regions to maximize construct rigidity. **(G)** The ability to rapidly synthesize a variety of porous scaffold crystals with varying DNA inserts should be yet another fruitful avenue. A large variety of reactive chemical moieties (e.g. click chemistry components) or bait ligands can be displayed to enhance guest capture. In prior work, we grew $ipCC1^{+10}$ crystals with DNA mismatches in the form of a 1 nt bulge (35). In preliminary work, we have grown $CC1^{+10}$ crystals with a variety of modifications including phosphorothioates, a G:G mismatch, pendant amines, a DNA:RNA hybrid strut, pendant carboxylic acids, and pendant fluorophores. The ability to grow CC1+10 crystals with significant variations in the DNA bode well for future attempts to use such variations to capture cognate proteins (e.g. for mismatch recognition). **(H)** In preliminary work, we also have grown $CC1^{+10}$ crystals with pendant ssDNA segments of 4, 6, or 15 nt, with multiple display sites and diffraction to 3.2Å. In addition to highlighting the remarkable modularity of the system, these crystals provide a motif suitable to capture proteins that bind ssDNA rather than dsDNA. The capture of nucleic acid guests is another exciting research direction that is underway. Executing multiple rounds of strand installation as pioneered by Hao *et al*. is an intriguing form of nanotechnology (17), and we ultimately aim to observe such transitions via ordered changes in the electron density map. Figure prepared in Inkscape.

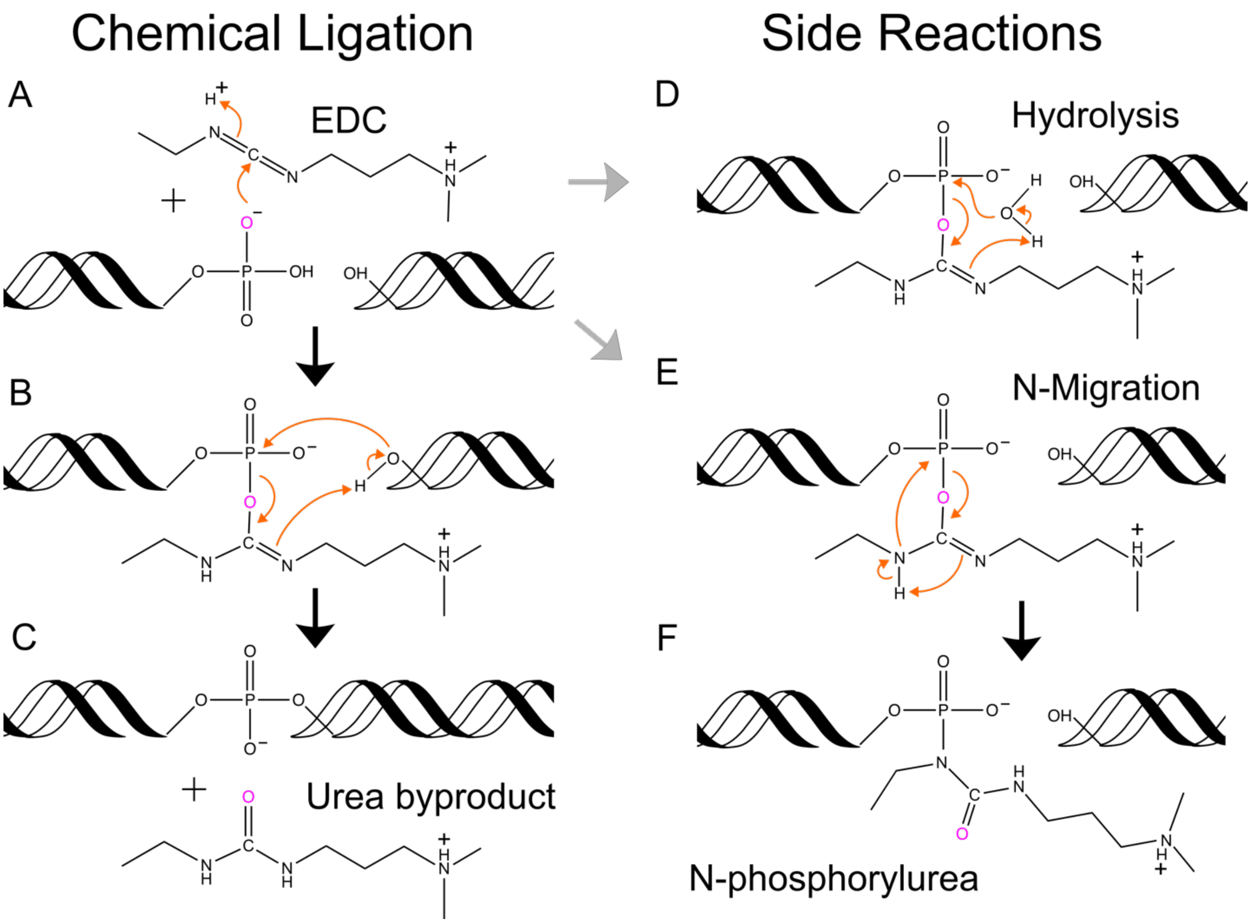


**Figure S26. Putative EDC chemical ligation mechanism and putative competing side reactions.** As demonstrated in prior publications, we use EDC for the chemical ligation of coaxial DNA interfaces within the crystals. Here we elaborate on the proposed chemical ligation mechanism and depict potential side reactions. **(A–C)** Productive Ligation. **(A)** A terminal phosphate (e.g., 3′-P) is activated by EDC to form a reactive O-phosphorylisourea intermediate. **(B)** The terminal hydroxyl (e.g., 5′-OH) of the neighboring strand attacks the phosphorus. **(C)** Formation of the native phosphodiester bond and release of the stable "urea" byproduct. Our prior work showed strong crystal stabilization despite incomplete ligation yields (34,36). A study by Hud *et al*. provides additional considerations (e.g. temperature, 3' phosphate hydrolysis, flanking base pair identities) for future efforts to optimize ligation yield (116). For example, pyrimidine-pyrimidine steps at nick sites (C|C or T|T) lead to higher ligation rates than purine-purine steps (G|G or A|A).

Here we provide schematics for reactions that could lead to incomplete ligation. These include:

**(D)** Hydrolysis. Water competes as a nucleophile, attacking the activated phosphorus. This regenerates the starting phosphate and consumes EDC but does not cap the DNA. **(E–F)** N-Migration (Dead End). **(E)** The activated intermediate might undergo an intramolecular rearrangement where an isourea nitrogen attacks the phosphorus (117). **(F)** This would result in a stable, inert N-phosphorylurea adduct, which would cap the DNA and prevent productive ligation. Figure prepared with ChemDraw Professional (Version 25.5.0.6237, Revvity Signals Software, Inc.).

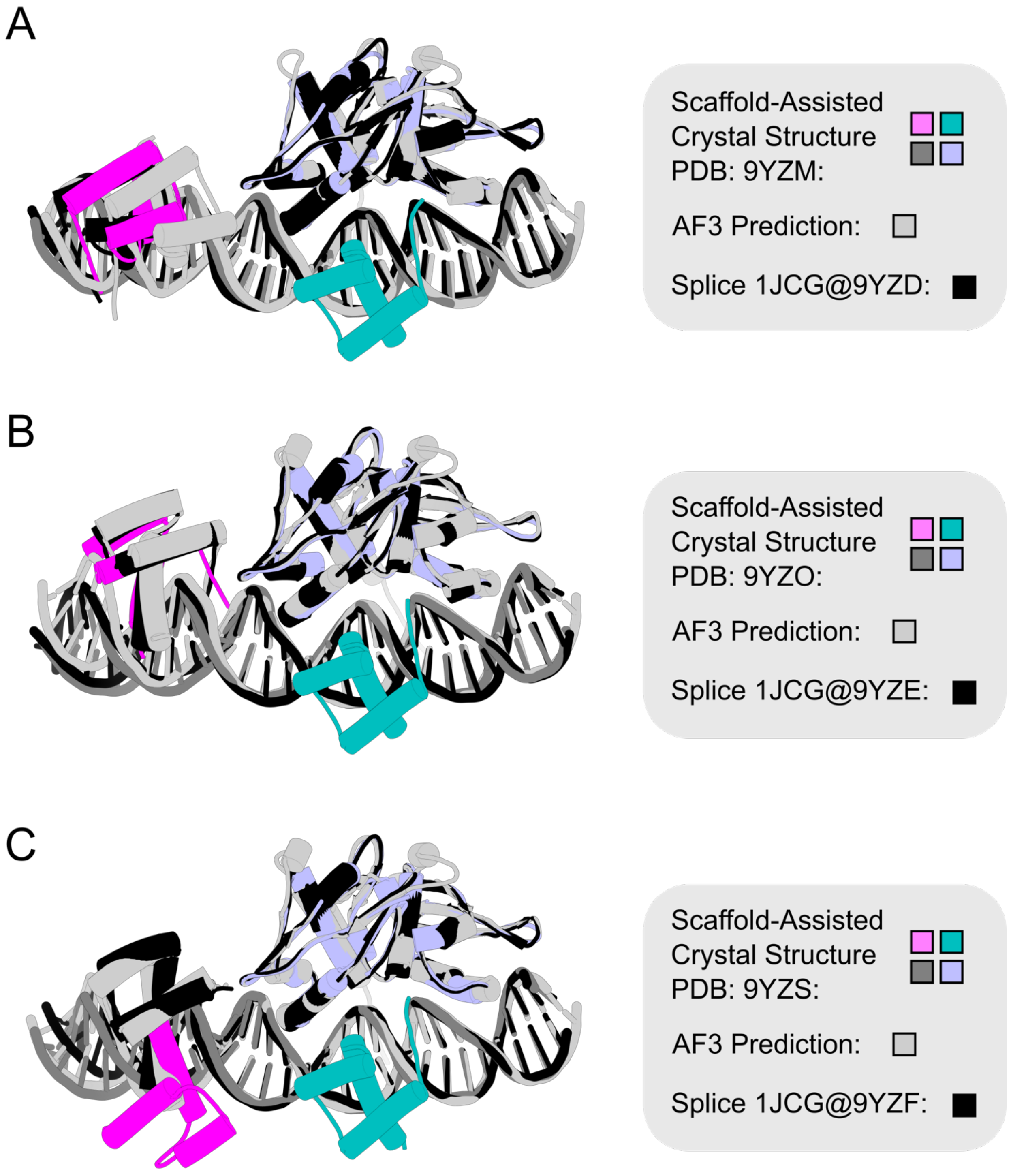
A
Scaffold-Assisted
Crystal Structure
PDB: 9YZM:
AF3 Prediction:
Splice 1JCG@9YZD:
B
Scaffold-Assisted
Crystal Structure
PDB: 9YZO:
AF3 Prediction:
Splice 1JCG@9YZE:
C
Scaffold-Assisted
Crystal Structure
PDB: 9YZS:
AF3 Prediction:
Splice 1JCG@9YZF:

**Figure S27. Structure Prediction Alignments of EVE-HD Binding**

Each panel consists of a superposition between a $CC1^{+10}$ crystal structure (multicolor), the AF3 prediction thereof (light gray), and the "expected binding" model (black) generated by splicing/grafting the 1JGG PDB model for EVE-HD onto the corresponding empty $CC1^{+10}$ structure (aligning the recognition DNA in 1JGG to the corresponding DNA in the empty $CC1^{+10}$ model). **(A)** $CC1^{+10}$ grown with asymmetric insert 5'CTAATTAGGC 3' and later loaded with EVE-HD (magenta/cyan, PDB 9YZM). Empty scaffold model 9YZD was used for the 1JGG grafting model (black). **(B)** $CC1^{+10}$ grown with asymmetric insert 5'TGATGAGCAG 3' and later loaded with EVE-HD (magenta/cyan, PDB 9YZO). Empty scaffold model 9YZE was used for the 1JGG grafting model (black). **(C)** $CC1^{+10}$ grown with asymmetric insert 5'TAATTAGGCCG 3' and later loaded with EVE-HD (magenta/cyan, PDB 9YZS). Empty scaffold model 9YZF was used for the 1JGG grafting model (black).

Summary of results: AF3 never predicted binding at the R1 site if there was one homeodomain guest and an accessible canonical site. However, if we supplied two homeodomains, AF3 reliably placed homedomains at the canonical and the R1 site. Likewise, if there was no canonical site, AF3 reliably placed homedomains at the R1 site. In the case of 9YZM, the simple splicing model was quite close. The AF3 prediction was quite close as well -- *if* we align the AF3 prediction to 9YZM via the 5'TAATTA 3' DNA. Instead, the visual difference in panel A is due to the AF3 model bending the DNA with respect to 9YZM (the 3'OH moves 1.4 nm towards the reader), and EVE-HD is carried along for the ride.

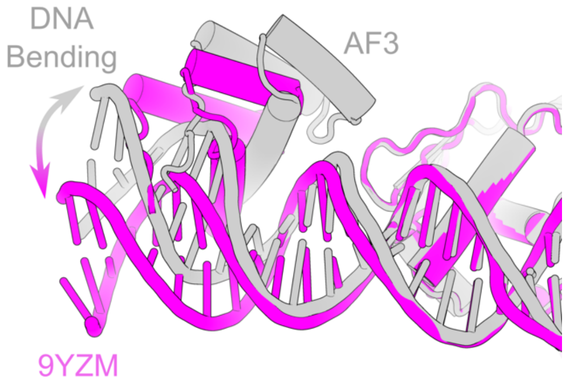

In the case of 9YZO, both the AF3 and the spliced model were good predictions of the binding experiment. Finally, in 9YZS, both the AF3 and the spliced model did not predict the experimental outcome. While all three models feature EVE-HD bound to the target site major groove, the true experimental EVE-HD binds this palindromic target in the opposite orientation supported by the palindromic binding sequence.  Figure prepared in PyMOL (82) and Inkscape.

**Table S1. CC1$^{+10}$ oligonucleotide sequences**

| **PDB Code** / *Purpose* | **CC1$^{+10}$ DNA Sequences** |
|---|---|
| **9Z08**<br>*HD site*<br>*and testing the separate insert strategy.* | 5′ – ACCTGTGACAAATTGCCCTCA – 3′<br>3′ – GACACTGTTTAACGGGAGTCT – 5′<br>5′ – GACGGTAATT – 3′<br>3′ – GCCATTAATG – 5′ |
| **9NCR**<br>*C-clamp site* | 5′ – GCCGGACCTGTGACAAATTGCCCTCAGCCCGp – 3′<br>3′ – pGGCCTGGACACTGTTTAACGGGAGTCGGGCC – 5′ |
| **9Z44**<br>*C-clamp site* | 5′ – pGGCCGGACCTGTGACAAATTGCCCTCAGCCC – 3′<br>3′ – GGCCTGGACACTGTTTAACGGGAGTCGGGCCp – 5′ |
| **9YZA**<br>*C-clamp site* | 5′ – pGCCGGACCTGTGACAAATTGCCCTCAGCCCG – 3′<br>3′ – GGCCTGGACACTGTTTAACGGGAGTCGGGCCp – 5′ |
| **9YZB**<br>*HD site* | 5′ – pTAATTACCTGTGACAAATTGCCCTCAGACGG – 3′<br>3′ – TAATGGACACTGTTTAACGGGAGTCTGCCATp – 5′ |
| **9Z1A, 9Z1B**<br>*HhaI Restriction Option* | 5′ – pCCTGTGACAAATTGCCCTCAGCCGCGCAGGC – 3′<br>3′ – GACACTGTTTAACGGGAGTCGGCGCGTCCGGp – 5′ |
| **9YZC**<br>*Test GC-rich insert* | 5′ – pCCTGTGACAAATTGCCCTCAGCGCGCGCGCG – 3′<br>3′ – GACACTGTTTAACGGGAGTCGCGCGCGCGCGp – 5′ |
| **9NB0, 9Z1D**<br>*HD site* | 5′ – pCCTGTGACAAATTGCCCTCAGGCTTGATGAG – 3′<br>3′ – GACACTGTTTAACGGGAGTCCGAACTACTCGp – 5′ |
| **9YZD, 9YZL, 9YZM, 9Z1C**. *HD site* | 5′ – pCCTGTGACAAATTGCCCTCAGCTAATTAGGC – 3′<br>3′ – GACACTGTTTAACGGGAGTCGATTAATCCGGp – 5′ |
| **9YZE, 9YZN, 9YZO, 9YZP, 9YZQ**. *HD site* | 5′ – pCCTGTGACAAATTGCCCTGCTTGATGAGCAG – 3′<br>3′ – GACACTGTTTAACGGGACGAACTACTCGTCGp – 5′ |
| **9YZF, 9YZR, 9YZS, 9YZT**. *HD site* | 5′ – pCCTGTGACAAATTGCCCTCATAATTAGGCCG – 3′<br>3′ – GACACTGTTTAACGGGAGTATTAATCCGGCGp – 5′ |
| **9YZG, 9YZU**<br>*bZip site* | 5′ – pCCTGTGACAAATTGCCCTCAGATGAGTCATA – 3′<br>3′ – GACACTGTTTAACGGGAGTCTACTCAGTATGp – 5′ |
| **9YZI**<br>*C-clamp site* | 5′ – pCCTGTGACAAATTGCCCTCAGCCCGGCCGGA – 3′<br>3′ – GGACACTGTTTAACGGGAGTCGGGCCGGCCTp – 5′ |
| **9YZJ**<br>*HD site* | 5′ – pCCTGTGACAAATTGCCCTCAGCGTAATTAGG – 3′<br>3′ – GGACACTGTTTAACGGGAGTCGCATTAATCCp – 5′ |
| **9Z55, 9Z4E**<br>*bZip site* | 5′ – pCCTGTGACAAATTGCCCTCAGCATGAGTCAT – 3′<br>3′ – GACACTGTTTAACGGGAGTCGTACTCAGTAGp – 5′ |

**Key:** Gray for the standard RepE54 footprint. Cyan for modifications inside the RepE54 footprint. Underline for target binding sites. Orange for enzyme recognition sites. “p” for phosphate.

**Table S2. Crystallization conditions of CC1$^{+10}$**

| PDB Code: | Crystallization Condition: |
|---|---|
| Abbreviations: | $Mg(OAc)_2$ = magnesium acetate, $Mg(CH_3COO)_2$; $Li_2SO_4$ = lithium sulfate; $(CH_3)_2AsO_2Na$ = sodium cacodylate; MES = 2-(N-morpholino)ethanesulfonic acid. |
| 9Z08 | 10 mM $Mg(OAc)_2$, 1.3 M $Li_2SO_4$, 50 mM $(CH_3)_2AsO_2Na$ pH 6.5 |
| 9YZB | 10 mM $Mg(OAc)_2$, 1.6 M $Li_2SO_4$, 50 mM MES pH 6.5 |
| 9NB0 | 15 mM $Mg(OAc)_2$, 0.6 M $Li_2SO_4$, 50 mM MES pH 6.5 |
| 9YZE | 20 mM $Mg(OAc)_2$, 0.3 M $Li_2SO_4$, 50 mM MES pH 6.5 |
| 9YZG | 20 mM $Mg(OAc)_2$, 1.4 M $Li_2SO_4$, 50 mM MES pH 6.5 |
| 9YZO | 20 mM $Mg(OAc)_2$, 1.7 M $Li_2SO_4$, 50 mM MES pH 6.5 |
| 9Z1D | 25 mM $Mg(OAc)_2$, 0.8 M $Li_2SO_4$, 50 mM MES pH 6.5 |
| 9YZS | 30 mM $Mg(OAc)_2$, 0.8 M $Li_2SO_4$, 50 mM MES pH 6.5 |
| 9YZR | 30 mM $Mg(OAc)_2$, 0.9 M $Li_2SO_4$, 50 mM MES pH 6.5 |
| 9NCR | 30 mM $Mg(OAc)_2$, 1.0 M $Li_2SO_4$, 50 mM MES pH 6.5 |
| 9YZM, 9YZT | 30 mM $Mg(OAc)_2$, 1.1 M $Li_2SO_4$, 50 mM MES pH 6.5 |
| 9YZL | 30 mM $Mg(OAc)_2$, 1.2 M $Li_2SO_4$, 50 mM MES pH 6.5 |
| 9YZD | 30 mM $Mg(OAc)_2$, 1.3 M $Li_2SO_4$, 50 mM MES pH 6.5 |
| 9YZC | 30 mM $Mg(OAc)_2$, 1.4 M $Li_2SO_4$, 50 mM MES pH 6.5 |
| 9YZJ | 30 mM $Mg(OAc)_2$, 1.7 M $Li_2SO_4$, 50 mM MES pH 6.5 |
| 9YZN, 9YZQ | 40 mM $Mg(OAc)_2$, 1.4 M $Li_2SO_4$, 50 mM MES pH 6.5 |
| 9YZP | 40 mM $Mg(OAc)_2$, 1.5 M $Li_2SO_4$, 50 mM MES pH 6.5 |
| 9Z55 | 40 mM $Mg(OAc)_2$, 1.8 M $Li_2SO_4$, 50 mM MES pH 6.5 |
| 9YZF | 50 mM $Mg(OAc)_2$, 1.1 M $Li_2SO_4$, 50 mM MES pH 6.5 |
| 9Z1C | 50 mM $Mg(OAc)_2$, 1.5 M $Li_2SO_4$, 50 mM MES pH 6.5 |
| 9YZI, 9Z1B, 9Z4E | 50 mM $Mg(OAc)_2$, 1.6 M $Li_2SO_4$, 50 mM MES pH 6.5 |
| 9YZU | 50 mM $Mg(OAc)_2$, 1.8 M $Li_2SO_4$, 50 mM MES pH 6.5 |
| 9Z44 | 60 mM $Mg(OAc)_2$, 1.8 M $Li_2SO_4$, 50 mM MES pH 6.5 |
| 9Z1A | 80 mM $Mg(OAc)_2$, 1.2 M $Li_2SO_4$, 50 mM MES pH 6.5 |
| 9YZA | 40 mM $Mg(OAc)_2$, 1.6 M $Li_2SO_4$, 50 mM MES pH 6.5 |

Entries are sorted first by divalent cation concentration, and secondarily by monovalent cation concentration. 9YZA is the only entry that has the variant RepE54: L53G, Q54G, E55G. See Fig. S2 for a visual depiction of these results.

**Table S3. CC1$^{+10}$ Oligonucleotide Sequences That Have Not Yet Crystallized**

| Purpose | DNA sequences |
|---|---|
| Optional HhaI Restriction | 5′ - pGCGCGACCTGTGACAAATTGCCCTCAGGCGC - 3′<br>3′ - GCGCTGGACACTGTTTAACGGGAGTCCGCGCp - 5′ |
| Optional HhaI Restriction | 5′ - pCCAAGCGCAACCTGTGACAAATTGCCCTCAG - 3′<br>3′ - GTTCGCGTTGGACACTGTTTAACGGGAGTCGp - 5′ |

**Table S4. CC1$^{+21}$ oligonucleotide sequences**

| **Status /PDB code /** *Strategy* | **dsDNA Sequence** |
|---|---|
| **Success. PDB:9YZK** *2-SB (CG/GC)* | 5′- pCGTATCTTCCCACCTGTGACAAATTGCCCTCAGACCTCATCC -3<br>3′- ATAGAAGGGTGGACACTGTTTAACGGGAGTCTGGAGTAGGGCp -5′ |
| **No diffraction on six crystals to date.** *1-SB (G/C)* | 5′- pGTATCTTCCCACCTGTGACAAATTGCCCTCAGACCTCATCCA -3′<br>3′- ATAGAAGGGTGGACACTGTTTAACGGGAGTCTGGAGTAGGTCp -5′ |
| **No crystals.** *Test 2-SB (AG/TC)* | 5′- pAGTATCTTCCCACCTGTGACAAATTGCCCTCAGACCTCATCC -3<br>3′- ATAGAAGGGTGGACACTGTTTAACGGGAGTCTGGAGTAGGTCp -5′ |
| **No crystals.** *Test 1-SB (G/C)* | 5′- pGGTATCTTCCCACCTGTGACAAATTGCCCTCAGACCTCATCCA -3′<br>3′- CATAGAAGGGTGGACACTGTTTAACGGGAGTCTGGAGTAGGTCp -5′ |
| **No crystals.** *Test 1-SB Asymmetric* | 5′- pCTGTGACAAATTGCCCTCAGACCTGTTATCATCCGATCCCAC -3′<br>3′- GGACACTGTTTAACGGGAGTCTGGACAATAGTAGGCTAGGGTp -5′ |
| **No crystals.** *Test blunt ends* | 5′- pTATCTTCCCACCTGTGACAAATTGCCCTCAGACCTCATCCAG -3′<br>3′- ATAGAAGGGTGGACACTGTTTAACGGGAGTCTGGAGTAGGTCp -5′ |
| **No crystals.** *Test 1-SB (A/T)* *HD site* | 5′- pTACATGCCGGACCTGTGACAAATTGCCCTCAGCCCGGGTAAT -3′<br>3′- TGTACGGCCTGGACACTGTTTAACGGGAGTCGGGCCCATTAAp -5′ |
| **No crystals.** *Test 3-SB (ACT/TGA)* | 5′- pAGTATCTTCCCACCTGTGACAAATTGCCCTCAGACCTCATCC -3′<br>3′- TAGAAGGGTGGACACTGTTTAACGGGAGTCTGGAGTAGGTCAp -5′ |
| **No crystals.** *Test 2-SB (TG/AC)* | 5′- pTGGAAGCCGCCTGTGACAAATTGCCCTCAGTGCTGTTGCGCT -3′<br>3′- CTTCGGCGGACACTGTTTAACGGGAGTCACGACAACGCGAACCp -5′ |
| **No crystals.** *Test 2-SB Asymmetric* **Non-complementary sticky overhangs* | 5′- pGCTGTGACAAATTGCCCTCAGGATGCTGTTGCGCTTGGACGC -3′<br>3′- ACACTGTTTAACGGGAGTCCTACGACAACGCGAACCTGCGGCp -5′ |

**Table S5. CC1$^{+21}$ crystallization conditions**

| PDB Code: | Crystallization Condition: |
|---|---|
| **Success. PDB:9YZK**<br>*2-SB (CG/GC)* | 10 mM magnesium acetate, 1300 mM lithium sulfate,<br>50 mM sodium cacodylate pH 6.5 |
| **Also grew crystals**, but no PDB code since the 3 crystals we tested had poor resolution.<br>*2-SB (CG/GC)* | 50 mM magnesium acetate, 1600 mM lithium sulfate,<br>50 mM sodium cacodylate pH 6.5 |
| **Also grew crystals**, but no PDB code since the 3 crystals we tested had poor resolution.<br>*2-SB (CG/GC)* | 50 mM magnesium acetate, 1400 mM lithium sulfate,<br>50 mM sodium cacodylate pH 6.5 |
| **Also grew crystals**, but crystals grown to date did not diffract (6 crystals attempted)<br>*1-SB (G/C)* | 50 mM magnesium acetate, 1600 mM lithium sulfate,<br>50 mM sodium cacodylate pH 6.5 |

Notably, in addition to CC1$^{+21}$ crystals successfully growing less frequently than CC1$^{+10}$ crystals (Table S1 and S3 versus Table S4), we also noted that the CC1$^{+21}$ crystals that did grow seemed to do so more slowly. This is currently an anecdotal observation, but it could nonetheless be useful for future efforts to improve the practical utility of the larger expansion size.

**Table S6. CC1$^{+10}$ XRD solvent tolerance**

| PDB Code | EDC Ligated? | XRD condition | XRD Resolution [Å] |
|---|---|---|---|
| 9Z1A | No | 40 mM magnesium acetate, 600 mM lithium sulfate, 25 mM MES pH 6.5, 25% glycerol | 3.2 |
| 9Z1B | Yes | 25 mM KCl, 2 mM $CaCl_2$, 30% glycerol, 5 mM Tris HCl pH 7.4 | 4.1 |
| 9YZD | No | 15 mM magnesium acetate, 650 mM lithium sulfate, 25 mM MES pH 6.5, 25% glycerol | 3.2 |
| 9Z1C | Yes | 25 mM KCl, 2 mM $CaCl_2$, 30% glycerol, 5 mM Tris HCl pH 7.4 | 3.1 |
| 9NB0 | No | 7.5 mM magnesium acetate, 300 mM lithium sulfate, 25 mM MES pH 6.5, 25% glycerol | 3.2 |
| 9Z1D | Yes | Final buffer resulting from a 1:1 mixture: 50 mM HEPES pH 6.5, 25 mM NaCl, 1.25 mM $MgCl_2$, and 400 mM Lithium sulfate, 7.5 mM Magnesium acetate, 12.5 mM MES, pH 6.5, 40 µM UBX-HD, 25% glycerol | 3.1 |

**Note:** the conditions listed in this Table correspond to the last buffer that crystals were incubated in, prior to liquid nitrogen freezing. Specifically, these conditions represent the contents of the cryo-preservation solution where crystals were incubated briefly (typically 1-10 seconds) prior to manual extraction via loop for flash freezing in liquid nitrogen. Readers interested in the original growth conditions should consult Table S2. Notably, 9Z1B and 9Z1C were treated identically to crystals loaded with guests (using the same buffer, Table S7). Thus, these crystals were (i) subjected to chemical ligation with EDC, (ii) incubated in 50 mM KCl, 4 mM $CaCl_2$, 10mM Tris, and 10% glycerol, then (iii) moved into the cryo-protection solution, a 1:1 mixture of the guest loading buffer and 50% glycerol.

**Narrative summary:** selected non-ligated crystals with three different DNA sequences (9Z1A, 9YZD, and 9NB0) diffracted reasonably when subjected to cryo-preservation conditions with high lithium sulfate (half the concentration in their mother liquor). A crystal (9Z1B) with the same DNA insert sequence as 9Z1A still diffracted when the salt was dramatically reduced. Another crystal (9Z1C) with the same DNA insert sequence as 9YZD diffracted just as well despite dramatic salt reduction. To demonstrate that reasonable XRD resolution is not limited to these specific conditions, we include entry 9Z1D, which demonstrated similar diffraction quality as 9NB0 but under more complex salt conditions that resulted from mixing two buffers.

**Table S7. $CC1^{+10}$ guest loading conditions**

| Guest Protein | PDB Code | Loading Condition |
|---|---|---|
| EVE-HD | 9YZL | 15 mM KCl, 4 mM $CaCl_2$, 10% glycerol, 10 mM Tris HCl pH 7.4, 30 μM EVE-HD |
| | 9YZM | 50 mM KCl, 4 mM $CaCl_2$, 10% glycerol, 10mM Tris HCl pH 7.4, 30 μM EVE-HD |
| | 9YZN | 25 mM KCl, 2 mM $CaCl_2$, 10% glycerol, 10 mM Tris HCl pH 7.4, 30 μM EVE-HD |
| | 9YZO | 50 mM KCl, 4 mM $CaCl_2$, 10% glycerol, 10 mM Tris HCl pH 7.4, 30 μM EVE-HD |
| | 9YZS | 50 mM KCl, 4 mM $CaCl_2$, 10% glycerol, 10 mM Tris HCl pH 7.4, 30 μM EVE-HD |
| UBX-HD | 9YZP | 50 mM KCl, 4 mM $CaCl_2$, 10% glycerol, 10 mM Tris HCl pH 7.4, 40 μM UBX-HD |
| | 9YZR | 35 mM KCl, 4 mM $CaCl_2$, 10% glycerol, 10 mM Tris HCl pH 7.4, 80 μM UBX-HD |
| ANTP-HD | 9YZT | 50 mM KCl, 4 mM $CaCl_2$, 10% glycerol, 10 mM Tris HCl pH 7.4, 23 μM ANTP-HD |
| EnH-eGFP | 9YZQ | 50 mM KCl, 4 mM $CaCl_2$, 10% glycerol, 10 mM Tris HCl pH 7.4, 22 μM EnH-eGFP |
| C-clamp | 9Z44 | 50 mM KCl, 4 mM $CaCl_2$, 10% glycerol, 10 mM Tris HCl pH 7.4, 90 μM C-clamp |
| bZip | 9YZU | 50 mM KCl, 4 mM $CaCl_2$, 10% glycerol, 10 mM Tris HCl pH 7.4, 33 μM bZip |
| | 9Z4E | 50 mM KCl, 4 mM $CaCl_2$, 10% glycerol, 10 mM Tris HCl pH 7.4, 100 μM bZip |

**Note:** the conditions here are the lengthy guest incubations (i.e. 24 hr or more). Each crystal was also briefly (1-10 s) subjected to a different condition for cryo preservation. The cryo preservation conditions were a 1:1 dilution of these loading conditions with 50% glycerol (v/v). Thus, when putative loaded crystals from this table were subjected to cryo-protection solutions, the final glycerol concentration would be 30%.

**Table S8. MAP_CHANNELS calculations for all CC1$^{+10}$ and CC1$^{+21}$ structures**

| PDB | EDC Ligated? | Guest Status | MAP_CHANNELS Solvent Content | MAP_CHANNELS Maximum Radius 1D [Å] | Reso-lution [Å] |
|---|---|---|---|---|---|
| 9NB0 | No | no | 0.82 | 25.6 | 3.2 |
| 9NCR | No | no | 0.83 | 27 | 3.2 |
| 9YZA | No | no | 0.83 | 27.2 | 3.1 |
| 9YZB | No | no | 0.83 | 27.4 | 4.1 |
| 9YZC | No | no | 0.83 | 27.8 | 3.2 |
| 9YZD | No | no | 0.83 | 27.2 | 3.1 |
| 9YZE | No | no | 0.84 | 26.4 | 4.1 |
| 9YZF | No | no | 0.83 | 27 | 3.1 |
| 9YZG | No | no | 0.82 | 26.6 | 3.1 |
| 9YZI | No | no | 0.84 | 26.4 | 4 |
| 9YZJ | No | no | 0.83 | 25.2 | 3 |
| 9YZL | Yes | EVE-HD | 0.79 | 25.4 | 3.0 |
| 9YZM | Yes | EVE-HD | 0.78 | 26.2 | 3.1 |
| 9YZN | Yes | EVE-HD | 0.8 | 26.4 | 3.2 |
| 9YZO | Yes | EVE-HD | 0.79 | 25.6 | 3.0 |
| 9YZP | Yes | UBX-HD | 0.82 | 27.4 | 3.7 |
| 9YZQ | Yes | EnH-eGFP | 0.82 | 26.4 | 3.8 |
| 9YZR | Yes | UBX-HD | 0.81 | 20.6 | 3.4 |
| 9YZS | Yes | EVE-HD | 0.78 | 22 | 3.1 |
| 9YZT | Yes | ANTP-HD | 0.79 | 20 | 3.3 |
| 9YZU | Yes | bZip | 0.8 | 25.8 | 3.1 |
| 9Z08 | No | no | 0.84 | 26.6 | 4.2 |
| 9Z1A | No | no | 0.83 | 27.8 | 3.1 |
| 9Z1B | Yes | no | 0.83 | 28 | 3.9 |
| 9Z1C | Yes | no | 0.83 | 26.4 | 3.0 |
| 9Z1D | Yes | no | 0.83 | 27 | 3.1 |
| 9Z44 | Yes | C-clamp | 0.84 | 28.4 | 7.2 |
| 9Z4E | Yes | bZip | 0.81 | 22.8 | 3.6 |
| 9Z55 | No | no | 0.83 | 25.6 | 3.9 |
| 9YZK | No | no | 0.9 | 42.6 | 5.2 |

Models are listed alphabetically, save for the larger expansion scaffold example CC1$^{+21}$ (9YZK) which is placed last. In summary, the MAP_CHANNELS data shows that the binding of very small guest proteins only reduces the solvent content by 5% or less (e.g. from ~83% to 78%), and guest installation need not noticeably impede the subsequent diffusion of other guest proteins. The largest observed pore constriction (for ANTP-HD loaded crystals) still would permit the diffusion of guest proteins up to 4 nm diameter.

**Table S9. Binding register indexing conventions:**

| Binding Register (named registers of particular interest) | Residue Indices | Typical Sequence | Position |
|---|---|---|---|
| R1 | 10-15 | AATTGC | Major Groove Opposite RepE54 |
| R2 | 18-23 | TCATAA | Major Groove Near RepE54 A76 |
| R:21-26 | 21-26 | TAATTA | Exposed Strut |
| R:22-27 | 22-27 | TGATGA | Exposed Strut |
| R:23-28 | 23-28 | TAATTA | Exposed Strut |
| R:1-5 | 1-5 | GCCGG | Exposed Strut |
| R:22-30 | 22-30 | ATGAGTCA | Exposed Strut |

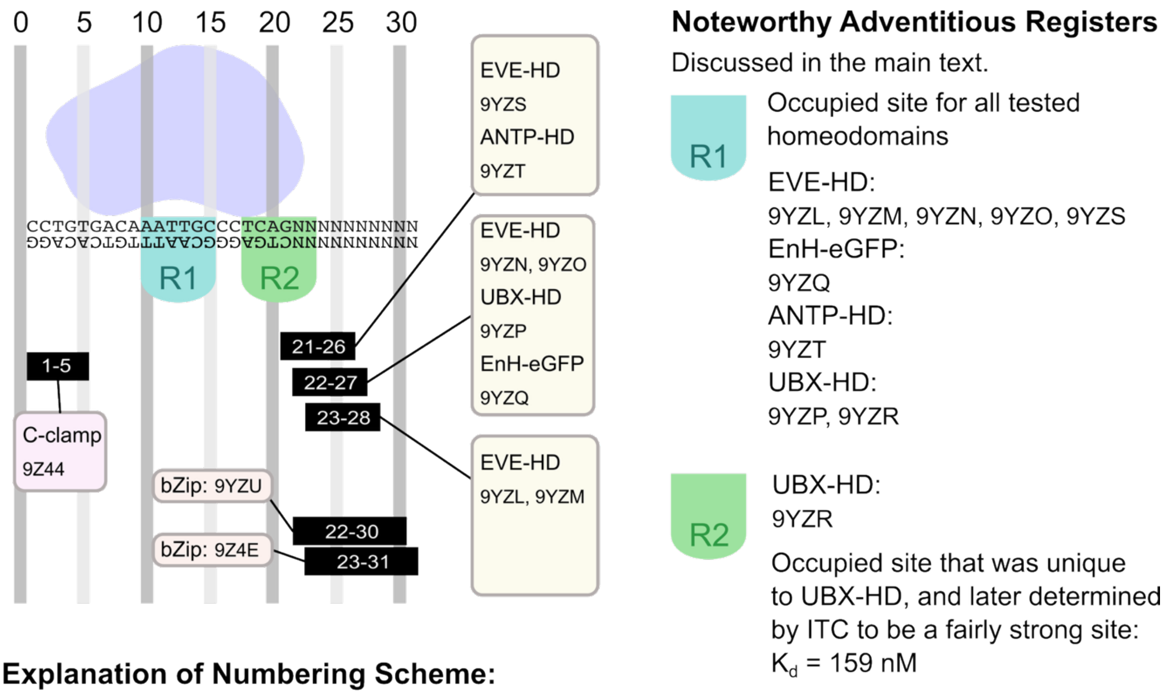


**Explanation of Numbering Scheme:**
To have a uniform base numbering scheme we use the asymmetric $CC1^{+10}$ as the starting model as shown, with the strand containing AAATT selected as the top strand.

**Table S10. RMSD for Guest Models Compared to Reference Models**

| Gues | PDB | Chain | Binding Sites | Comparison | $RMSD_{C\alpha}$ [Å] | $\#C_{\alpha}$ |
|---|---|---|---|---|---|---|
| EVE-HD | 9YZL | D | R1 (AATTGC) | 1JGG.B | 1.28 | 56 |
| | | E | R:23-28 (TAATTA) | TAATTG | 1.45 | 5 |
| | 9YZM | D | R1 (AATTGC) | 1JGG.B | 1.19 | 56 |
| | | E | R:23-28 (TAATTA) | TAATTG | 1.14 | 53 |
| | 9YZN | D | R1 (AATTGC) | 1JGG.B | 1.35 | 56 |
| | | E | R:22-27 (TGATGA) | TAATTG | 0.77 | 56 |
| | 9YZO | D | R1 (AATTGC) | 1JGG.B | 1.44 | 56 |
| | | E | R:22-27 (TGATGA) | TAATTG | 0.78 | 56 |
| | 9YZS | D | R1 (AATTGC) | 1JGG.B | 1.16 | 57 |
| | | E | R:21-26 (TAATTA) | TAATTG | 0.81 | 55 |
| UBX-HD | 9YZP | D | R1 (AATTGC) | 1B8I.A | 0.72 | 54 |
| | | n.d. | R:22-27 (TGATGA) | CCATAA | n.d. | n.d. |
| | 9YZR | E | R1 (AATTGC) | 1B8I.A | 1.83 | 37 |
| | | D | R2 (TCATAA) | CCATAA | 2.42 | 45 |
| EnH-eGFP | 9YZQ | D | R1 (AATTGC) | 3HDD.A | 0.65 | 55 |
| | | n.d. | R:22-27 (TGATGA) | TAATTA | n.d. | n.d. |
| ANTP-HD | 9YZT | D | R1 (AATTGC) | 4XID.A | 0.56 | 54 |
| | | E | R:21-26 (TAATTA) | TAATGG | 2.45 | 44 |
| C-clamp | 9Z44 | D | R:1-5 (GCCGG) | 7DTA.A GCCGG | 0.52 | 19 |
| bZip | 9YZU | D/E | R:22-30 (ATGAGTCA) | 1YSA.CD ATGAGTCA | 0.73 | 50 |
| bZip | 9Z4E | D/E | R:23-31 (ATGAGTCA) | 1YSA.CD ATGAGTCA | 2.57 | 49 |

Each modeled guest DBD was superimposed (all matching alpha carbons) onto an existing chain (or chains) from a PDB reference model. Homeodomain $RMSD_{C\alpha}$ values above 1Å typically arose from variations in the N-terminus placement, while $RMSD_{C\alpha}$ values above 2Å arose from less ordered and less complete guest DBD structures. Changes in chain E drove the higher $RMSD_{C\alpha}$ value for the bZip in 9Z4E. Collectively, these results show that these model guest proteins adopt conformations similar to existing reference structures despite solution condition differences and, in many cases, DNA sequence differences. ANTP-HD has affinity for binding both TAATTA and TAATGG (118). The existing ANTP-HD:DNA examples in the PDB (1AHD, 4XIC, 9ANT, 4XID) correspond to the latter. To our knowledge, our new entry 9YZT is the first showing the ANTP-HD bound to TAATTA. Similarly, existing UBX-HD complexes in the PDB (1B8I, 4CYC, 4UUS) show UBX-HD bound to CCATAA, which differs from our 9YZR sequence TCATAA. Finally, 1JGG is the sole pre-existing complex for EVE-HD:TAATGG. Thus, our EVE-HD complexes are likewise new.

**Table S11. Protein sequences**

| Protein Name | Amino Acid Sequence |
|---|---|
| RepE54 Transcription Factor | MRGSHHHHHHGSMAETAVINHKKRKNSPRIVQSNDLTEAAYSL SRDQKRMLYLFVDQIRKSDGTLQEHDGICEIHVAKYAEIFGLTS AEASKDIRQALKSFAGKEVVFYRPEEDAGDEKGYESFPWFIKPA HSPSRGLYSVHINPYLIPFFIGLQNRFTQFRLSETKEITNPYAMRL YESLCQYRKPDGSGIVSLKIDWIIERYQLPQSYQRMPDFRRRFLQ VCVNEINSRTPMRLSYIEKKKGRQTTHIVFSFRDITSMTTG |
| RepE54 variant (L53G, Q54G, E55G, I116C) *Mutations underlined* | MRGSHHHHHHGSMAETAVINHKKRKNSPRIVQSNDLTEAAYSL SRDQKRMLYLFVDQIRKSDGT**GGG**HDGICEIHVAKYAEIFGLTS AEASKDIRQALKSFAGKEVVFYRPEEDAGDEKGYESFPWFIKPA HSPSRGLYSVHINPYLIPFFIGLQNRFTQFRLSETKEITNPYAMRL YESLCQYRKPDGSGIVSLKIDWIIERYQLPQSYQRMPDFRRRFLQ VCVNEINSRTPMRLSYIEKKKGRQTTHIVFSFRDITSMTTG |
| Engrailed Homeodomain fused to eGFP (EnH-eGFP) *Linker sequence** *underlined* | MEKRPRTAFSSEQLARLKREFNENRYLTERRRQQLSSELGLNEA QIKIWFQNKRAKIKKSTSQFYLNEMVSKGEELFTGVVPILVELD GDVNGHKFSVSGEGEGDATYGKLTLKFICTTGKLPVPWPTLVTT LTYGVQCFSRYPDHMKQHDFFKSAMPEGYVQERTIFFKDDGNY KTRAEVKFEGDTLVNRIELKGIDFKEDGNILGHKLEYNYNSHNV YIMADKQKNGIKVNFKIRHNIEDGSVQLADHYQQNTPIGDGPVL LPDNHYLSTQSALSKDPNEKRDHMVLLEFVTAAGITLGMDELY KHHHHHH |
| Even-skipped Homeodomain (Eve-HD) | GTGSVRRYRTAFTRDQLGRLEKEFYKENYVSRPRRCELAAQLN LPESTIKVWFQNRRMKDKRQR |
| Ultrabithorax Homeodomain (UBX-HD) | GTRRRGRQTYTRYQTLELEKEFHTNHYLTRRRRIEMAHALCLTE RQIKIWFQNRRMKLKKEI |
| Antennapedia Homeodomain (AntP-HD) | GTRKRGRQTYTRYQTLELEKEFHFNRYLTRRRRIEIAHALCLTE RQIKIWFQNRRMKWKKEN |
| Our bZip: adapted from GCN4. See Fig. S5. | DPAALKRARNTEAARRSRARKLQRMKKQC |
| Our C-clamp: with and w/o TAMRA See Fig. S6 | Acetyl-KKCRKVYGMERRDLWCTACRWKKACQRF{Lys(TAMRA)} ------------------------------------------------------------------------------ KKCRKVYGMERRDLWCTACRWKKACQRF |

*Regarding the linker sequence origin: it has the same amino acid content as the TEV protease cleavage sequence (119) but is a non-cleavable control due to reversal of the sequence. Histags colored cyan.

**Table S12. X-ray diffraction statistics for CC1$^{+10}$ PDB entry 9Z08**

| **PDB code** | **9Z08** |
|---|---|
| **Data collection** | |
| Light source | Synchrotron |
| Wavelength (Å) | 1 |
| Resolution Range (Å) | 37.31-4.01 (4.44 -4.22) |
| Space group | I121 |
| **Unit cell dimensions** | |
| a, b, c (Å) | 75.045, 133.505, 135.653 |
| $\alpha$, β, ɣ (°) | 90.00, 96.08, 90.00 |
| Unique reflections | 19241 (1068) |
| Multiplicity | 1.0 (1.0) |
| Completeness (%) | 97.62 (86.56) |
| Mean I/sigma(I) | 3.51 (0.76) |
| Wilson B-factor | 146.58 |
| R-merge | 0.0891 (1.3) |
| R-meas | 0.1260 (1.839) |
| R-pim | 0.0891 (1.3) |
| CC1/2 | 0.996 (0.465) |
| **Refinement** | |
| Reflections used in refinement | 9405 (1068) |
| Reflections used for Rfree | 940 (117) |
| R-work | 0.2398 (0.3558) |
| R-free | 0.2537 (0.4238) |
| Number of non-hydrogen atoms | 2980 |
| Macromolecules | 2980 |
| Ligands | 2 |
| Solvent | 0 |
| Protein residues | 213 |
| RMS (bonds) (Å) | 0.21 |
| RMS (angles) (°) | 0.41 |
| Ramachandran favored (%) | 98 |
| Ramachandran allowed (%) | 2 |
| Ramachandran outliers (%) | 0 |
| Rotamer outliers (%) | 0 |
| Clashscore | 3 |
| Average B-factor | 181 |

**Table S13. X-ray diffraction statistics for CC1$^{+10}$ PDB entries 9YZA and 9NCR**

| PDB code | 9YZA | 9NCR |
|---|---|---|
| **Data collection** | | |
| Light source | Synchrotron | Synchrotron |
| Wavelength (Å) | 1 | 1 |
| Resolution Range (Å) | 46.67-3.05 (3.12-3.05) | 34.76-3.24 (3.32-3.24) |
| Space group | I121 | I121 |
| **Unit cell dimensions** | | |
| a, b, c (Å) | 74.35, 124.35, 137.54 | 76.03, 124.86, 139.09 |
| $\alpha$, β, ɣ (°) | 90.00, 90.30, 90.00 | 90.00, 91.85, 90.00 |
| Unique reflections | 39868 (218) | 42728 (79) |
| Multiplicity | 1.8 (3.5) | 1 (1.1) |
| Completeness (%) | 95.37 (99.1) | 96.62 (90.90) |
| Mean I/sigma(I) | 6.6 (1.07) | 5.56 (0.93) |
| Wilson B-factor | 102.99 | 104.68 |
| R-merge | 0.079 (2.277) | 0.052 (1.415) |
| R-meas | 0.094 (2.695) | 0.074 (1.415) |
| R-pim | 0.050 (1.428) | 0.052 (1.415) |
| CC1/2 | 0.999 (0.603) | 0.998 (0.417) |
| **Refinement** | | |
| Reflections used in refinement | 22801(1255) | 20000 (1217) |
| Reflections used for Rfree | 2279 (142) | 1994 (130) |
| R-work | 0.2959 (0.6554) | 0.2203 (0.4211) |
| R-free | 0.3090 (0.5821) | 0.2310 (0.4611) |
| Number of non-hydrogen atoms | 3044 | 3060 |
| Macromolecules | 3044 | 3060 |
| Ligands | 2 | 2 |
| Solvent | 0 | 0 |
| Protein residues | 217 | 216 |
| RMS (bonds) (Å) | 0.17 | 0.18 |
| RMS (angles) (°) | 0.41 | 0.41 |
| Ramachandran favored (%) | 98 | 98 |
| Ramachandran allowed (%) | 2 | 2 |
| Ramachandran outliers (%) | 0 | 0 |
| Rotamer outliers (%) | 1 | 0 |
| Clashscore | 7 | 7 |
| Average B-factor | 132 | 179 |

**Table S14. X-ray diffraction statistics for CC1$^{+10}$ PDB entries 9YZB and 9Z1A**

| PDB code | 9ZYB | 9Z1A |
|---|---|---|
| **Data collection** | | |
| Light source | Synchrotron | Synchrotron |
| Wavelength (Å) | 1 | 1 |
| Resolution Range (Å) | 38.81-3.19 (3.27-3.19) | 46.81-3.10 (3.17-3.10) |
| Space group | I121 | I121 |
| **Unit cell dimensions** | | |
| a, b, c (Å) | 75.53, 130.59, 135.53 | 74.75, 124.53, 137.76 |
| $\alpha$, β, ɣ (°) | 90.00, 90.13, 90.00 | 90.00, 90.31, 90.00 |
| Unique reflections | 38587 (2738) | 21447 (553) |
| Multiplicity | 1.0 (1.0) | 3.2 (3.2) |
| Completeness (%) | 90.74 (88.62) | 100 (80.7) |
| Mean I/sigma(I) | 3.36 (0.88) | 3.2 (0.9) |
| Wilson B-factor | 79.7 | 100.86 |
| R-merge | 0.1158 (3.925) | 0.191 (0.904) |
| R-meas | 0.1638 (5.551) | 0.224 (1.06) |
| R-pim | 0.1158 (3.925) | 0.127 (0.603) |
| CC1/2 | 0.996 (0.252) | 0.999 (0.739) |
| **Refinement** | | |
| Reflections used in refinement | 19949 (966) | 21445 (1107) |
| Reflections used for Rfree | 1987 (101) | 2140 (122) |
| R-work | 0.3045 (0.4672) | 0.3070 (0.6290) |
| R-free | 0.3281 (0.4944) | 0.3351 (0.6628) |
| Number of non-hydrogen atoms | 3110 | 2936 |
| Macromolecules | 3110 | 2936 |
| Ligands | 1 | |
| Solvent | 0 | 0 |
| Protein residues | 221 | 216 |
| RMS (bonds) (Å) | 0.16 | 0.14 |
| RMS (angles) (°) | 0.42 | 0.34 |
| Ramachandran favored (%) | 98 | 98 |
| Ramachandran allowed (%) | 2 | 2 |
| Ramachandran outliers (%) | 0 | 0 |
| Rotamer outliers (%) | 0 | 0 |
| Clashscore | 4 | 3 |
| Average B-factor | 101 | 124 |

**Table S15. X-ray diffraction statistics for CC1$^{+10}$ PDB entries 9YZC and 9NB0**

| PDB code | 9YZC | 9NB0 |
|---|---|---|
| **Data collection** | | |
| Light source | Synchrotron | Synchrotron |
| Wavelength (Å) | 1 | 1 |
| Resolution Range (Å) | 46.80-3.16 (3.23-3.16) | 39.55-3.20 (3.28-3.20) |
| Space group | I121 | I121 |
| **Unit cell dimensions** | | |
| a, b, c (Å) | 76.03, 124.86, 139.09 | 75.53, 119.24, 139.79 |
| $\alpha$, β, ɣ (°) | 90.00, 91.85, 90.00 | 90.00, 90.63, 90.00 |
| Unique reflections | 21911 (4255) | 19630 (3755) |
| Multiplicity | 3.5 (3.6) | 3.2 (2.0) |
| Completeness (%) | 99.1 (99.7) | 97.61 (96.06) |
| Mean I/sigma(I) | 4.2 (1.1) | 14.6 (3.13) |
| Wilson B-factor | 100.86 | 109.47 |
| R-merge | 0.186 (1.062) | 0.02144 (0.213) |
| R-meas | 0.221 (1.251) | 0.03032 (0.3012) |
| R-pim | 0.118 (0.655) | 0.02144 (0.213) |
| CC1/2 | 0.995 (0.737) | 1 (0.995) |
| **Refinement** | | |
| Reflections used in refinement | 21911 (1160) | 19481 (1186) |
| Reflections used for Rfree | 2190 (138) | 1963 (137) |
| R-work | 0.2893 (0.4323) | 0.1785 (0.3437) |
| R-free | 0.2994 (0.4269) | 0.1968 (0.3910) |
| Number of non-hydrogen atoms | 3070 | 3070 |
| Macromolecules | 3070 | 3070 |
| Ligands | 5 | 1 |
| Solvent | 0 | 0 |
| Protein residues | 215 | 216 |
| RMS (bonds) (Å) | 0.16 | 0.42 |
| RMS (angles) (°) | 0.43 | 0.73 |
| Ramachandran favored (%) | 98 | 98 |
| Ramachandran allowed (%) | 2 | 2 |
| Ramachandran outliers (%) | 0 | 0 |
| Rotamer outliers (%) | 0 | 0 |
| Clashscore | 5 | 7 |
| Average B-factor | 98 | 165 |

**Table S16. X-ray diffraction statistics for CC1$^{+10}$ PDB entries 9YZD and 9YZE**

| PDB code | 9YZD | 9YZE |
|---|---|---|
| **Data collection** | | |
| Light source | Synchrotron | Synchrotron |
| Wavelength (Å) | 1 | 1 |
| Resolution Range (Å) | 45.29-3.05 (3.16-3.05) | 45.86-4.07 (4.28-4.07) |
| Space group | I121 | I121 |
| **Unit cell dimensions** | | |
| a, b, c (Å) | 76.03, 124.86, 139.09 | 76.03, 124.86, 139.09 |
| $\alpha$, β, ɣ (°) | 90.00, 91.85, 90.00 | 90.00, 91.85, 90.00 |
| Unique reflections | 23118 (1365) | 26992 (3349) |
| Multiplicity | 3.4 (3.5) | 3.2 (2.9) |
| Completeness (%) | 96.91 (99.4) | 92.5 (78.7) |
| Mean I/sigma(I) | 7.4 (1.1) | 2.2 (1.0) |
| Wilson B-factor | 92.15 | 95.14 |
| R-merge | 0.099 (0.913) | 0.796 (0.731) |
| R-meas | 0.118 (1.080) | 0.945 (0.886) |
| R-pim | 0.063 (0.572) | 0.526 (0.493) |
| CC1/2 | 0.973 (0.779) | 0.954(0.373) |
| **Refinement** | | |
| Reflections used in refinement | 23116 (1236) | 9975 (1135) |
| Reflections used for Rfree | 2278 (129) | 987 (122) |
| R-work | 0.2535 (0.4228) | 0.2470 (0.3735) |
| R-free | 0.2645 (0.4521) | 0.2757 (0.4248) |
| Number of non-hydrogen atoms | 3040 | 2981 |
| Macromolecules | 3040 | 2981 |
| Ligands | 1 | 1 |
| Solvent | 0 | 0 |
| Protein residues | 216 | 217 |
| RMS (bonds) (Å) | 0.23 | 0.18 |
| RMS (angles) (°) | 0.47 | 0.45 |
| Ramachandran favored (%) | 98 | 98 |
| Ramachandran allowed (%) | 2 | 2 |
| Ramachandran outliers (%) | 0 | 0 |
| Rotamer outliers (%) | 0 | 0 |
| Clashscore | 6 | 4 |
| Average B-factor | 163 | 131 |

**Table S17. X-ray diffraction statistics for CC1$^{+10}$ PDB entries 9YZF and 9YZG**

| PDB code | 9YZF | 9YZG |
|---|---|---|
| **Data collection** | | |
| Light source | Synchrotron | Synchrotron |
| Wavelength (Å) | 1 | 1 |
| Resolution Range (Å) | 46.22-3.07 (3.13-3.07) | 46.33-3.09 (3.16-3.09) |
| Space group | I121 | I121 |
| **Unit cell dimensions** | | |
| a, b, c (Å) | 74.55, 121.56, 142.89 | 74.17, 127.04, 131.72 |
| $\alpha$, β, ɣ (°) | 90.00, 95.17, 90.00 | 90.00, 91.20, 90.00 |
| Unique reflections | 23419 (4252) | 22096 (3992) |
| Multiplicity | 3.5 (3.7) | 3.5 (3.6) |
| Completeness (%) | 98.5 (98.7) | 98.7 (99.0) |
| Mean I/sigma(I) | 6.6 (1.3) | 7.7 (1.0) |
| Wilson B-factor | 90.27 | 97.8 |
| R-merge | 0.101 (0.900) | 0.096 (1.394) |
| R-meas | 0.120 (1.055) | 0.114 (1.639) |
| R-pim | 0.064 (0.547) | 0.061 (0.854) |
| CC1/2 | 0.997 (0.853) | 0.993 (0.669) |
| **Refinement** | | |
| Reflections used in refinement | 23302 (1267) | 21971 (1209) |
| Reflections used for Rfree | 2314 (138) | 2190 (134) |
| R-work | 0.2323 (0.5056) | 0.2736 (0.4703) |
| R-free | 0.2408 (0.5519) | 0.3007 (0.4898) |
| Number of non-hydrogen atoms | 3099 | 3059 |
| Macromolecules | 3099 | 3059 |
| Ligands | 1 | 2 |
| Solvent | 0 | 0 |
| Protein residues | 219 | 217 |
| RMS (bonds) (Å) | 0.19 | 0.16 |
| RMS (angles) (°) | 0.43 | 0.42 |
| Ramachandran favored (%) | 97 | 97 |
| Ramachandran allowed (%) | 2 | 2 |
| Ramachandran outliers (%) | 1 | 1 |
| Rotamer outliers (%) | 0 | 0 |
| Clashscore | 5 | 6 |
| Average B-factor | 150 | 159 |

**Table S18. X-ray diffraction statistics for CC1$^{+10}$ PDB entries 9YZI and 9YZJ**

| PDB code | 9YZI | 9YZJ |
|---|---|---|
| **Data collection** | | |
| Light source | Synchrotron | Synchrotron |
| Wavelength (Å) | 1 | 1 |
| Resolution Range (Å) | 46.70-4.00 (4.21-4.00) | 46.01-2.97 (3.16-2.97) |
| Space group | I121 | I121 |
| **Unit cell dimensions** | | |
| a, b, c (Å) | 74.90, 121.62, 138.04 | 73.78, 132.65, 126.77 |
| $\alpha$, β, ɣ (°) | 99.00, 90.41, 90.00 | 90.00, 90.26, 90.00 |
| Unique reflections | 10367 (2874) | 24810 (4023) |
| Multiplicity | 4.0 (3.3) | 3.5 (3.5) |
| Completeness (%) | 98.4 (96.6) | 98.4 (99) |
| Mean I/sigma(I) | 3.2 (0.9) | 11.4 (1.0) |
| Wilson B-factor | 151.42 | 99.16 |
| R-merge | 0.342 (1.551) | 0.065 (1.102) |
| R-meas | 0.403 (1.982) | 0.078 (1.301) |
| R-pim | 0.211 (1.067) | 0.042 (0.684) |
| CC1/2 | 0.958 (0.291) | 0.998 (0.798) |
| **Refinement** | | |
| Reflections used in refinement | 10313 (1291) | 21738 (1199) |
| Reflections used for Rfree | 1022 (144) | 2172 (126) |
| R-work | 0.2961 (0.3818) | 0.2721 (0.4371) |
| R-free | 0.3216 (0.4159) | 0.2917 (0.5107) |
| Number of non-hydrogen atoms | 2853 | 2896 |
| Macromolecules | 2853 | 2896 |
| Ligands | 4 | 4 |
| Solvent | 0 | 0 |
| Protein residues | 203 | 221 |
| RMS (bonds) (Å) | 0.26 | 0.45 |
| RMS (angles) (°) | 0.51 | 0.71 |
| Ramachandran favored (%) | 97 | 96 |
| Ramachandran allowed (%) | 3 | 3 |
| Ramachandran outliers (%) | 0 | 1 |
| Rotamer outliers (%) | 0 | 3 |
| Clashscore | 8 | 17 |
| Average B-factor | 224 | 156 |

**Table S19. X-ray diffraction statistics for CC1$^{+10}$ PDB entry 9Z55**

| | |
|---|---|
| PDB code | 9Z55 |
| **Data collection** | |
| Light source | Synchrotron |
| Wavelength (Å) | 1 |
| Resolution Range (Å) | 64.94-3.94 (4.12-3.94) |
| Space group | C121 |
| **Unit cell dimensions** | |
| a, b, c (Å) | 146.48, 129.88, 73.32 |
| $\alpha$, β, ɣ (°) | 90.00, 118.58, 90.00 |
| Unique reflections | 10668 (3014) |
| Multiplicity | 7.0 (7.1) |
| Completeness (%) | 99.5 (99.9) |
| Mean I/sigma(I) | 8.8 (2.1) |
| Wilson B-factor | 189.68 |
| R-merge | 0.095 (0.804) |
| R-meas | 0.103 (0.867) |
| R-pim | 0.039 (0.322) |
| CC1/2 | 0.998 (0.926) |
| **Refinement** | |
| Reflections used in refinement | 10599 (1189) |
| Reflections used for Rfree | 1058 (127) |
| R-work | 0.2968 (0.4202) |
| R-free | 0.3161 (0.4580) |
| Number of non-hydrogen atoms | 2862 |
| Macromolecules | 2862 |
| Ligands | 0 |
| Solvent | 0 |
| Protein residues | 216 |
| RMS (bonds) (Å) | 0.25 |
| RMS (angles) (°) | 0.49 |
| Ramachandran favored (%) | 96 |
| Ramachandran allowed (%) | 4 |
| Ramachandran outliers (%) | 0 |
| Rotamer outliers (%) | 0 |
| Clashscore | 6 |
| Average B-factor | 299 |

**Table S20. X-ray diffraction statistics for CC1+21 PDB entry 9YZK**

| PDB code | 9YZK |
|---|---|
| **Data collection** | |
| Light source | Synchrotron |
| Wavelength (Å) | 1 |
| Resolution Range (Å) | 47.53-5.10 (5.37-5.10) |
| Space group | I121 |
| **Unit cell dimensions** | |
| a, b, c (Å) | 75.84, 163.02, 192.28 |
| $\alpha$, β, ɣ (°) | 90.00, 98.61, 90.00 |
| Unique reflections | 14099 (3944) |
| Multiplicity | 3.4 (3.1) |
| Completeness (%) | 98.4 (97.7) |
| Mean I/sigma(I) | 4.1 (0.2) |
| Wilson B-factor | 304.94 |
| R-merge | 0.119 (5.581) |
| R-meas | 0.142 (6.708) |
| R-pim | 0.075 (3.673) |
| CC1/2 | 0.997 (0.196) |
| **Refinement** | |
| Reflections used in refinement | 9121 (1050) |
| Reflections used for Rfree | 910 (117) |
| R-work | 0.2763 (0.5614) |
| R-free | 0.3040 (0.6073) |
| Number of non-hydrogen atoms | 3367 |
| Macromolecules | 3367 |
| Ligands | 1 |
| Solvent | 0 |
| Protein residues | 224 |
| RMS (bonds) (Å) | 0.22 |
| RMS (angles) (°) | 0.52 |
| Ramachandran favored (%) | 97 |
| Ramachandran allowed (%) | 3 |
| Ramachandran outliers (%) | 0 |
| Rotamer outliers (%) | 0 |
| Clashscore | 6 |
| Average B-factor | 383 |

**Table S21. X-ray diffraction statistics for $CC1^{+10}$ PDB entries 9Z1B and 9Z1C**

| **PDB code** | **9Z1B** | **9Z1C** |
|---|---|---|
| **Data collection** | | |
| Light source | Synchrotron | Synchrotron |
| Wavelength (Å) | 1 | 1 |
| Resolution Range (Å) | 42.58-3.88 (4.06-3.88) | 45.27-3.04 (3.11-3.04) |
| Space group | I121 | I121 |
| **Unit cell dimensions** | | |
| a, b, c (Å) | 72.98, 126.16, 135.73 | 72.72, 129.49, 131.28 |
| $\alpha$, β, ɣ (°) | 90.00, 91.06, 90.00 | 90.00, 90.95, 90.00 |
| Unique reflections | 22544 (4078) | 23207 (4199) |
| Multiplicity | 3.5 (3.6) | 3.5 (3.6) |
| Completeness (%) | 98.8 (99.0) | 99.0 (99.4) |
| Mean I/sigma(I) | 7.8 (1.6) | 10.3 (1.3) |
| Wilson B-factor | 121.72 | 92.33 |
| R-merge | 0.118 (0.747) | 0.085 (1.153) |
| R-meas | 0.139 (0.876) | 0.100 (1.355) |
| R-pim | 0.073 (0.455) | 0.053 (0.706) |
| CC1/2 | 0.998 (0.878) | 1.0 (0.833) |
| **Refinement** | | |
| Reflections used in refinement | 10987 (1257) | 23075 (1310) |
| Reflections used for Rfree | 1111 (146) | 2298 (142) |
| R-work | 0.2743 (0.3454) | 0.3000 (0.5731) |
| R-free | 0.3081 (0.3631) | 0.3251 (0.5991) |
| Number of non-hydrogen atoms | 2905 | 2916 |
| Macromolecules | 2905 | 2916 |
| Ligands | 11 | 4 |
| Solvent | 0 | 0 |
| Protein residues | 216 | 216 |
| RMS (bonds) (Å) | 0.18 | 0.15 |
| RMS (angles) (°) | 0.4 | 0.38 |
| Ramachandran favored (%) | 97 | 98 |
| Ramachandran allowed (%) | 2 | 2 |
| Ramachandran outliers (%) | 1 | 0 |
| Rotamer outliers (%) | 0 | 0 |
| Clashscore | 6 | 5 |
| Average B-factor | 200 | 122 |

**Table S22. X-ray diffraction statistics for CC1$^{+10}$ PDB entry 9Z1D**

| PDB code | 9Z1D |
|---|---|
| **Data collection** | |
| Light source | Synchrotron |
| Wavelength (Å) | 1 |
| Resolution Range (Å) | 39.72-3.06 (3.13-3.06) |
| Space group | I121 |
| **Unit cell dimensions** | |
| a, b, c (Å) | 72.38, 124.54, 136.54 |
| $\alpha$, β, ɣ (°) | 90.00, 90.17, 90.00 |
| Unique reflections | 22756 (4075) |
| Multiplicity | 3.6 (3.6) |
| Completeness (%) | 99.4 (98.9) |
| Mean I/sigma(I) | 9.1 (0.9) |
| Wilson B-factor | 113.12 |
| R-merge | 0.129 (1.300) |
| R-meas | 0.153 (1.523) |
| R-pim | 0.081 (0.789) |
| CC1/2 | 0.724 (0.869) |
| **Refinement** | |
| Reflections used in refinement | 22199 (1001) |
| Reflections used for Rfree | 2227 (112) |
| R-work | 0.2929 (0.7702) |
| R-free | 0.3184 (0.9681) |
| Number of non-hydrogen atoms | 2848 |
| Macromolecules | 2848 |
| Ligands | 1 |
| Solvent | 0 |
| Protein residues | 216 |
| RMS (bonds) (Å) | 0.17 |
| RMS (angles) (°) | 0.43 |
| Ramachandran favored (%) | 98 |
| Ramachandran allowed (%) | 2 |
| Ramachandran outliers (%) | 0 |
| Rotamer outliers (%) | 0 |
| Clashscore | 4 |
| Average B-factor | 153 |

**Table S23. X-ray diffraction statistics for CC1$^{+10}$ PDB entries 9YZL and 9Z4E**

| PDB code | 9ZYL | 9Z4E |
|---|---|---|
| **Data collection** | | |
| Light source | Synchrotron | Synchrotron |
| Wavelength (Å) | 1 | 1 |
| Resolution Range (Å) | 48.31-3.02 (3.19-3.02) | 65.07-3.63 (3.78-3.63) |
| Space group | I 1 2 1 | C 1 2 1 |
| **Unit cell dimensions** | | |
| a, b, c (Å) | 73.36, 120.29, 134.40 | 159.04, 115.33, 72.72 |
| $\alpha$, β, ɣ (°) | 90.00, 91.47, 90.00 | 90.00, 116.50, 90.00 |
| Unique reflections | 22518 (3649) | 13262 (3166) |
| Multiplicity | 3.5 (3.6) | 7.1 (7.2) |
| Completeness (%) | 98.3 (98.5) | 99.2 (99.6) |
| Mean I/sigma(I) | 3.8 (0.9) | 10 (1.9) |
| Wilson B-factor | 86.65 | 149.49 |
| R-merge | 0.198 (1.428) | 0.104 (0.886) |
| R-meas | 0.234 (1.675) | 0.113 (0.955) |
| R-pim | 0.124 (0.868) | 0.042 (0.352) |
| CC1/2 | 0.966 (0.622) | 0.999 (0.902) |
| **Refinement** | | |
| Reflections used in refinement | 22417 (1243) | 13202 (1303) |
| Reflections used for Rfree | 2223 (137) | 1319 (153) |
| R-work | 0.2972 (0.4570) | 0.2473 (0.3725) |
| R-free | 0.3056 (0.4581) | 0.2853 (0.4225) |
| Number of non-hydrogen atoms | 3490 | 3172 |
| Macromolecules | 3490 | 3172 |
| Ligands | 2 | 3 |
| Solvent | 0 | 0 |
| Protein residues | 280 | 266 |
| RMS (bonds) (Å) | 0.18 | 0.20 |
| RMS (angles) (°) | 0.44 | 0.41 |
| Ramachandran favored (%) | 97 | 98 |
| Ramachandran allowed (%) | 3 | 2 |
| Ramachandran outliers (%) | 0 | 0 |
| Rotamer outliers (%) | 0 | 0 |
| Clashscore | 6 | 9 |
| Average B-factor | 139 | 214 |

**Table S24. X-ray diffraction statistics for $CC1^{+10}$ PDB entries 9YZM and 9YZN**

| PDB code | 9ZYM | 9ZYN |
|---|---|---|
| **Data collection** | | |
| Light source | Synchrotron | Synchrotron |
| Wavelength (Å) | 1 | 1 |
| Resolution Range (Å) | 44.48-3.07 (3.14-3.07) | 46.80-3.16 (3.23-3.16) |
| Space group | I 1 2 1 | I 1 2 1 |
| **Unit cell dimensions** | | |
| a, b, c (Å) | 73.78, 125.19, 131.23 | 76.03, 124.86, 139.09 |
| $\alpha$, β, γ (°) | 90.00, 92.26, 90.00 | 90.00, 91.85, 90.00 |
| Unique reflections | 22050 (4010) | 23358 (4253) |
| Multiplicity | 3.5 (3.7) | 3.5 (3.6) |
| Completeness (%) | 98.9 (99.3) | 98.7 (99.4) |
| Mean I/sigma(I) | 7.6 (1.3) | 10.5 (1.9) |
| Wilson B-factor | 87.54 | 93.44 |
| R-merge | 0.123 (1.054) | 0.077 (0.819) |
| R-meas | 0.145 (1.237) | 0.092 (0.962) |
| R-pim | 0.077 (0.643) | 0.049 (0.501) |
| CC1/2 | 0.998 (0.741) | 0.999 (0.863) |
| **Refinement** | | |
| Reflections used in refinement | 21973 (1229) | 21733 (1189) |
| Reflections used for Rfree | 2194 (131) | 2166 (136) |
| R-work | 0.2594 (0.4232) | 0.2618 (0.3895) |
| R-free | 0.2691 (0.4468) | 0.2827 (0.4011) |
| Number of non-hydrogen atoms | 3733 | 3699 |
| Macromolecules | 3733 | 3699 |
| Ligands | 2 | 1 |
| Solvent | 0 | 0 |
| Protein residues | 325 | 329 |
| RMS (bonds) (Å) | 0.16 | 0.14 |
| RMS (angles) (°) | 0.37 | 0.35 |
| Ramachandran favored (%) | 96 | 97 |
| Ramachandran allowed (%) | 3 | 2 |
| Ramachandran outliers (%) | 1 | 1 |
| Rotamer outliers (%) | 0 | 0 |
| Clashscore | 4 | 3 |
| Average B-factor | 139 | 117 |

**Table S25. X-ray diffraction statistics for CC1$^{+10}$ PDB entries 9YZO and 9YZP**

| PDB code | 9YZO | 9YZP |
|---|---|---|
| **Data collection** | | |
| Light source | Synchrotron | Synchrotron |
| Wavelength (Å) | 1 | 1 |
| Resolution Range (Å) | 46.68-2.96 (3.02-2.96) | 47.16-3.67 (3.80-3.67) |
| Space group | I 1 2 1 | I 1 2 1 |
| **Unit cell dimensions** | | |
| a, b, c (Å) | 74.11, 118.26, 141.68 | 73.92, 126.61, 137.89 |
| $\alpha$, β, ɣ (°) | 90.00, 90.89, 90.00 | 90.00, 90.92, 90.00 |
| Unique reflections | 24985 (4020) | 13732 (3297) |
| Multiplicity | 3.5 (3.6) | 3.3 (3.4) |
| Completeness (%) | 97.8 (98.2) | 98.4 (99.3) |
| Mean I/sigma(I) | 7.4 (0.9) | 2.8 (0.7) |
| Wilson B-factor | 90.27 | 157.05 |
| R-merge | 0.098 (1.102) | 0.202 (1.619) |
| R-meas | 0.116 (1.293) | 0.243 (1.923) |
| R-pim | 0.062 (0.671) | 0.132 (1.026) |
| CC1/2 | 0.997 (0.663) | 0.997 (0.368) |
| **Refinement** | | |
| Reflections used in refinement | 24816 (1216) | 13695 (1228) |
| Reflections used for Rfree | 2467 (140) | 1356 (127) |
| R-work | 0.2898 (0.4673) | 0.2720 (0.4252) |
| R-free | 0.3121 (0.5115) | 0.2949 (0.4167) |
| Number of non-hydrogen atoms | 3650 | 3177 |
| Macromolecules | 3650 | 3177 |
| Ligands | 1 | 1 |
| Solvent | 0 | 0 |
| Protein residues | 329 | 272 |
| RMS (bonds) (Å) | 0.22 | 0.45 |
| RMS (angles) (°) | 0.50 | 0.69 |
| Ramachandran favored (%) | 96 | 98 |
| Ramachandran allowed (%) | 3 | 2 |
| Ramachandran outliers (%) | 1 | 0 |
| Rotamer outliers (%) | 1 | 0 |
| Clashscore | 10 | 10 |
| Average B-factor | 104 | 226 |

**Table S26. X-ray diffraction statistics for $CC1^{+10}$ PDB entries 9YZQ and 9YZR**

| PDB code | 9YZQ | 9YZR |
|---|---|---|
| **Data collection** | | |
| Light source | Synchrotron | Synchrotron |
| Wavelength (Å) | 1 | 1 |
| Resolution Range (Å) | 46.92-3.75 (3.90-3.75) | 45.29-3.39 (3.48-3.39) |
| Space group | I 1 2 1 | I 1 2 1 |
| **Unit cell dimensions** | | |
| a, b, c (Å) | 74.12, 122.40, 140.44 | 76.03, 124.86, 139.09 |
| $\alpha$, β, ɣ (°) | 90.00, 90.38, 90.00 | 90.00, 91.85, 90.00 |
| Unique reflections | 15454 (3568) | 21916 (3999) |
| Multiplicity | 3.4 (3.4) | 3.5 (3.6) |
| Completeness (%) | 97.5 (95.1) | 98.8 (99.3) |
| Mean I/sigma(I) | 4.1 (0.4) | 6.5 (1.1) |
| Wilson B-factor | 135.03 | 92.55 |
| R-merge | 0.159 (4.541) | 0.134 (1.332) |
| R-meas | 0.190 (5.424) | 0.159 (1.566) |
| R-pim | 0.102 (2.925) | 0.085 (0.816) |
| CC1/2 | 0.998 (0.436) | 0.998 (0.658) |
| **Refinement** | | |
| Reflections used in refinement | 12659 (1231) | 17764 (1200) |
| Reflections used for Rfree | 1263 (144) | 1767 (127) |
| R-work | 0.2951 (0.4783) | 0.2652 (0.4375) |
| R-free | 0.3382 (0.4722) | 0.2976 (0.4531) |
| Number of non-hydrogen atoms | 3198 | 3409 |
| Macromolecules | 3198 | 3409 |
| Ligands | 2 | 1 |
| Solvent | 0 | 0 |
| Protein residues | 272 | 298 |
| RMS (bonds) (Å) | 0.38 | 0.14 |
| RMS (angles) (°) | 0.63 | 0.37 |
| Ramachandran favored (%) | 97 | 97 |
| Ramachandran allowed (%) | 2 | 3 |
| Ramachandran outliers (%) | 1 | 0 |
| Rotamer outliers (%) | 0 | 0 |
| Clashscore | 7 | 5 |
| Average B-factor | 221 | 146 |

**Table S27. X-ray diffraction statistics for CC1$^{+10}$ PDB entries 9YZS and 9YZT**

| PDB code | 9YZS | 9YZT |
|---|---|---|
| **Data collection** | | |
| Light source | Synchrotron | Synchrotron |
| Wavelength (Å) | 1 | 1 |
| Resolution Range (Å) | 46.92-3.08 (3.15-3.08) | 46.13-3.30 (3.39-3.30) |
| Space group | I 1 2 1 | I 1 2 1 |
| **Unit cell dimensions** | | |
| a, b, c (Å) | 74.19, 122.96, 135.18 | 72.75, 126.04, 134.95 |
| $\alpha$, β, ɣ (°) | 90.00, 91.86, 90.00 | 90.00, 90.24, 90.00 |
| Unique reflections | 22226 (4000) | 22292 (4017) |
| Multiplicity | 3.5 (3.6) | 3.5 (3.6) |
| Completeness (%) | 98.9 (98.9) | 98.8 (98.5) |
| Mean I/sigma(I) | 6.7 (1.3) | 6.1 (1.6) |
| Wilson B-factor | 92.55 | 118.28 |
| R-merge | 0.099 (1.101) | 0.186 (1.488) |
| R-meas | 0.143 (1.293) | 0.220 (1.754) |
| R-pim | 0.076 (0.673) | 0.115 (0.919) |
| CC1/2 | 0.998 (0.777) | 0.864 (0.730) |
| **Refinement** | | |
| Reflections used in refinement | 22106 (1250) | 17991 (1183) |
| Reflections used for Rfree | 1250 (142) | 1798 (144) |
| R-work | 0.2728 (0.4119) | 0.2827 (0.4684) |
| R-free | 0.2906 (0.4524) | 0.2921 (0.4972) |
| Number of non-hydrogen atoms | 3815 | 3624 |
| Macromolecules | 3815 | 3624 |
| Ligands | 2 | 1 |
| Solvent | 0 | 0 |
| Protein residues | 329 | 315 |
| RMS (bonds) (Å) | 0.21 | 0.14 |
| RMS (angles) (°) | 0.41 | 0.37 |
| Ramachandran favored (%) | 95 | 97 |
| Ramachandran allowed (%) | 4 | 3 |
| Ramachandran outliers (%) | 1 | 0 |
| Rotamer outliers (%) | 0 | 0 |
| Clashscore | 6 | 6 |
| Average B-factor | 161 | 149 |

**Table S28. X-ray diffraction statistics for CC1[+10] PDB entries 9YZU and 9Z44**

| PDB code | 9YZU | 9Z44 |
|---|---|---|
| **Data collection** | | |
| Light source | Synchrotron | Synchrotron |
| Wavelength (Å) | 1 | 1 |
| Resolution Range (Å) | 63.71-3.05 (3.12-3.05) | 47.99-7.20 |
| Space group | C 1 2 1 | I 1 2 1 |
| **Unit cell dimensions** | | |
| a, b, c (Å) | 152.28, 127.19, 72.98 | 73.49, 127.65, 141.18 |
| $\alpha$, β, ɣ (°) | 90.00, 117.60, 90.00 | 90.00, 91.98, 90.00 |
| Unique reflections | 23188 (4136) | 1895 |
| Multiplicity | 7.0 (6.6) | 1.0 |
| Completeness (%) | 98.6 (97.6) | 99.14 |
| Mean I/sigma(I) | 11.3 (2.0) | 2.12 |
| Wilson B-factor | 85.24 | 312.80 |
| R-merge | 0.130 (1.060) | 0.688 |
| R-meas | 0.140 (1.151) | 0.818 |
| R-pim | 0.053 (0.443) | 0.437 |
| CC1/2 | 0.999 (0.891) | 0.301 |
| **Refinement** | | |
| Reflections used in refinement | 23071 (1353) | 1895 |
| Reflections used for Rfree | 2235 (118) | 192 |
| R-work | 0.2247 (0.3769) | 0.2804 |
| R-free | 0.257 (0.3390) | 0.3453 |
| Number of non-hydrogen atoms | 3445 | 2821 |
| Macromolecules | 3445 | 2821 |
| Ligands | 2 | 0 |
| Solvent | 0 | 0 |
| Protein residues | 278 | 237 |
| RMS (bonds) (Å) | 0.24 | 0.16 |
| RMS (angles) (°) | 0.46 | 0.42 |
| Ramachandran favored (%) | 98 | 96 |
| Ramachandran allowed (%) | 2 | 3 |
| Ramachandran outliers (%) | 0 | 1 |
| Rotamer outliers (%) | 0 | 0 |
| Clashscore | 5 | 4 |
| Average B-factor | 124 | 309 |
